# Light-cycle time-restricted feeding remodels a hidden layer of the cardiac transcriptome through sex-specific transcript switching

**DOI:** 10.64898/2026.09.22.753546

**Authors:** Shrishti Naidu, Abhilash Prabhat, Bailey Peck, Tanya Seward, Elizabeth S. Schroder, Brian P. Delisle, Yuan Wen

## Abstract

Light-cycle time-restricted feeding disrupts daily cardiovascular and thermoregulatory rhythms, but the molecular effects of light-cycle time-restricted feeding on the heart have been measured only at the level of total gene expression. We used Oxford Nanopore long-read RNA sequencing to resolve the full-length ventricular transcriptome from male and female mice under ad libitum feeding or light-cycle time-restricted feeding across the 24-hour cycle. Greater than 20% of cardiac transcripts represent unannotated variants of known genes absent from the current GENCODE reference annotation. Light-cycle time-restricted feeding reorganizes transcript usage across hundreds of genes, including genes encoding splicing regulators, largely without changing total gene expression. The genes affected are sex-specific, with fewer than 2% of changes shared at the gene, transcript, and transcript-usage levels. We show that transcript-level regulation is a previously underrecognized component of the cardiac response to altered feeding behavior, undetected by conventional short-read approaches.

**HIGHLIGHTS:**

- Feeding behavior remodels the cardiac transcriptome at gene and transcript levels
- >20% of cardiac transcripts identified are absent from current reference annotations
- Transcript switching occurs without changes in total gene expression
- Feeding elicits sex-specific responses across gene, transcript, and transcript usage

## INTRODUCTION

Feeding at the wrong time of day disrupts cardiac physiology. In humans, irregular meal timing and late-night eating are associated with a greater risk of obesity, type 2 diabetes, and cardiovascular disease^1–5^. In nocturnal mice, light-cycle time-restricted feeding (LRF) shifts physiological rhythms in heart rate, ventricular repolarization (QT intervals), mean arterial pressure, and core body temperature, and exacerbates arrhythmia phenotypes in long QT syndrome models^6–8^. At the molecular level, LRF affects cellular circadian clocks, which are ubiquitous transcriptional-translational feedback loops that generate ∼24-hour rhythms in gene expression and cellular function, enabling organisms to anticipate predictable environmental changes^4,9^. The central circadian clock, located in the hypothalamic suprachiasmatic nucleus (SCN), is entrained by light-dark cycles. The SCN synchronizes peripheral clocks through neurohumoral signaling and by regulating the timing of feeding behavior. LRF causes misalignment between the central clock and peripheral clocks, including those in the heart^10–12^. LRF uncouples peripheral tissue clocks from the SCN without altering SCN gene expression rhythms^11^, phase-shifting and dampening clock gene expression in the heart^10^. Yet the molecular programs mediating these cardiac effects have been characterized predominantly by quantitative PCR (qPCR) and short-read RNA sequencing, methods limited to quantifying total gene expression^10–12^.

Genes produce multiple transcripts through alternative splicing, and their relative proportions can shift without changes in total gene expression, a post-transcriptional regulatory layer with direct consequences for gene function^13^. For some major cardiac genes, different transcripts or splice variants determine cardiac tissue function. For example, titin’s two major cardiac splice isoforms differ in stiffness^14^. N2B, the shorter isoform, confers greater myofilament stiffness than N2BA, the longer isoform. Their relative proportions vary across species with different heart rates^14^. The disease relevance of this post-transcriptional regulatory layer is also well established. In rats, a loss-of-function mutation in *RBM20* disrupts exon inclusion in titin, *CaMKIIδ*, *LDB3*, and *CACNA1C*, producing dilated cardiomyopathy with fibrosis and arrhythmia^15^. Humans carrying *RBM20* mutations show a similar phenotype and share conserved *RBM20*-dependent splicing changes in these same genes^15^. Here, we test whether LRF, a physiological behavioral change rather than a genetic mutation, is sufficient to reorganize the full-length cardiac transcriptome and splicing landscape.

Male and female hearts show extensive baseline differences in gene expression patterns^16^. We hypothesized that, despite this sex divergence, LRF would elicit a conserved subset of differentially expressed genes (DEGs) and differentially expressed transcripts (DETs) between sexes. To isolate these steady-state transcriptional and post-transcriptional responses to LRF independent of time-of-day effects, we collected ventricular tissue across four zeitgeber time points over the 24-hour cycle. Short-read sequencing fragments transcripts and must computationally infer which exon combinations were present in each original molecule, introducing statistical uncertainty that grows with transcript isoform complexity^17,18^. Here, we apply Oxford Nanopore long-read RNA sequencing to ventricular tissue from male and female mice under ad libitum feeding (ALF), during which nocturnal mice naturally consume most of their food during the dark cycle, or under LRF, which restricts feeding to the light cycle. Because each read represents a single full-length molecule, long-read sequencing resolves identity directly, enabling gene-level, transcript-level, and transcript usage analyses within the same experiment^19–22^.

We identified >20% of cardiac transcripts as unannotated variants of known genes. LRF altered temporal regulation of gene expression. 68–74% of DETs and 91–95% of differential transcript usage (DTU) changes were independent of total gene expression changes. LRF elicited largely sex-specific cardiac transcriptional responses, with fewer than 2% of changes shared between sexes across gene expression, transcript abundance, and transcript usage. Genes with DTU under LRF were enriched for multiple directional post-transcriptional splicing events in both sexes. These findings show that a behavior can induce DET changes not captured by standard RNA sequencing, with implications for disease studies that rely on DETs, since a subset of DETs appears sensitive to behavioral change in a sex-specific manner.

## RESULTS

### LRF alters cardiac clock gene expression profiles

To test the effect of LRF on the cardiac transcriptome, ventricular tissue from ALF- and LRF-fed male and female mice was collected at four zeitgeber time points (ZT; ZT1, ZT7, ZT13, and ZT19) and sequenced using Oxford Nanopore long-read sequencing (Figures 1A and 1B). Sequencing yield, read length, read-length N50, mapping rate, split-alignment fraction, and the number of transcripts detected per sample were consistent across ALF and LRF samples (Figures S1A–S1E and S1G). Gene body coverage was uniform in all samples (3′/5′ bias index 0.79–1.05) and closely matched between ALF and LRF within each sex (Figure S2), and samples clustered tightly by sex with high within-group correlation (Figure S3). Core clock genes (*Bmal1*, *Clock, Per2*, *Per3*, *Cry1*, *Cry2*), the accessory loop components *Nr1d1* and *Nr1d2*, and the clock output gene *Dbp* showed altered temporal expression profiles under LRF relative to ALF in both sexes, consistent with previously reported phase shifts in cardiac clock gene oscillations in male and female mice^10,12^ (Figure 1C).

**Figure 1.**
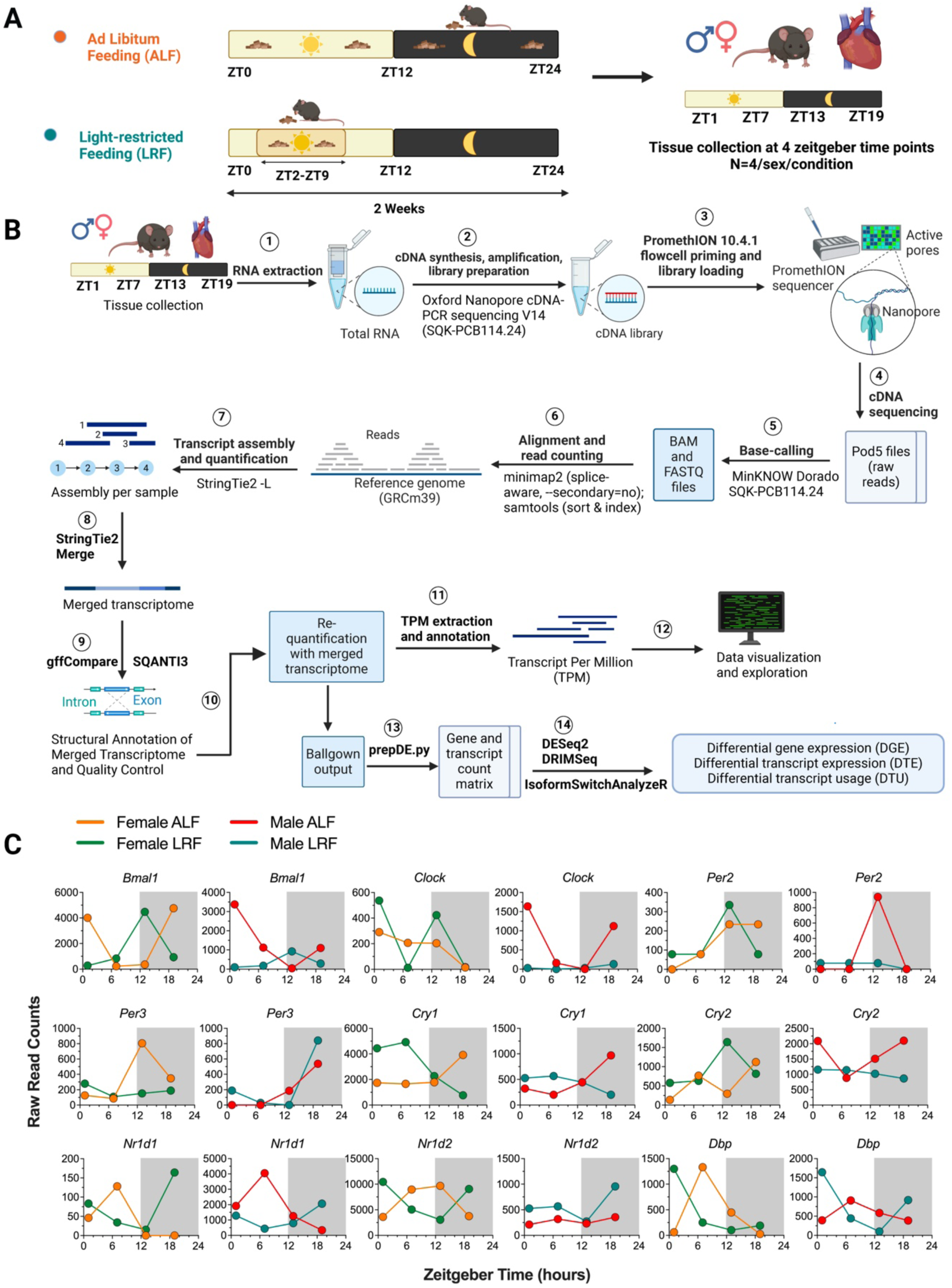
Long-read sequencing resolves the cardiac transcriptome and confirms light-cycle time-restricted feeding phase-shifts clock gene expression in both sexes. (A) Experimental design. Wild-type SV129 mice were maintained on ad libitum feeding (ALF) or light-cycle time-restricted feeding (LRF; food available at zeitgeber time [ZT] 2– 9) under a 12-hour light/12-hour dark cycle for two weeks. Ventricular tissue was collected at four ZT points (ZT1, ZT7, ZT13, and ZT19) from both sexes under each feeding condition (N = 4 per sex per condition; one biological replicate per time point; 16 samples total). (B) Long-read RNA sequencing and bioinformatic analysis pipeline. Full-length cDNA libraries were prepared using the Oxford Nanopore Technologies (ONT) cDNA-PCR Barcoding Kit V14 (SQK-PCB114-24) and sequenced on the PromethION platform with R10.4.1 flow cells. Raw signals were basecalled with Dorado, reads were aligned to GRCm39 with minimap2, and transcripts were assembled per sample using StringTie2 in long-read mode. Per-sample assemblies were merged into a unified transcriptome, structurally annotated with gffcompare, and independently evaluated for transcript quality with SQANTI3. The merged transcriptome was requantified to generate gene- and transcript-level count matrices for differential gene expression (DGE), differential transcript expression (DTE), and differential transcript usage (DTU) analyses. Numbered steps indicate workflow order. (C) Temporal expression profiles of core clock genes, the accessory loop components, and the clock output gene across the 24-hour cycle in female (left columns) and male (right columns) ventricle under ALF and LRF. Values are raw gene-level read counts. Each data point represents one biological replicate. Gray shading indicates the dark cycle (ZT12–ZT24). Orange, female ALF; green, female LRF; red, male ALF; teal, male LRF.

### One in five cardiac transcripts are absent from current annotations

Principal component analysis (PCA) of all 16 samples showed that PC1 explained 56.2% of total variance and separated male and female samples completely, while PC2 explained 4.2% (Figure 2A). Within each sex, PC1 (females 18.3%, males 19.1%) and PC2 (females 16.1%, males 15.2%) together captured variation associated with feeding condition and zeitgeber time, with PC1 showing separation by feeding condition and PC2 associated with ZT in both females and males (Figures 2B and 2C). Because male and female samples were prepared and sequenced in separate batches at different sequencing depths, within-sex analyses were used for all comparisons presented in the main figures. Comparisons between sexes were performed on depth-matched count matrices and are presented as a secondary analysis in the supplementary materials.

**Figure 2.**
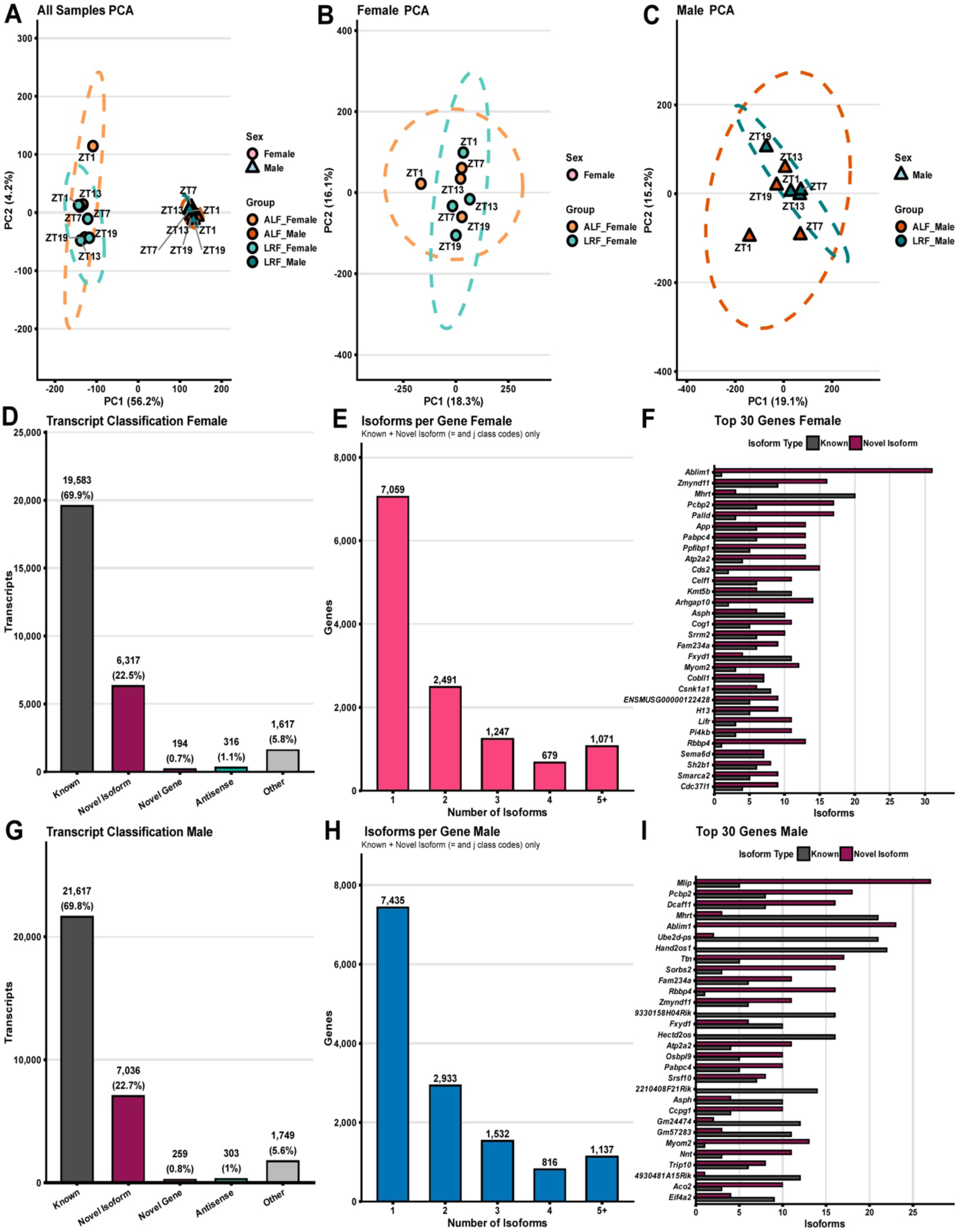
Long-read sequencing reveals a sex-stratified cardiac transcriptome with extensive unannotated transcript isoform diversity. (A) Principal component analysis (PCA) of all 16 samples based on log_2_(transcripts per million [TPM] + 1)-transformed transcript expression values (transcripts with TPM ≥ 1 in at least 2 of 16 samples; 31,621 transcripts retained). PC1 (56.2%) separated samples by sex; PC2 (4.2%) captured within-sex variation. Circles, females; triangles, males. Each point is labeled by zeitgeber time (ZT). Dashed ellipses represent 95% confidence ellipses per sex. Light orange, ad libitum feeding (ALF) females; light teal, light-cycle time-restricted feeding (LRF) females; dark orange, ALF males; dark teal, LRF males. Colors and symbols apply throughout (A)–(C). (B and C) Sex-stratified PCA in females (B; PC1 = 18.3%, PC2 = 16.1%) and males (C; PC1 = 19.1%, PC2 = 15.2%). Dashed ellipses represent 95% confidence ellipses per feeding condition group. (D and G) Number of transcripts detected (TPM ≥ 1 in at least one sample within the respective sex) by structural annotation category in females (D) and males (G), after exclusion of artifact-class transcripts (gffcompare class codes i, e, s, p). Categories: charcoal, Known; magenta, Novel Isoform; pink, Novel Gene; green, Antisense; gray, Other. Counts and percentages are shown above each bar. (E and H) Distribution of the number of transcript isoforms detected per gene in females (E) and males (H), restricted to Known and Novel Isoform classes (gffcompare class codes “=” and “j”). Genes are grouped into bins of 1, 2, 3, 4, or ≥5 transcript isoforms. (F and I) Top 30 genes ranked by total transcript isoform count in females (F) and males (I). Charcoal bars, Known isoforms; magenta bars, Novel Isoforms.

Downsampling all samples to a common depth of approximately 1.72 million reads and requantifying against the same merged transcriptome changed coverage bias by less than 1.5% in either sex (female 0.804 to 0.809; male 1.022 to 1.008) and left it unchanged between feeding conditions at both depths (Table S1), indicating that the sex difference in coverage profile reflects a stable batch characteristic rather than a depth artifact.

Transcripts were classified structurally with gffcompare against the GENCODE vM36 annotation. Across the merged transcriptome of 305,327 assembled transcripts, 90.5% matched a reference transcript exactly and 7.5% represented novel junction combinations within known genes, with all remaining classes below 1.2% (Figure S1F). Among transcripts detected per sample, 69.9% in females and 69.8% in males matched the reference annotation exactly, while 22.5% and 22.7%, respectively, were novel transcript isoforms of known genes (Figures 2D and 2G). The higher proportion among detected transcripts reflects enrichment for expressed transcript isoforms relative to the full assembled catalog. Independent classification with SQANTI3 was concordant, assigning 91.2% of catalog transcripts as full-splice matches and 8.6% as novel transcript isoforms of annotated genes (Figure S4). Approximately one in five transcripts detected in the mouse heart therefore represents a novel transcript isoform of a known gene not present in the current reference annotation.

Novel transcript isoforms were structurally distinct from annotated transcripts, with considerably longer and more exon-rich structures. Median lengths were 2,515–2,584 bp across 10–11 exons, compared with 877 bp across 3 exons for full-splice matches (Data S1). Quality-control attributes supported their integrity at both transcript ends. PolyA motifs were detected in 82.2–89.9% of novel transcript isoforms and cap analysis of gene expression (CAGE) peak support in 54.1–58.1%, both exceeding the corresponding values for full-splice matches (74.0% and 22.8%; Figure S5), and intra-priming artifacts were less frequent in novel transcript isoforms than in full-splice matches (1.3–1.8% versus 4.5%; Data S1). Most splice sites in novel transcript isoforms corresponded to annotated donor or acceptor positions (85.5–87.9%; Figure S5). Novel transcript isoforms formed by new combinations of annotated splice sites showed 97.0% canonical splice junctions, comparable to full-splice matches at 98.9% (Figure S5). Among expressed transcripts, these transcript isoforms showed short-read junction support in 83.2%, compared with 94.6% for full-splice matches (Figure S6). Transcript isoforms containing an unannotated splice site showed lower rates on both measures (5.5% canonical junctions, Figure S5; 9.2% short-read junction support, Figure S6). In this class, 90.9% paired an annotated splice donor with a novel acceptor positioned a median of 3 bp from an annotated acceptor, and 92.8% within 10 bp, indicating small shifts in acceptor position rather than use of distant alternative sites (Data S2). Because the short-read reference libraries were generated from separate animals maintained under constant darkness, junction coverage was used as supporting rather than defining evidence.

Most genes expressed a single dominant transcript isoform, with progressively fewer genes expressing two, three, four, or five or more transcript isoforms in both sexes (Figures 2E and 2H). Among the most transcript isoform-rich genes were well-characterized cardiac genes involved in sarcomere organization (*Ttn*, *Myom2*), Ca^2+^ handling (*Atp2a2*), RNA processing (*Pcbp2*, *Celf1*, *Srsf10*), mitochondrial metabolism (*Nnt*, *Aco2*), and the cardiac noncoding RNA *Mhrt* (Figures 2F and 2I). Transcript classification and transcript isoform diversity per gene were consistent across feeding conditions within each sex (Figure S7).

### Gene-level responses to LRF diverge by sex

Gene-level differential expression was assessed with DESeq2 comparing LRF to ALF within each sex, with zeitgeber time included as a covariate and differential expression defined as p < 0.01 and absolute log_2_ fold change (|log_2_FC|) ≥ 1. Most genes were expressed under both feeding conditions in each sex (76.6% in females, 80.8% in males), with 12.2% detected exclusively under ALF and 11.3% exclusively under LRF in females, and 10.6% and 8.7%, respectively, in males (Figures 3A and 3B, and S8A and S8B). Condition-specific gene sets overlapped minimally between sexes, with 137 of the ALF-specific and 97 of the LRF-specific genes shared, against 1,598–2,075 genes specific to a single sex and condition (Figures S8C–S8G).

**Figure 3.**
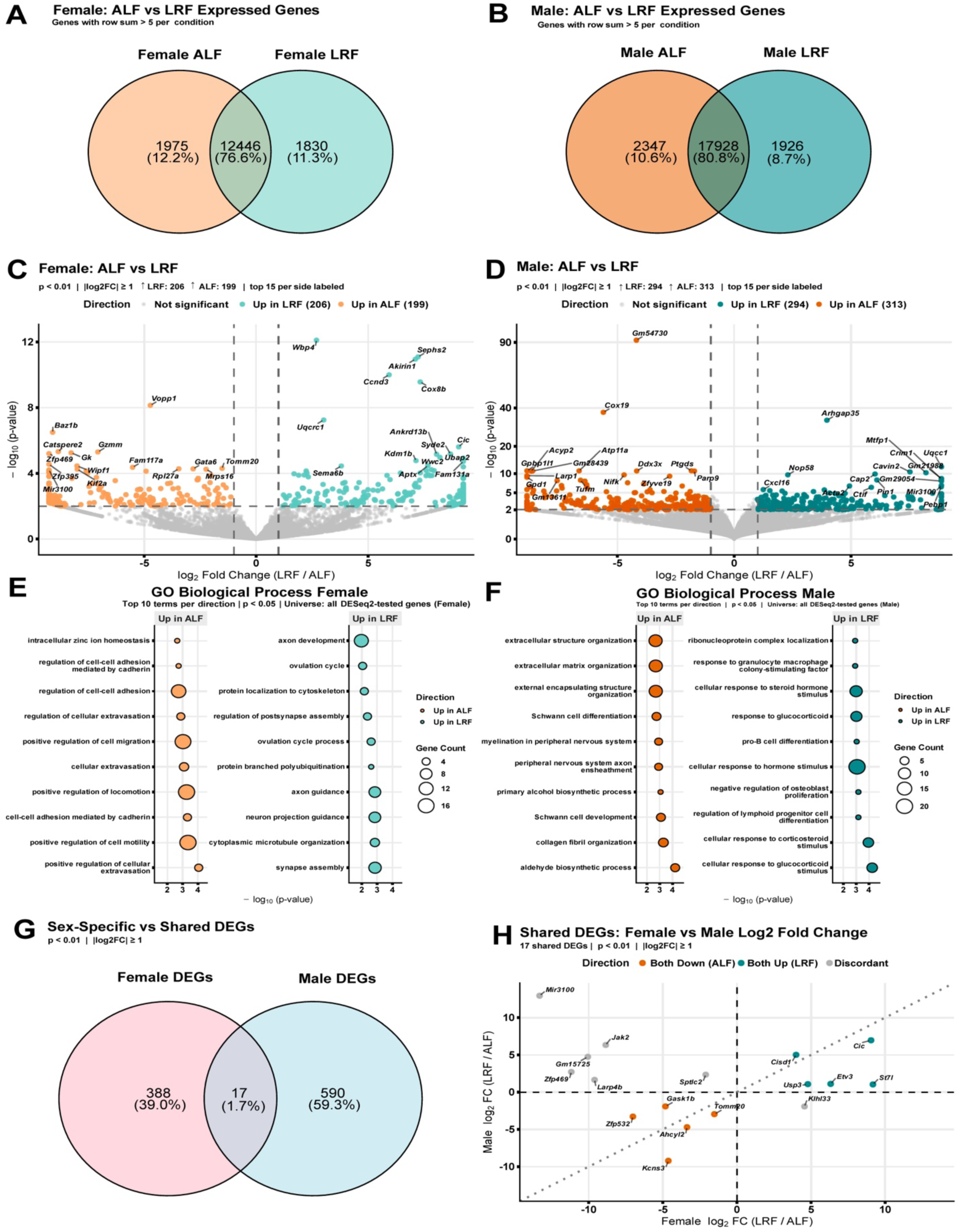
Differential cardiac gene expression under light-cycle time-restricted feeding is largely sex-specific. (A and B) Venn diagrams of expressed genes between ad libitum feeding (ALF) and light-cycle time-restricted feeding (LRF) in females (A) and males (B). A gene was considered expressed in a given sex-condition group if the sum of raw counts across all four samples exceeded 5. Counts and percentages are shown for ALF-only, LRF-only, and shared genes. Only shared genes were tested for differential expression in (C) and (D). (C and D) Volcano plots of gene-level differential expression between ALF and LRF in females (C; 206 higher under LRF, 199 higher under ALF) and males (D; 294 higher under LRF, 313 higher under ALF). The x-axis shows log_2_ fold change (log_2_FC; LRF/ALF); the y-axis shows −log_10_(p-value). Differentially expressed genes (DEGs) were defined by p < 0.01 and |log_2_FC| ≥ 1 (dashed lines). The top 15 significant DEGs per direction are labeled. Teal, higher under LRF; orange, higher under ALF; gray, not significant. (E and F) Gene Ontology (GO) biological process enrichment for DEGs in females (E) and males (F), computed separately for genes higher under ALF (left, orange) and genes higher under LRF (right, teal). The x-axis shows −log_10_(p-value); dot size is proportional to gene count. Top 10 enriched terms per direction are shown (p < 0.05). (G) Overlap of DEGs between sexes. Female-only: 388 (39.0%); male-only: 590 (59.3%); shared: 17 (1.7%). (H) Scatter plot comparing log_2_FC (LRF/ALF) of the 17 shared DEGs in females (x-axis) versus males (y-axis). All 17 genes are labeled, ranked by fold change and p-value. Teal, concordant upregulation under LRF in both sexes; orange, concordant upregulation under ALF in both sexes; gray, discordant direction between sexes. N = 4 biological replicates per sex per condition (one per zeitgeber time [ZT]).

LRF altered expression of 405 genes in females (206 higher under LRF, 199 higher under ALF) and 607 genes in males (294 higher under LRF, 313 higher under ALF; Figures 3C and 3D). Gene Ontology (GO) enrichment revealed distinct biological processes in each sex. In females, genes higher under LRF were enriched for synapse assembly, cytoplasmic microtubule organization, and axon development and guidance, along with ovulation cycle-related terms that likely reflect a small, broadly annotated gene set, while genes higher under ALF were enriched for regulation of cell-cell adhesion, cell migration, locomotion, and cell motility (Figure 3E). In males, genes higher under LRF were enriched for ribonucleoprotein complex localization and cellular response to corticosteroid and glucocorticoid stimulus, while genes higher under ALF were enriched for extracellular matrix organization and Schwann cell differentiation and development (Figure 3F).

Cross-sex comparison showed that 388 differentially expressed genes (39.0%) were detected only in females and 590 (59.3%) only in males, with 17 genes (1.7%) shared between sexes (Figure 3G). Among shared genes, a concordant subset changed in the same direction in both sexes: *Cic*, *Etv3*, *Cisd1*, *Usp3*, and *St7l* were higher under LRF, while *Gask1b*, *Zfp532*, *Ahcyl2*, *Tomm20*, and *Kcns3* were higher under ALF. The remaining shared genes, including *Mir3100*, *Jak2*, *Sptlc2*, *Klhl33*, *Zfp469*, *Larp4b*, and *Gm15725*, changed in opposite directions between sexes (Figure 3H). Together, these results show that LRF alters largely distinct sets of genes in male and female hearts.

Between-sex comparisons within each feeding condition, performed on depth-matched matrices, identified 5,419 differentially expressed genes under ALF and 5,483 under LRF (Figures S9A–S9D), an order of magnitude more than the feeding-condition comparisons within each sex. Genes higher in females were enriched for morphogenesis and Wnt signaling terms, and genes higher in males for ribosome and ribonucleoprotein complex biogenesis, under both conditions (Figures S9E and S9F). Of these genes, 3,869 (55.0%) were differentially expressed under both feeding conditions and changed concordantly (Figures S9G and S9H).

### Transcript-level responses to LRF are sex-specific and extend beyond gene expression changes

Transcript-level differential expression was assessed with DESeq2 comparing LRF to ALF within each sex, with zeitgeber time included as a covariate and significance defined as Benjamini–Hochberg adjusted p-value (padj) < 0.05 and |log_2_FC| ≥ 1. Most transcripts were expressed under both feeding conditions in each sex (63.2% in females, 69.3% in males), with 18.5% detected exclusively under ALF and 18.3% exclusively under LRF in females, and 16.5% and 14.3%, respectively, in males (Figures 4A and 4B, and S10A and S10B). Condition-specific transcript sets overlapped minimally between sexes, with 731 of the ALF-specific and 625 of the LRF-specific transcripts shared, against 6,440–9,775 transcripts specific to a single sex and condition (Figure S10E). Female ALF-specific transcripts were enriched for regulation of mRNA splicing and mRNA processing, whereas male LRF-specific transcripts were enriched predominantly for DNA repair and cellular stress response terms (Figures S10C and S10D); transcripts shared between sexes were enriched for neuronal morphogenesis and cAMP response under ALF and for nuclear RNA export and microtubule-based transport under LRF (Figures S10F and S10G). Novel transcript isoforms of known genes comprised 18–19% of condition-specific transcripts in females and 13% in males (Figure S10H), and 72–79% arose from genes expressed under both feeding conditions (Figure S10I).

**Figure 4.**
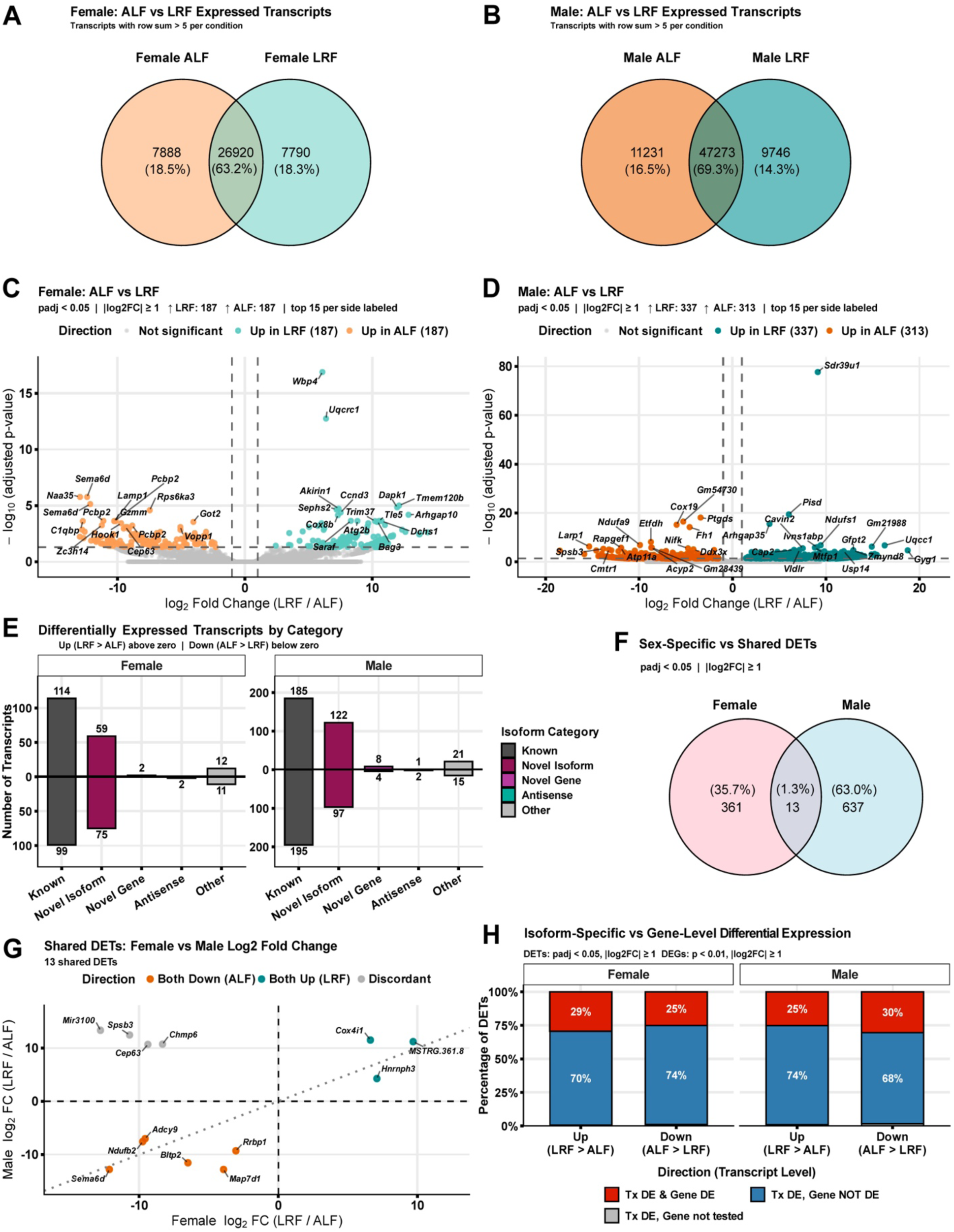
Differential cardiac transcript expression under light-cycle time-restricted feeding reveals widespread transcript isoform-specific regulation. (A and B) Venn diagrams of expressed transcripts between ad libitum feeding (ALF) and light-cycle time-restricted feeding (LRF) in females (A) and males (B). A transcript was considered expressed within a sex-condition group if the sum of raw counts across all four samples exceeded 5. Numbers and percentages indicate condition-specific and shared transcript counts. Shared transcripts were tested for differential expression in (C) and (D). (C and D) Volcano plots of transcript-level differential expression between ALF and LRF in females (C) and males (D). The x-axis shows log_2_ fold change (log_2_FC; LRF/ALF); the y-axis shows −log_10_(adjusted p-value [padj]). Differentially expressed transcripts (DETs) were defined as padj < 0.05 and |log_2_FC| ≥ 1 (dashed lines). The 15 most significant DETs per direction are labeled. Teal, higher under LRF; orange, higher under ALF; gray, not significant. Females: 187 higher under LRF, 187 higher under ALF. Males: 337 higher under LRF, 313 higher under ALF. (E) Bidirectional bar charts showing the number of DETs classified by gffcompare transcript isoform annotation category: charcoal, Known; magenta, Novel Isoform; pink, Novel Gene; green, Antisense; gray, Other, for females (left) and males (right). Bars above zero indicate transcripts higher under LRF; bars below zero indicate transcripts higher under ALF. Numbers on bars indicate transcript counts per category and direction. (F) Venn diagram showing sex-specific and shared DETs. Female-specific: 361 (35.7%); male-specific: 637 (63.0%); shared: 13 (1.3%). (G) Scatter plot comparing log_2_FC (LRF/ALF) between females (x-axis) and males (y-axis) for the 13 shared DETs. Orange, concordant upregulation under ALF; teal, concordant upregulation under LRF; gray, discordant direction between sexes. All 13 transcripts are labeled. (H) Stacked bar charts showing the percentage of DETs with concurrent gene-level differential expression (DE; Tx DE, transcript-level differential expression), stratified by transcript-level direction and sex. Red, transcript DE with concurrent gene-level DE (gene-level p < 0.01, |log_2_FC| ≥ 1); blue, transcript DE but parent gene not DE; gray, transcript DE but parent gene not tested or lacking a valid reference identifier. DET threshold: padj < 0.05, |log_2_FC| ≥ 1. N = 4 biological replicates per sex per condition (one per zeitgeber time [ZT]).

LRF altered expression of 374 transcripts in females (187 higher under LRF, 187 higher under ALF) and 650 transcripts in males (337 higher under LRF, 313 higher under ALF; Figures 4C and 4D). Known annotated transcripts formed the largest category in both sexes, but novel transcript isoforms of known genes accounted for approximately one third of differentially expressed transcripts (134 in females, 219 in males), with smaller contributions from novel gene, antisense, and other categories (Figure 4E). The top 50 differentially expressed transcripts in each sex are shown in Figure S12.

Cross-sex comparison showed that 361 differentially expressed transcripts (35.7%) were female-specific and 637 (63.0%) male-specific, with 13 transcripts (1.3%) shared between sexes (Figure 4F). Among shared transcripts, *Cox4i1*, *Hnrnph3*, and one novel transcript (StringTie2 identifier MSTRG.361.8) were higher under LRF in both sexes, while *Ndufb2*, *Adcy9*, *Sema6d*, *Rrbp1*, *Bltp2*, and *Map7d1* were higher under ALF in both sexes. *Mir3100*, *Spsb3*, *Cep63*, and *Chmp6* changed in opposite directions between sexes (Figure 4G).

In females, 70% of transcripts higher under LRF and 74% of those higher under ALF arose from genes not differentially expressed at the gene level; in males, these proportions were 74% and 68% (Figure 4H). This shows that most differentially expressed transcripts had no corresponding gene-level change.

Between-sex comparisons within each feeding condition, performed on depth-matched matrices, identified 8,487 differentially expressed transcripts under ALF and 8,514 under LRF (Figures S11A–S11D), exceeding the feeding-condition comparisons within each sex by more than tenfold. Novel transcript isoforms accounted for approximately one quarter of these transcripts; 4,811 (39.5%) were differentially expressed under both feeding conditions and changed concordantly; and 72–88% also showed a corresponding gene-level change (Figures S11E–S11J and S13).

### Transcript usage shifts under LRF are widespread, sex-specific, and independent of gene output

Differential transcript usage was assessed with DRIMSeq comparing LRF to ALF within each sex, with zeitgeber time included as a covariate and significance defined as gene-level Benjamini–Hochberg false discovery rate (BH-FDR) < 0.05. LRF induced transcript isoform usage changes in both sexes, with 174 genes showing significant DTU in females and 339 in males (Figure 5A). The magnitude of transcript isoform usage change was similar between sexes, and DTU genes were predominantly classified as major transcript isoform switches, in which the dominant transcript isoform changed between conditions (max |Δ transcript isoform fraction| ≥ 0.15), or partial redistribution of transcript isoform proportions without a change in the dominant transcript isoform (max |Δ transcript isoform fraction| ≥ 0.08), with minor shifts rare in both sexes. In females, 91 genes (52%) showed major switches and 79 (45%) showed partial redistribution; in males, 181 genes (53%) showed major switches and 157 (46%) showed partial redistribution (Figures 5B and 5C).

**Figure 5.**
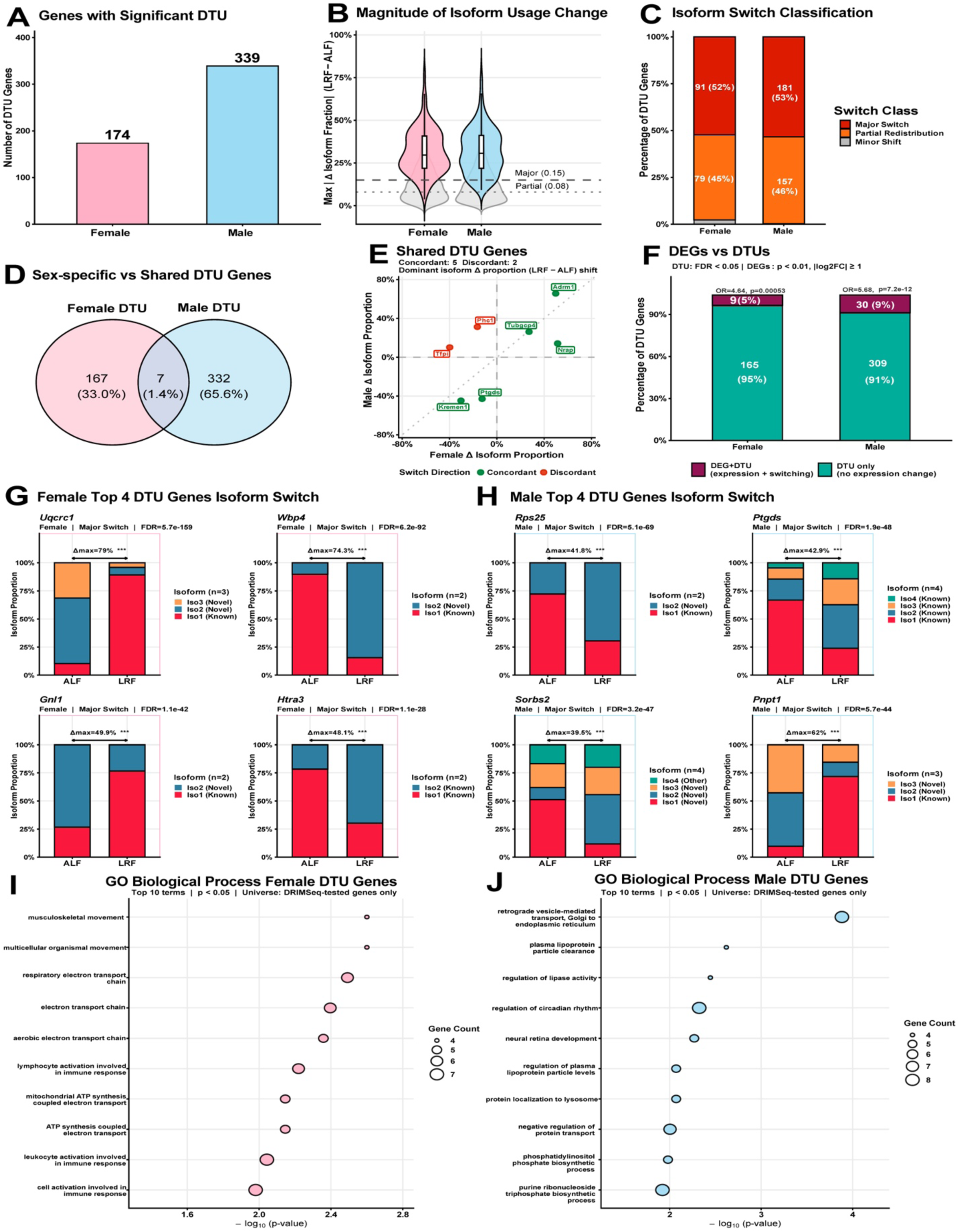
Transcript isoform switching under light-cycle time-restricted feeding is sex-specific and largely independent of gene expression changes. (A) Number of genes with significant differential transcript usage (DTU; DRIMSeq Benjamini–Hochberg false discovery rate [BH-FDR] < 0.05) in females (174; pink) and males (339; blue). (B) Violin plots showing the distribution of maximum absolute transcript isoform proportion change (|Δ|; light-cycle time-restricted feeding [LRF] − ad libitum feeding [ALF]) per gene for all DRIMSeq-tested genes (gray) and significant DTU genes (pink, females; blue, males). Dashed line, major switch threshold (|Δ| ≥ 0.15); dotted line, partial redistribution threshold (|Δ| ≥ 0.08). (C) Classification of DTU genes by switch magnitude. Major switch (|Δ| ≥ 0.15; red): female 91 (52%), male 181 (53%). Partial redistribution (|Δ| ≥ 0.08; orange): female 79 (45%), male 157 (46%). Minor shift (gray): female 4 (2%), male 1 (<1%). (D) Venn diagram of sex-specific and shared DTU genes. Female-only: 167 (33.0%); male-only: 332 (65.6%); shared: 7 (1.4%). (E) Scatter plot of dominant transcript isoform proportion change (Δ; LRF − ALF) for the 7 shared DTU genes in females (x-axis) versus males (y-axis). Green, concordant switching direction (5 genes); red, discordant switching direction (2 genes). (F) Stacked bar charts showing the proportion of DTU genes with concurrent gene-level differential expression (DE; gene-level p < 0.01, absolute log_2_ fold change [|log_2_FC|] ≥ 1). Maroon, DTU with a concurrent differentially expressed gene (DEG); teal, DTU without gene-level DE. Females: 9 (5%) DEG+DTU, 165 (95%) DTU only. Males: 30 (9%) DEG+DTU, 309 (91%) DTU only. Odds ratio (OR) and p-value from Fisher’s exact test are shown per sex. (G and H) Representative transcript isoform proportion plots for the top 4 DTU genes per sex, ranked first by switch magnitude class and then by DRIMSeq BH-FDR, in females (G: *Uqcrc1*, *Wbp4*, *Gnl1*, *Htra3*) and males (H: *Rps25*, *Ptgds*, *Sorbs2*, *Pnpt1*). Stacked bars show mean transcript isoform proportions under ALF (left) and LRF (right). Maximum absolute transcript isoform proportion change (Δmax) and gene-level BH-FDR significance are annotated above each plot (***, BH-FDR < 0.001). (I and J) Gene Ontology (GO) biological process enrichment for DTU genes in females (I) and males (J). The x-axis shows −log_10_(p-value); dot size is proportional to gene count. Top 10 enriched terms are shown (p < 0.05). N = 4 biological replicates per sex per condition (one per zeitgeber time [ZT]).

Cross-sex comparison showed that 167 DTU genes (33.0%) were detected only in females and 332 (65.6%) only in males, with 7 genes (1.4%) shared between sexes (Figure 5D). Among the shared genes, five showed concordant transcript isoform shifts in both sexes, including *Adrm1*, *Nrap*, *Kremen1*, *Tubgcp4*, and *Ptgds*, while *Tfpi* and *Phc1* showed discordant switching patterns between sexes (Figure 5E). Transcript isoform proportion plots for all seven shared genes are shown separately for females and males (Figure S15).

The majority of DTU events occurred without gene-level differential expression. In females, 95% of DTU genes showed transcript isoform switching without a corresponding gene expression change, and in males this proportion was 91% (Figure 5F). DTU genes were nonetheless significantly enriched for concurrent gene-level differential expression in both sexes (female odds ratio [OR] = 4.64, p = 5.3 × 10^−4^; male OR = 5.68, p = 7.2 × 10^−12^), although the absolute overlap remained modest, at 5% of female and 9% of male DTU genes. Representative top DTU genes illustrate the scale of transcript isoform proportion changes between feeding conditions. In females, *Uqcrc1*, *Wbp4*, *Gnl1*, and *Htra3* showed major transcript isoform switches involving both known and novel transcript isoforms (Figure 5G). In males, *Rps25*, *Ptgds*, *Sorbs2*, and *Pnpt1* similarly showed large transcript isoform proportion shifts (Figure 5H). Transcript isoform proportion plots for additional female and male cardiac DTU genes, arranged by functional category, are shown in Figures S16 and S17, respectively.

GO enrichment (p < 0.05) of DTU genes revealed distinct functional profiles in each sex (Figures 5I and 5J). In females, the most significantly enriched terms were musculoskeletal movement and multicellular organismal movement, followed by terms associated with mitochondrial energy metabolism, including respiratory electron transport chain and ATP synthesis-coupled electron transport, as well as lymphocyte and leukocyte activation (Figure 5I). In males, enriched terms included retrograde vesicle transport from the Golgi to the endoplasmic reticulum, plasma lipoprotein particle clearance, regulation of lipase activity, and regulation of circadian rhythm (Figure 5J). Together, these results show that LRF alters transcript isoform usage in the heart in a predominantly sex-specific manner and largely independently of gene-level differential expression.

Between-sex comparisons within each feeding condition, performed on the depth-matched matrix, identified 177 DTU genes under ALF and 166 under LRF (Figures S14A–S14C), comparable to or fewer than the feeding-condition comparisons within each sex, unlike at the gene and transcript expression levels where sex effects were an order of magnitude larger. Most sex-related DTU genes were condition-specific; more than half also showed sex-differential expression consistent with the large number of sex-biased genes, and GO enrichment differed by condition, with secretion-related processes under ALF and developmental growth under LRF (Figures S14D–S14J). To understand the structural basis of these transcript isoform switches, we next characterized the splicing events underlying DTU genes in each sex.

### Transcript switches under LRF involve multiple splicing event types in both sexes

Splicing events were annotated for all significant DTU genes using IsoformSwitchAnalyzeR, examining eight event types: exon skipping, multi-exon skipping, mutually exclusive exons, intron retention, alternative 5′ splice site (Alt. 5′), alternative 3′ splice site (Alt. 3′), alternative transcription start site (Alt. TSS), and alternative transcription termination site (Alt. TTS). For each DTU gene, we defined a switch pair of transcript isoforms representing the two sides of its transcript isoform-usage change: the transcript isoform dominant under ALF and the transcript isoform dominant under LRF. For major switch genes, these are by definition two different transcript isoforms, so the switch pair captures the actual identity change. Partial redistribution and minor shift genes have no such identity change, so their switch pair was instead the transcript isoform with the largest proportion gain and the transcript isoform with the largest proportion loss. All analyses were performed independently within each sex. The top 20 DTU genes ranked by transcript isoform proportion shift are shown for each sex in Figure 6A, the majority of which were classified as major switch genes in both sexes.

**Figure 6.**
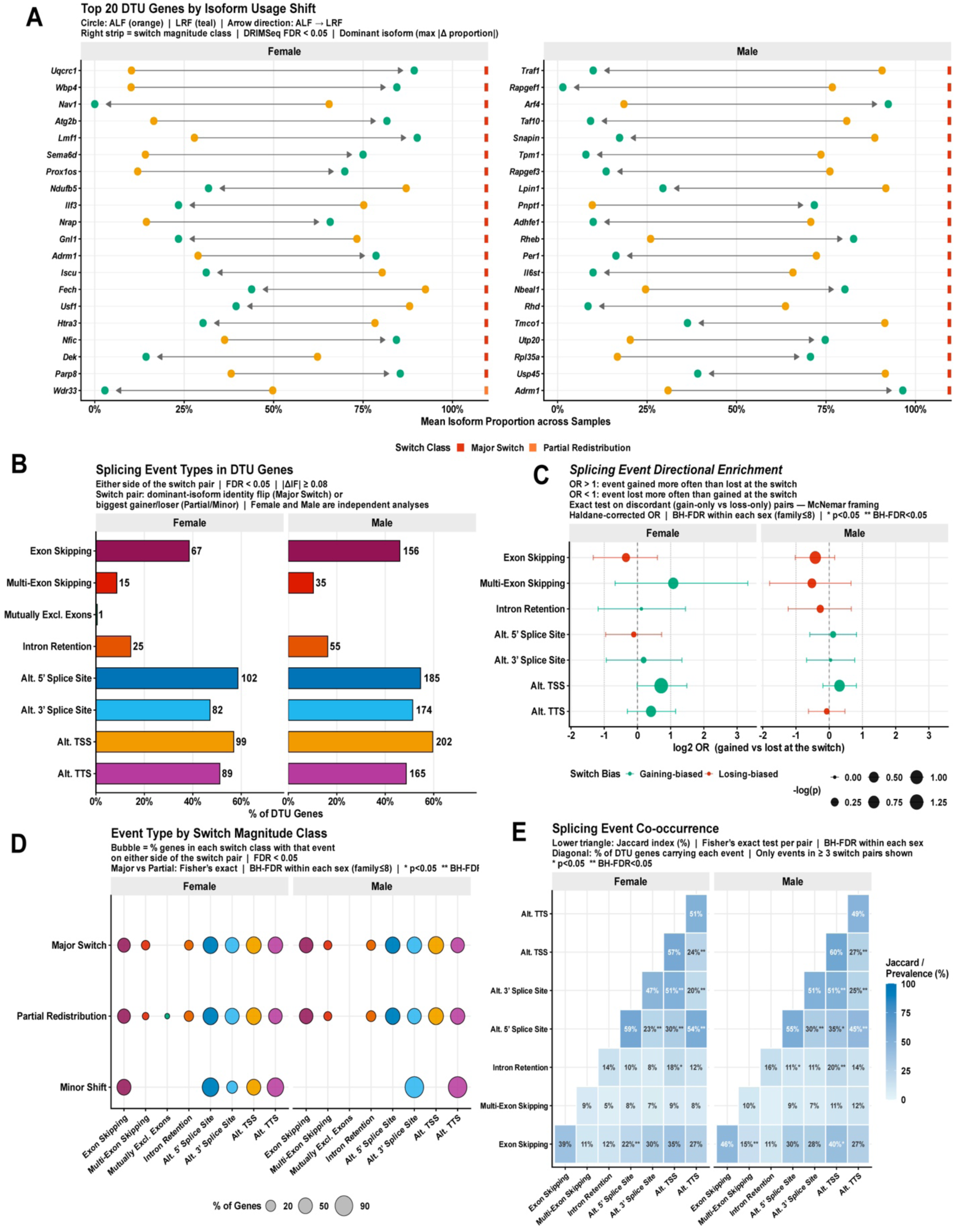
Splicing architecture of differentially used transcript isoforms reveals co-occurring alternative splice site and transcription boundary events. (A) Top 20 differential transcript usage (DTU) genes per sex ranked by maximum absolute transcript isoform proportion shift. Orange circles, mean proportion under ad libitum feeding (ALF); teal circles, mean proportion under light-cycle time-restricted feeding (LRF); arrows, direction of change from ALF to LRF. Right strip indicates switch magnitude class (red, major switch; orange, partial redistribution). (B) Prevalence of eight annotated splicing event types across DTU genes in females (left) and males (right). For each DTU gene, a switch pair of transcript isoforms was defined (dominant transcript isoform under ALF versus LRF for major switch genes; largest-gain versus largest-loss transcript isoform for partial redistribution and minor shift genes). A gene was counted as carrying an event if either side of its switch pair did. Numbers to the right of bars indicate gene counts; the x-axis shows the percentage of DTU genes carrying each event. (C) Directional enrichment of splicing events at the switch. Events were classified as gained (LRF-side transcript isoform only), lost (ALF-side transcript isoform only), or concordant (both or neither). Odds ratio (OR) = (n gained + 0.5) / (n lost + 0.5); OR > 1, event gained more often than lost (teal); OR < 1, event lost more often than gained (red). Error bars, exact binomial 95% confidence interval. Dot size is proportional to −log_10_(p). No event type reached a Benjamini–Hochberg false discovery rate (BH-FDR) < 0.05 in either sex (minimum uncorrected p = 0.053). (D) Splicing event prevalence by switch magnitude class. Bubble size indicates the percentage of genes in each class (major switch, partial redistribution, minor shift) carrying each event type. No comparison between major switch and partial redistribution genes reached significance in either sex (Fisher’s exact test, Benjamini– Hochberg corrected). (E) Co-occurrence of splicing event pairs across DTU genes. Lower triangle, Jaccard index (%) with significance from Fisher’s exact test; diagonal, event prevalence. Only event types present in at least three DTU genes were included. *p < 0.05; **BH-FDR < 0.05. Eight splicing event types were examined: exon skipping, multi-exon skipping, mutually exclusive exons, intron retention, alternative 5′ splice site (Alt. 5′), alternative 3′ splice site (Alt. 3′), alternative transcription start site (Alt. TSS), and alternative transcription termination site (Alt. TTS). ΔIF, transcript isoform fraction change (LRF − ALF). All analyses were restricted to genes with significant DTU (DRIMSeq BH-FDR < 0.05), performed independently within each sex. N = 4 biological replicates per sex per condition (one per zeitgeber time [ZT]).

Alternative splice site and transcription boundary events were the most frequently detected event types in both sexes (Figure 6B). In females, Alt. 5′ splice site was the most prevalent event (n = 102, 58.6%), followed by Alt. TSS (n = 99, 56.9%), Alt. TTS (n = 89, 51.1%), and Alt. 3′ splice site (n = 82, 47.1%). In males, Alt. TSS was the most prevalent (n = 202, 59.6%), followed by Alt. 5′ splice site (n = 185, 54.6%), Alt. 3′ splice site (n = 174, 51.3%), and Alt. TTS (n = 165, 48.7%). Exon skipping was also common, detected in 67 (38.5%) female and 156 (46.0%) male DTU genes. Intron retention affected 25 (14.4%) female and 55 (16.2%) male DTU genes; multi-exon skipping was infrequent (15 female DTUs, 8.6%; 35 male DTUs, 10.3%), and mutually exclusive exons were detected in only one female DTU gene and were absent in males.

We next asked whether any event type was preferentially gained or lost as transcript isoforms switched, and whether any event type distinguished major switch genes from partial redistribution genes. For each event type, genes were classified as gained (event present on the LRF transcript isoform only), lost (event present on the ALF transcript isoform only), or unchanged (event present on both transcript isoforms or neither); only gained and lost genes are informative about direction and were compared using an exact test analogous to McNemar’s test, with Benjamini–Hochberg correction applied within each sex across the eight event types. No event type showed a significant directional bias toward the incoming or outgoing transcript isoform in either sex after correction (minimum uncorrected p = 0.053; Figure 6C). Separately, we tested whether each event type’s prevalence differed between major switch and partial redistribution genes using Fisher’s exact test, again with Benjamini–Hochberg correction within each sex. No event type differed significantly between switch magnitude classes in either sex (Figure 6D). Together, these results indicate that while specific splicing mechanisms are broadly prevalent among DTU genes (Figure 6B) and frequently co-occur within the same genes (Figure 6E), no single mechanism preferentially drives the direction or magnitude of transcript isoform switching under LRF.

Co-occurrence analysis revealed that alternative splice site and transcription boundary events co-occurred non-randomly across DTU genes in both sexes (Figure 6E). Alt. 3′ splice site and Alt. TSS showed the strongest co-occurrence in both females (Jaccard = 51%, BH-FDR < 0.05) and males (51%, BH-FDR < 0.05). Alt. 5′ splice site co-occurred significantly with Alt. TTS (female: 54%; male: 45%; both BH-FDR < 0.05), with Alt. 3′ splice site (female: 23%; male: 30%; both BH-FDR < 0.05), and with Alt. TSS (female: 30%, BH-FDR < 0.05; male: 35%, nominal p < 0.05). Alt. TSS also co-occurred significantly with Alt. TTS (female: 24%; male: 27%; both BH-FDR < 0.05). Intron retention co-occurred significantly with Alt. TSS in both sexes (female: 18%, nominal p < 0.05; male: 20%, BH-FDR < 0.05) and with Alt. 5′ splice site in males (11%, nominal p < 0.05). Exon skipping co-occurred with Alt. 5′ splice site in females (22%, BH-FDR < 0.05), with multi-exon skipping in males (15%, BH-FDR < 0.05), and with Alt. TSS in males (40%, nominal p < 0.05). Together, these results show that transcript isoform switching under LRF is characterized by alternative splice site and transcription boundary events that frequently co-occur within the same DTU genes in both sexes, with intron retention and exon skipping additionally linked to transcription start site usage.

## DISCUSSION

A behavioral change in feeding pattern induced by LRF, through the circadian misalignment between central and peripheral clocks, is sufficient to reorganize the cardiac transcriptome. As expected, LRF phase-shifted clock gene oscillations in both sexes, confirming that the intervention engaged the circadian system. Beyond this, LRF induced widespread changes in which transcript a gene produces, largely without altering total gene output. The genes affected were almost entirely sex-specific, yet the scale of the transcript remodeling and its independence from gene expression were consistent across sexes. These findings identify transcript-level regulation as a previously underrecognized layer of the cardiac response to temporal metabolic disruption, one that gene-level transcriptomics alone cannot detect.

Altered metabolic cues in the form of time-restricted feeding to the light cycle uncouple peripheral tissue clocks, including the heart, from the SCN by up to 12 hours, without corresponding changes in SCN gene expression rhythms^7,11^. Consistent with these studies, we found that *Bmal1*, *Per2*, *Cry2*, *Rev-erbα* (*Nr1d1*), *Dbp*, and other circadian clock genes were phase-shifted by LRF in our dataset, previously described in both male^10^ and female mice^12^. Heart-specific loss of *Bmal1* reduces expression of genes in the fatty acid oxidative pathway, the tricarboxylic acid cycle, and the mitochondrial respiratory chain, and causes progressive heart failure^23^, indicating the cardiomyocyte clock controls programs beyond timekeeping. The clock also regulates RNA splicing, as loss of *Clock* or *Bmal1* disrupts genome-wide alternative splicing in pancreatic beta cells through the RNA-binding protein THRAP3^24^, and alternative splicing responds to food-related cues^25^. In our dataset, *Thrap3* underwent transcript isoform switching in male hearts under LRF, providing independent biological validation that the transcript-level changes we detect reflect genuine regulatory circuitry already implicated in clock-driven splicing. Long-read sequencing has so far been applied almost exclusively to characterize transcript-level dysregulation in disease states^21,22,26–28^. Our findings show that a comparable degree of transcript reorganization can be induced by an everyday physiological behavior in an otherwise healthy heart, raising the possibility that some disease-associated transcript signatures reflect behavioral or environmental exposure rather than pathology itself.

Long-read sequencing also revealed the extent to which the cardiac transcriptome remains unannotated. Approximately one in five cardiac transcripts represented a novel transcript isoform of a known gene (22.5% in females, 22.7% in males), consistent with published long-read studies detecting 15–31% novel transcript isoforms in cardiac tissue^26,27^. Among major switch genes, a novel transcript was the dominant transcript isoform under LRF in 37% of cases in females and 38% of cases in males, indicating that the cardiac response to LRF substantially engages unannotated transcript variants.

Prior timed-feeding gene expression studies used common methods such as qPCR and short-read RNA sequencing, which cannot resolve full-length or novel transcripts^11,12^, and would have missed the primary response we report. 68–74% of transcript-level and 91–95% of transcript isoform-usage changes occurred without any gene-level differential expression. Caloric restriction drives similar transcript isoform remodeling across six tissues (white adipose tissue, liver, hypothalamus, gastrocnemius muscle, testes, and stomach), with 94% of loci showing differential transcript isoform usage exhibiting no concurrent differential expression^29^. LRF alone, without caloric restriction, is therefore sufficient to engage this response in the heart.

LRF engaged sex-specific pathways through transcript isoform switching. In female hearts, DTU genes were enriched for mitochondrial energy metabolism, with switches spanning complexes I and III, including *Uqcrc1,* whose knockdown produces left ventricular structural changes and mitochondrial defects^30^. Cardiac *Bmal1* loss similarly disrupts mitochondrial morphology and function^23^, and estrogen receptor β modulates cardiac mitochondrial respiration^31^, positioning the female mitochondrial program at the intersection of clock disruption and sex hormone signaling. Contractile genes were also affected, including *Bag3,* whose myofilament expression is sex-dependent in dilated cardiomyopathy^32^. The enrichment for lymphocyte and leukocyte activation among female DTU genes may reflect circadian regulation of cardiac immune cell activity, though the specific immune populations contributing to this signal in bulk ventricular tissue remain to be identified. In male hearts, DTU genes were enriched for lipid handling, mitochondrial respiratory chain function, and circadian rhythm regulation, with switches in *Atp5f1a*, *Uqcrc2*, *Ndufb8*, and other respiratory chain subunits. *Tpm1* underwent a major transcript isoform switch; *Tpm1* transcript isoform composition changes are associated with dilated cardiomyopathy and heart failure^33^, with PTBP1 and RBFOX2 regulating its splicing antagonistically^34^. *Cacna1c*, which produces transcript isoforms with distinct QT interval effects^35^ and is one of the *RBM20*-regulated transcripts implicated in dilated cardiomyopathy^15^, showed transcript isoform redistribution. *Per1* underwent a major transcript isoform switch, showing that LRF remodels core clock output at the transcript isoform level. Together, these findings show that LRF drives largely distinct transcript isoform programs in male and female hearts, with mitochondrial remodeling as a shared point of convergence.

Fewer than 2% of changes were shared between sexes at the gene, transcript, or transcript-usage level. This near-complete divergence is consistent with what the baseline transcriptome predicts^16^. Sex was the single largest source of transcriptomic variance in this dataset, explaining 56.2% of transcriptome variance, and between-sex comparisons within each feeding condition identified an order of magnitude more differentially expressed genes and transcripts than did the feeding-condition comparisons within each sex. The cardiomyocyte clock actively maintains the sex-specific landscape. BMAL1 regulates sex-specific expression of ion channel and autonomic signaling genes, and cardiomyocyte-specific *Bmal1* knockout substantially reduces transcriptomic sex differences^16,36^. LRF acts on a transcriptome that is already sexually dimorphic. The resulting divergence in transcript remodeling does not reflect male and female hearts responding differently to the same signal; it reflects the same signal acting on different molecular substrates. The few shared responses, including the concordant *Ptgds* switch linking circadian disruption to inflammatory prostaglandin output, likely reflect the subset of transcripts regulated by sex-independent clock targets. For long-read studies of transcript-level disease signatures, this raises a specific concern. Pooling sexes, or analyzing only one, risks either masking a true shared response or misattributing a sex-specific baseline difference to the perturbation itself. Sex should be treated as a biological variable at the transcript level, not only at the gene level.

Despite targeting different genes, LRF altered transcript architecture through the same types of splicing events in both sexes. Alternative splice site and transcription boundary events dominated in both sexes, and no event type showed significant directional bias in either sex. These events co-occurred within the same genes more often than expected by chance, showing that LRF restructures transcript architecture at multiple points simultaneously rather than at isolated splice sites. The splicing machinery itself was a target. *Hnrnph3* was concordantly higher under LRF in both sexes, while sex-specific splicing regulators underwent transcript isoform switches, including *Celf1*, whose re-expression in adult hearts disrupts transverse tubule organization and Ca^2+^ handling^37^, and *Ptbp1*, whose switch follows a pattern seen in diabetic cardiomyopathy^38^. This remodeling of the splicing regulators themselves suggests a feed-forward mechanism in which LRF-driven transcript isoform changes in splicing factors propagate further transcript remodeling across the transcriptome.

LRF reorganizes the cardiac transcriptome primarily at the transcript isoform level. Approximately one in five (>20%) cardiac transcripts is a novel transcript isoform that actively participates in this response, revealing a transcriptome more complex than current annotations capture. The remodeling is sex-specific in the pathways affected, yet both sexes execute it through a shared post-transcriptional architecture of co-occurring splicing events. This sex-specific molecular signature may be clinically relevant. Irregular meal timing is linked to cardiovascular disease^1^. The sex difference in rhythmic mRNA expression that we observe here may be associated with sex-specific differences in the timing of cardiac arrhythmias, though this exact connection remains to be tested^36^. Transcript isoform-level remodeling therefore represents a candidate molecular mechanism underlying this sex-specific vulnerability, an early cardiac signature of metabolic misalignment that gene-level transcriptomics cannot detect.

Whether these RNA-level changes alter cardiac electrical activity, contractile function, or protein output remains a key open question.

## LIMITATIONS OF THE STUDY

One limitation of this study was the sample size (N = 4 per sex per condition, with one biological replicate per time point), which limited statistical power for formal rhythmicity modeling and precise phase-shift quantification. Male and female libraries were prepared in separate batches at different sequencing depths; primary comparisons were performed within each sex, and between-sex comparisons used depth-matched matrices, though the larger number of male DTU events may partly reflect greater sequencing depth. Novel transcript models were supported by independent short-read junction data, CAGE peaks, and polyA motifs; targeted reverse-transcription PCR (RT-PCR) and protein-level validation of key transcript isoform switches represent important future steps. Bulk ventricular sequencing cannot resolve cell-type-specific contributions. Findings are from a single inbred strain on a two-week protocol and require testing in additional genetic backgrounds, longer time frames, and humans with irregular eating patterns. Future work should also identify the RNA-binding proteins that relay feeding-time signals to the spliceosome and define how sex hormones contribute to the sex-divergent transcript isoform responses observed here.

## MATERIAL AND METHODS

Materials and methods are provided as a **Supplementary Data** document.

## DATA AVAILABILITY

**Supplementary Figures S1-S17** are provided in **Supplementary Data** document. **Extended Data 1 and Data 2, Table S1** and have been deposited in Figshare (10.6084/m9.figshare.33960214). The sequencing data is pending GEO accession number.

## AUTHOR CONTRIBUTIONS

Conceptualization, B.P.D., E.A.S., and Y.W.; Methodology, S.N., A.P., and T.S; Investigation, S.N., B.P., Y.W., and A.P.; Writing – Original Draft, B.P.D., E.A.S., Y.W., and S.N.; Writing – Review & Editing, B.P.D., E.A.S., Y.W., S.N., A.P., and B.P.; Funding Acquisition, B.P.D., E.A.S., and Y.W.; Resources, B.P.D., E.A.S., and Y.W.; Supervision, B.P.D., E.A.S., and Y.W.

## ACKNOWLEDGMENTS

National Heart, Lung, and Blood Institute R01HL172813 to B.P.D. and E.A.S. National Institute of Arthritis and Musculoskeletal and Skin Diseases R00AR081367 to Y.W. A.P. is supported by Pathway to Independence Grant from Diabetes and Research Priority Area at the Barnstable Brown Diabetes Center, University of Kentucky. The content is solely the responsibility of the authors and does not necessarily represent the official views of the NIH. Claude AI (Sonnet 4.6) was used for formatting and copyediting purposes.

