## Supplementary Data for "Light-cycle time-restricted feeding remodels a hidden layer of the cardiac transcriptome through sex-specific transcript switching"

#### MATERIAL AND METHODS

##### Animals

All animal procedures were performed in a facility accredited by the Association for Assessment and Accreditation of Laboratory Animal Care (AAALAC) and were approved by the Institutional Animal Care and Use Committee (IACUC protocol #2019-3304) at the University of Kentucky. Wild-type SV129 male and female mice (*Mus musculus*) aged 6 months were used. Mice were housed at 24 ± 1 °C under a 12-hour (h) light (200 lux) / 12 h dark (0 lux) cycle with ad libitum access to food and water for one week before the start of the experiment.

##### Feeding paradigm and tissue collection

To initiate the restricted feeding protocol, food was removed at zeitgeber time 9 (ZT9; 9 h after lights on) on the first day. Beginning the following day, food availability was restricted to a 7 h window during the light cycle between ZT2 and ZT9, and this schedule was maintained for two weeks (light-cycle-restricted feeding, LRF). The ad libitum feeding (ALF) group continued to have unrestricted access to food and water for the same two-week period.

After two weeks, ventricular tissue was collected at four zeitgeber time points (ZT1, ZT7, ZT13, and ZT19) for each sex and feeding condition to capture the full 24 h cycle. Sample size was N = 4 per sex per condition, with one biological replicate per zeitgeber time point, giving 16 samples in total. Tissues were flash-frozen in liquid nitrogen and stored at −80 °C.

##### RNA isolation

For each sample, 75–100 mg of tissue was transferred into 1.5 mL DNA LoBind tubes containing 100–200 zirconium oxide beads (1 mm, RNase-free) and 1 mL TRIzol reagent. Tissue was lysed and homogenized using a Bullet Blender for 1–2 min at speed 8. Following lysis, samples were centrifuged at 12,000 × g for 5 min at 4 °C. The clear supernatant was transferred to a new tube and incubated for 5 min to allow complete dissociation of the nucleoprotein complex. Chloroform (0.2 mL) was added and the sample was thoroughly mixed. After a 5 min incubation, samples were centrifuged at 12,000 × g for 15 min at 4 °C. The aqueous phase containing RNA was carefully transferred to a new tube without disturbing the interphase. An equal volume of isopropanol was added, and the sample was vortexed and incubated at room temperature (25 °C) for 10 min, followed by centrifugation at 12,000 × g for 10 min at 4 °C. The supernatant was removed and the RNA pellet was washed twice with 1 mL 75% ethanol, briefly vortexed, and centrifuged at 7,500 × g for 5 min at 4 °C. The supernatant was discarded, the pellet was air-dried for 5–10 min, and RNA was resuspended in 30 µL RNase-free water. The final RNA product was incubated with DNase I and SUPERase-In for 15 min at 37 °C. RNA quality and concentration were assessed using a Qubit fluorometer and an Agilent TapeStation. All RNA samples had an RNA integrity number equivalent (RINe) greater than 7.

**Nanopore cDNA library preparation**

Long-read RNA sequencing libraries were generated using the Oxford Nanopore Technologies cDNA-PCR Barcoding Kit V14 according to the manufacturer's instructions. For each sample, 500 ng of total RNA was used as input for reverse transcription and strand-switching, followed by PCR amplification to generate full-length cDNA libraries compatible with R10.4.1 PromethION flow cells. Male and female samples were prepared and sequenced in two independent batches; the resulting difference in sequencing depth was addressed by depth matching (see below). Following PCR amplification, libraries were purified using AMPure XP beads, quantified using a Qubit fluorometer, and assessed for fragment size distribution using an Agilent Bioanalyzer. Within each batch, libraries were pooled in equimolar ratios based on fragment length and concentration. Rapid sequencing adapters were added according to the manufacturer's instructions, and prepared libraries were loaded onto R10.4.1 PromethION flow cells.

##### Nanopore sequencing and basecalling

Sequencing was performed on the Oxford Nanopore Technologies PromethION platform using 72 h sequencing runs controlled by MinKNOW software. Raw electrical signal data were recorded in POD5 format. Basecalling and barcode demultiplexing were performed simultaneously using Dorado (v0.9.1) with the super-accurate (SUP) model (dna_r10.4.1_e8.2_400bps_sup@v5.0.0). Barcode detection was configured using the SQK-PCB114-24 kit setting, enabling automatic demultiplexing of barcoded samples during basecalling. Basecalled and demultiplexed reads were exported in BAM format for downstream analysis.

##### Raw read processing and genome alignment

Basecalled and demultiplexed BAM files were converted to FASTQ format using samtools (v1.12); no quality or length filtering was applied.

Splice-aware alignment to the mouse reference genome (GRCm39 primary assembly) was performed using minimap2[1] (v2.30-r1287). The minimap2 genome index (GRCm39.mmi) was generated from the reference FASTA using minimap2 -d. A reduced k-mer size (-k14) was used to improve sensitivity given Oxford Nanopore error profiles. Secondary alignments were suppressed (--secondary=no) so that each read retained only its single best (primary) alignment; supplementary alignments were retained.

The reference genome (GRCm39.primary_assembly.genome.fa) and gene annotation from GENCODE release vM36 (gencode.vM36.primary_assembly.annotation.gtf) were used throughout. Alignment output was converted to BAM, sorted, and indexed using samtools[2] (v1.12), and alignment statistics were generated using samtools flagstat.

**Sequencing quality control**

Sequencing quality control was performed for all 16 samples. Read-level metrics (total reads, total bases, mean and median read length, read length N50, and read quality scores) were obtained with NanoStat (v1.6.0) from each FASTQ file. Alignment metrics (total, primary, supplementary, and unmapped alignments; mapped bases; and average alignment length) were obtained with samtools flagstat and samtools stats (v1.12). Metrics were compared across samples to confirm consistency prior to downstream analysis.

**Transcriptome assembly and quantification with StringTie2**

Transcriptome assembly and quantification were performed using StringTie2[3] (v3.0.3) in long-read mode (-L) to accommodate full-length Oxford Nanopore cDNA reads. For each of the 16 samples, reference-guided transcript assembly was conducted using the GENCODE vM36 annotation provided with the -G option, enabling annotation-guided transcript reconstruction while allowing identification of novel transcript isoforms. Per-sample transcript assemblies were merged using stringtie --merge together with the GENCODE vM36 annotation to generate a unified transcriptome reference (merged_isoforms.gtf). This merged annotation ensured consistent transcript identifiers across all samples and retained isoforms detected in any sample.

Transcript abundance was then estimated against the merged transcriptome (-L -e -B -p 8), with each sample written to a separate output directory. The -e option restricted quantification to transcripts defined in the merged reference, ensuring consistent expression estimates across samples, and -B generated Ballgown-compatible output. Gene- and transcript-level raw count matrices were generated from the per-sample GTF files using prepDE.py with a sample list mapping sample identifiers to their corresponding GTF paths, producing a transcript-level count matrix across all 16 samples.

In addition to raw counts, transcripts per million (TPM) values were extracted directly from the TPM attribute field in the per-sample StringTie GTF files. Per-sample TPM tables were merged across all samples to generate an isoform-level TPM expression matrix containing 305,327 transcripts. TPM values were used for visualization and exploratory analyses, whereas all statistical analyses were performed using raw count matrices.

**Depth-matched dataset**

Because male and female libraries were prepared and sequenced in separate batches at different sequencing depths, a depth-matched dataset was generated to provide depth-equalized count matrices for comparisons between sexes and to allow coverage profiles to be compared at matched depth. Primary reads were counted for each aligned BAM file using samtools view -c -F 0x904, excluding secondary, supplementary, and unmapped alignments. The target depth was set to the lowest primary-read count observed across the 16 samples (1,721,349 reads).

Subsampling was performed using samtools (v1.22.1) with samtools view --subsample --subsample-seed 42, which hashes read names so that all alignment records belonging to a given read, including supplementary alignments, are retained or discarded together. Achieved depths ranged from 1,720,316 to 1,722,824 primary reads, and subsampled BAM files were indexed with samtools index.

Depth-matched alignments were requantified using StringTie2 in estimate-only long-read mode (-L -e -B -p 8) against an unmodified copy of the merged transcriptome, ensuring that the transcript set was identical to that used in the primary analysis. Transcript- and gene-level count matrices were generated from the per-sample GTF files using prepDE.py.

**Gene body coverage and coverage bias index**

To assess positional coverage bias along transcripts, gene body coverage profiles were computed using geneBody_coverage.py from the RSeQC package (v5.0.4). A BED12 representation of the GENCODE vM36 primary assembly annotation was used as the transcript reference. Transcripts with a summed exonic length of at least 1,000 bp were retained, and 5,000 of these were selected at random using a fixed seed (42). The resulting reference set spanned all autosomes and both sex chromosomes (median exonic length 2,500 bp, median 7 exons per transcript). Coverage was reported across 100 equally spaced bins spanning each transcript from the 5′ to the 3′ end. Profiles were computed on both the full-depth and depth-matched alignments using this identical transcript set, allowing direct comparison of coverage at the two depths.

To quantify positional coverage bias, a coverage bias index was calculated for each sample as the ratio of mean coverage in the distal 3′ region (bins 81–100) to mean coverage in the proximal 5′ region (bins 1–20). Index values above 1.5 were taken to indicate 3′ bias, values below 0.67 to indicate 5′ enrichment, and intermediate values to indicate approximately uniform coverage across the transcript body.

**Structural annotation of the merged transcriptome using gffcompare**

Structural classification of assembled transcripts was performed using gffcompare[4] (v0.12.10) by comparing the merged transcriptome assembly (merged_isoforms.gtf) against the GENCODE vM36 mouse reference annotation using the -r option. gffcompare produced an annotated transcript model file, a summary statistics file, and a tracking file. gffcompare does not modify transcript structures or abundance estimates; it provides structural annotation describing the relationship of each assembled transcript to the reference transcriptome.

Each transcript was assigned a class code describing its structural relationship to the reference. Exact matches to annotated transcripts were labeled “=”, novel splice variants sharing at least one splice junction with a reference transcript were labeled “j”, intergenic transcripts were labeled “u”, and exonic overlaps on the opposite strand were labeled “x”. Additional class codes represent other structural categories, including transcripts contained within and intron-compatible with a reference transcript (c), containment of a reference transcript (reverse containment, k), retained introns in which all introns are matched or retained (m), retained introns in which not all introns are matched or covered (n), other same-strand overlap with reference exons (o), transcripts fully contained within a reference intron (i), transcripts containing a reference transcript within their intron (y), and possible polymerase run-on with no actual overlap (p). The distribution of class codes across the merged transcriptome was: = 276,294; j 23,002; k 3,646; u 1,473; x 357; n 135; c 133; m 119; o 77; i 77; p 13; y 1.

**Transcript-level count matrix annotation**

Transcript-level raw count matrices generated by prepDE.py contained transcript identifiers and raw counts but lacked structural classification and reference-consistent gene metadata. To annotate the matrix, transcript-level fields were extracted from the gffcompare annotated GTF, including transcript_id, gene_id (StringTie locus identifier), ref_gene_id, ref_tx_id (reference transcript identifier, corresponding to cmp_ref in gffcompare output), class_code, and genomic coordinates (chromosome, start, end, strand). A readable class_description field was added for each class_code to facilitate biological interpretation. Gene-level metadata (gene_name and gene_type) were obtained from the GENCODE vM36 annotation to maintain annotation version consistency. Annotation merging was performed using transcript_id as the unique identifier, preserving isoform-level resolution. When available, ref_gene_id was used as the Ensembl gene identifier; transcripts lacking reference matches retained their original StringTie gene identifiers (MSTRG loci, the prefix StringTie assigns to assembled loci without a reference match). Raw count values were not altered during this step. The resulting annotated transcript-level matrix contained structural classification, genomic coordinates, reference gene metadata, and raw counts for all 16 samples.

**Gene-level count matrix reconstruction and annotation**

Although prepDE.py outputs a gene-level count matrix, this matrix was not used for downstream analyses because it assigns StringTie locus identifiers (XLOC and MSTRG) as gene identifiers rather than reference-consistent Ensembl gene identifiers, which prevents direct biological interpretation and compatibility with reference-based differential expression tools. Instead, gene-level counts were reconstructed from the annotated transcript-level count matrix using transcript-to-gene mappings derived from the gffcompare annotated GTF. For each transcript, the Ensembl ref_gene_id was used when present; transcripts lacking a reference match retained their original StringTie gene identifiers (MSTRG loci) to preserve novel gene loci. Gene-level counts were calculated by summing counts across all transcripts assigned to the same gene identifier. Version suffixes were removed from Ensembl gene identifiers to ensure naming consistency, while StringTie locus identifiers were preserved unchanged. Gene-level metadata (gene_name, gene_type, chromosome, start, end, strand) were extracted from GENCODE vM36 gene annotations and merged with the reconstructed count matrix. Total counts were preserved after aggregation (456,211,712) and per-sample library sizes were unchanged. The final annotated gene-level matrix contained 78,298 reference genes and 540 novel loci, for a total of 78,838 genes. The gene-level count matrix retained all genes, including those with low or zero counts across samples; expression filtering was applied separately within each differential expression analysis, as described below.

**Isoform-level TPM matrix re-annotation**

The isoform-level TPM matrix described above (305,327 transcripts across 16 samples) was annotated using the same procedure applied to the transcript-level count matrix, and TPM values were neither recalculated nor modified. The final annotated isoform-level TPM matrix retained one row per transcript and contained structural classification, reference-consistent gene metadata, genomic coordinates, and TPM values for all 16 samples.

**Short-read RNA sequencing data**

Publicly available short-read RNA sequencing data from mouse ventricle were obtained from the Gene Expression Omnibus (GEO): GSE262714[5] for orthogonal validation of long-read transcript models. Eight vehicle-treated samples were selected, comprising both sexes at four circadian time (CT) points (CT26, CT30, CT38, and CT42): GSM8174356, GSM8174358, GSM8174362, and GSM8174364 (male), and GSM8174332, GSM8174334, GSM8174338, and GSM8174340 (female). Each GEO sample had been sequenced across three Sequence Read Archive (SRA) runs. Runs were downloaded using prefetch and converted to FASTQ using fasterq-dump --split-files (SRA Toolkit), then concatenated per sample in identical order for read 1 and read 2 to preserve read pairing, as the three runs represent the same sequencing library rather than independent replicates. Read counts were verified to be identical between mate files for every sample.

Assignment of GEO sample identifiers to sample names in the published count matrix was confirmed by matching the GEO SOFT sample titles to the selected samples, and Xist expression confirmed the annotated sex of each sample (female median 5,195 counts; male median 2 counts). Merged FASTQ files were verified for integrity prior to use.

Short reads were realigned to GRCm39 rather than reusing the processed output of the original study, which was aligned to GRCm38; splice junction coordinates are not interchangeable between genome builds, and conversion between assemblies would introduce error at the positions where precision is most critical. Alignment was performed by SQANTI3 internally using STAR, as described below.

**SQANTI3 quality control with short-read integration**

The merged transcriptome was independently evaluated using SQANTI3 (v5.5.4) as an orthogonal quality-control assessment complementing the primary gffcompare classification. SQANTI3 quality control was run with the merged transcriptome (merged_isoforms.gtf) as input, the GENCODE vM36 annotation as reference, and the GRCm39 primary assembly as reference genome. Transcription start site support at the 5′ end was assessed against the mouse refTSS v3.1 GRCm39 annotation, and polyA motif support at the 3′ end against the SQANTI3 mouse and human polyA motif list. Structural classification was run as a single chunk (-n 1) with open reading frame (ORF) prediction disabled (--skipORF), as coding potential was not evaluated in this study.

Short-read junction support was provided using the --short_reads option with a file of filenames listing the eight paired-end libraries described above. This option was selected rather than --SR_bam because calculation of minimum junction coverage requires the splice junction output that SQANTI3 generates by running STAR internally from raw FASTQ input. STAR (v2.7.11b) was used to build a genome index from the GRCm39 primary assembly without supplied splice junction annotation, and to align each short-read library once, yielding 1,487,193 splice junctions, of which 4,746 lacked strand information from STAR and were therefore assigned to both strands. Because the index was built without annotation guidance, short-read splice junctions were detected independently of the reference annotation against which the long-read transcriptome was classified. Isoform-level short-read expression was estimated using kallisto (v0.51.1) with a k-mer length of 31. Three transcripts lacking strand information were discarded during the correction stage, yielding 305,324 classified isoforms.

Each isoform was assigned a structural category (full-splice match, incomplete-splice match, novel in catalog, novel not in catalog, antisense, fusion, intergenic, genic, or genic intron) together with quality attributes including canonical splice junction status, annotated donor and acceptor site usage, minimum short-read junction coverage across all junctions of the isoform, cap analysis of gene expression (CAGE) peak support at the 5′ end, polyA motif support at the 3′ end, predicted reverse-transcriptase template switching (RT-switching), and the percentage of adenines in the genomic window downstream of the transcript termination site (used as an intra-priming indicator). A separate junction-level output table was generated reporting, for each splice junction, its novelty status, splice site dinucleotide, donor and acceptor site categories, distance to the nearest annotated splice site, and per-sample short-read coverage. The classification and junction tables are provided as Data S1 and Data S2.

#### QUANTIFICATION AND STATISTICAL ANALYSIS

##### Transcript detection and isoform diversity

All analyses in this section used the unfiltered isoform-level TPM matrix (305,327 transcripts), with structural annotation derived from the gffcompare annotated GTF. Transcripts assigned artifact-associated class codes were excluded from these analyses: i (transcript fully contained within a reference intron), e (single-exon transcript overlapping a reference intron, a possible pre-mRNA fragment), s (intron match on the opposite strand, a possible mapping error), and p (possible polymerase run-on). Transcripts were assigned to five biologically interpretable categories based on gffcompare class codes: Known (class code “=”), Novel Isoform (class code “j”), Novel Gene (class code “u”), Antisense (class code “x”), and Other (all remaining non-artifact class codes).

The number of transcripts detected per sample was defined as the count of transcripts with TPM ≥ 1 in that individual sample, computed independently for each of the 16 samples using the unfiltered TPM matrix. Transcript detection and classification were evaluated independently within each sex and within each sex-condition group (Female ALF, Female LRF, Male ALF, Male LRF). For within-sex analyses, a transcript was considered detected if it had TPM ≥ 1 in at least one sample belonging to that sex across all four zeitgeber time points and both feeding conditions. For within-group analyses, a transcript was considered detected if it had TPM ≥ 1 in at least one of the four samples belonging to that sex-condition group.

For isoform-per-gene quantification, only transcripts categorized as Known or Novel Isoform were retained, as these represent transcript models supported by reference annotation or novel splice junction evidence. Isoform counts were computed per gene using the reference gene name when available, or the StringTie gene identifier for novel loci. The number of distinct transcript isoforms per gene was counted, and genes were grouped into bins of 1, 2, 3, 4, or ≥ 5 isoforms. The top 30 genes ranked by total isoform count were identified separately for each sex and each sex-condition group and visualized as horizontal grouped bar charts showing Known and Novel Isoform transcripts separately.

**Clock gene expression profiles**

Temporal expression profiles of clock genes (*Bmal1*, *Clock*, *Per2*, *Per3*, *Cry1*, *Cry2*, *Nr1d1*, *Nr1d2*, *Dbp*) were extracted from the annotated gene-level count matrix and plotted against zeitgeber time for each sex and feeding condition. Values shown are gene-level counts; profiles are compared descriptively between feeding conditions within a sex rather than between sexes, and are not intended for quantitative cross-sample inference.

**Principal component analysis and sample correlation**

For principal component analysis (PCA) and pairwise correlation analyses, transcripts were retained if they had TPM ≥ 1 in at least two of the 16 samples, yielding a filtered matrix of 31,621 transcripts. Retained TPM values were $\log_2$-transformed as $\log_2(\text{TPM} + 1)$. Transcripts with zero variance across all samples were removed prior to PCA to prevent undefined scaling. PCA was performed using the prcomp function in R with unit-variance scaling (scale. = TRUE) on the transposed sample-by-transcript matrix, for all 16 samples combined and separately within female and male subsets. Group-level 95% confidence ellipses were drawn using stat_ellipse based on a multivariate normal distribution (level = 0.95).

Pairwise Pearson correlation coefficients were computed across all 16 samples from the same $\log_2$-transformed filtered matrix using the cor function in R. The resulting 16 × 16 correlation matrix was visualized as a heatmap using the pheatmap package. Hierarchical clustering was applied independently to both rows and columns using Euclidean distance and complete linkage. Sample annotations indicating sex, feeding condition, and group were included as colored annotation bars.

#### Gene-level differential expression analysis

Gene-level differential expression analysis was performed using the annotated gene count matrix. Prior to analysis, StringTie-derived MSTRG loci lacking valid genomic coordinate information (missing or empty chromosome annotation) were removed. This MSTRG artifact filter was applied identically in all downstream gene-level analyses. Gene identifiers were standardized by removing version suffixes from Ensembl identifiers prior to all comparison steps.

Gene expression filtering was performed in two sequential steps within each sex. First, a gene was considered expressed within a given sex-condition group if the sum of raw counts across all four samples in that group exceeded 5. Second, only genes expressed under both feeding conditions within a given sex were retained for differential expression testing; the intersection of expressed genes between ALF and LRF was computed within each sex, and only these shared genes entered DESeq2[6]. Genes expressed exclusively under one condition within a sex were excluded from differential testing and exported separately for condition-specific analyses.

Differential expression analysis was performed independently for female and male samples using DESeq2 (v1.46.0) under a negative binomial generalized linear model. The design formula ~ZT + Condition was applied to the shared gene count matrix for each sex, with ALF specified as the reference level for the Condition factor and zeitgeber time (ZT1, ZT7, ZT13, ZT19) modeled as a categorical covariate to account for time-of-day variation. Size factor normalization used the median-of-ratios method. Wald tests were used to estimate $\log_2$ fold changes for the LRF versus ALF contrast. Gene annotations were merged with DESeq2 results after deduplication by version-stripped gene identifier, retaining one annotation entry per gene. Missing adjusted p values were replaced with 1 and missing $\log_2$ fold change values with 0 prior to downstream classification and visualization. Because the two-stage expression filter described above removed low-expressed genes prior to testing, substantially reducing the number of hypotheses relative to the full annotation, unadjusted p values were used as the primary significance threshold at the gene level, and gene-level differential expression results are accordingly treated as exploratory and interpreted alongside the adjusted values. Differentially expressed genes were defined as $p < 0.01$ combined with an absolute $\log_2$ fold change $\geq 1$, and Benjamini-Hochberg adjusted p values were computed and exported for all tested genes.

Volcano plots were generated separately for each sex, with $\log_2$ fold change on the x-axis and $-\log_{10}$(p value) on the y-axis. For visualization only, $\log_2$ fold change values were truncated at the 99.5th percentile of absolute values to limit the influence of extreme outliers. The top 15 significant genes per direction were labeled, prioritizing annotated gene symbols over MSTRG-derived identifiers.

Sex-shared differentially expressed genes were identified by intersecting version-stripped gene identifiers between the female and male significant gene sets. $\text{Log}_2$ fold changes from both analyses were compared in scatter plots, with genes classified as concordant when regulation occurred in the same direction in both sexes and discordant when in opposite directions. All shared genes were labeled, ranked by the mean of their female and male p values.

**Condition-specific gene expression analysis**

Condition-specific gene expression analysis identified genes expressed exclusively under one feeding condition within each sex, independent of differential expression testing. The same MSTRG artifact filter, version suffix standardization, and expression criterion described above were applied, with gene expression defined separately for each sex-condition group. Within each sex, genes were classified as ALF-only, LRF-only, or shared, based on set difference and intersection operations on version-stripped gene identifiers. Cross-sex comparisons were performed by intersecting condition-specific gene sets between females and males to identify genes consistently ALF-specific or LRF-specific across both sexes.

**Transcript-level differential expression analysis**

Transcript-level differential expression analysis used the annotated transcript count matrix. Prior to analysis, MSTRG-derived transcripts lacking valid genomic coordinate information were removed using the same filtering logic applied at the gene level, and transcript identifiers were standardized by removing version suffixes.

Transcript expression filtering was performed in three sequential steps within each sex. First, a transcript was considered expressed within a given sex-condition group if the sum of raw counts across all four samples in that group exceeded 5. Second, only transcripts expressed under both feeding conditions within a given sex were retained, computed as the intersection of expressed transcripts between ALF and LRF within each sex. Third, artifact-class transcripts assigned gffcompare class codes i, e, s, or p were removed from the shared set prior to differential testing. Transcripts expressed exclusively under one condition were excluded from differential testing and exported separately for condition-specific analyses.

Differential expression analysis was performed independently for female and male samples using DESeq2 (v1.46.0) under a negative binomial generalized linear model with the design formula ~ZT + Condition, ALF as reference level, and zeitgeber time modeled as a categorical covariate. Size factor normalization used the median-of-ratios method, and Wald tests estimated $\log_2$ fold changes for the LRF versus ALF contrast. Transcript annotations were merged with DESeq2 results using transcript identifier as the key. Missing adjusted p values were replaced with 1, missing $\log_2$ fold change values with 0, and missing category labels with “Other”. Because a larger number of features was tested at the transcript level, Benjamini-Hochberg adjusted p values were used as the significance threshold. Differentially expressed transcripts were defined as adjusted $p < 0.05$ combined with an absolute $\log_2$ fold change $\geq 1$. Full result tables and size-factor-normalized counts were exported for all tested transcripts.

Each differentially expressed transcript was classified by its gffcompare isoform annotation category and summarized separately for each direction of change in each sex. To assess the relationship between transcript-level and gene-level differential expression, each transcript was classified based on the differential expression status of its parent gene. Parent gene identifiers were extracted from the ref_gene_id field after removing version suffixes and matched against the gene-level DESeq2 results. Transcripts were classified into three categories: concurrent gene-level differential expression when the parent gene was also significant at the gene level ($p < 0.01$ and $|\log_2$ fold change$| \geq 1$); isoform-specific regulation when the parent gene was tested but not significant at the gene level; and not tested when the parent gene was absent from the gene-level tested set or lacked a valid reference gene identifier.

Sex-shared differentially expressed transcripts were identified by intersecting transcript identifiers between the female and male significant sets, and $\log_2$ fold changes were compared in scatter plots with concordance classified as above. For expressed transcript Venn diagrams, expressed transcript sets were computed per sex-condition group as described above.

**Condition-specific transcript expression analysis**

Condition-specific transcript expression analysis identified transcripts detected exclusively under one feeding condition within each sex, independent of differential expression testing. The MSTRG artifact filter was applied, and artifact-class transcripts (class codes i, e, s, p) were excluded before computing expressed transcript sets. Transcript expression was defined separately for each sex-condition group using the same criterion described above, and transcripts were classified within each sex as ALF-only, LRF-only, or shared. Cross-sex comparisons were performed by intersecting condition-specific transcript sets between females and males.

The isoform category composition of each condition-specific transcript set was examined by classifying transcripts according to their gffcompare annotation category and computing the percentage in each category within each set. The parent gene status of each condition-specific transcript was determined by mapping the version-stripped ref_gene_id to gene-level expression detection sets derived independently from the gene count matrix, applying the same MSTRG filter and sum of counts > 5 criterion within each sex-condition group. Each parent gene was classified as gene ALF-only, gene LRF-only, gene shared, or gene not detected.

**Comparisons between sexes at gene and transcript level**

Comparisons between male and female ventricle were performed separately within each feeding condition using the depth-matched gene- and transcript-level count matrices described above, so that all libraries entering these models carried equal primary-read depth. Because male and female libraries were prepared and sequenced in separate batches, batch is fully confounded with sex in this design; depth matching equalizes sequencing depth but cannot separate technical batch effects from biological sex differences, and between-sex results are therefore interpreted as descriptive. The same MSTRG artifact filter and version suffix standardization applied to the within-sex analyses were applied here. Expression filtering used the same two-step procedure: a gene or transcript was considered expressed within a sex-condition group if the sum of raw counts across the four samples in that group exceeded 5, and only features expressed in both sexes within a given feeding condition were retained for testing.

Differential expression between sexes was tested using DESeq2 (v1.46.0) with the design formula ~ZT + Sex, with Female as the reference level and zeitgeber time modeled as a categorical covariate. Wald tests were used to estimate $\log_2$ fold changes for the Male versus Female contrast, performed independently for ALF and LRF samples. At the gene level, differentially expressed genes were defined as $p < 0.01$ combined with an absolute $\log_2$ fold change $\geq 1$. At the transcript level, differentially expressed transcripts were defined as Benjamini-Hochberg adjusted $p < 0.05$ combined with an absolute $\log_2$ fold change $\geq 1$. Volcano plots, category breakdowns by gffcompare structural class, condition-specific and shared feature comparisons, and heatmaps of the top 50 differentially expressed transcripts per condition were generated from these results. For transcript-level results, the differential expression status of each parent gene was determined by re-running the gene-level model on the same samples, allowing transcripts to be classified as showing concurrent gene-level differential expression or isoform-specific regulation.

Gene Ontology (GO) biological process enrichment was performed separately for each condition and direction (higher in males and higher in females), with the background universe defined as all gene symbols tested by DESeq2 for that condition, using the parameters described below.

**Differential transcript usage analysis**

Differential transcript usage (DTU) between ALF and LRF was assessed independently for female and male samples using DRIMSeq[7] (v1.34.0), which models isoform proportion changes using a Dirichlet-multinomial framework[8]. The annotated transcript count matrix was used as input, with transcripts mapped to their parent genes using the ref_gene_id field. Prior to modeling, genes with fewer than two expressed isoforms were excluded from each sex-specific dataset. Transcript count matrices were then filtered using dmFilter with the following criteria: total gene expression above 10 counts in all samples within that sex (min_gene_expr = 10, min_samps_gene_expr = n_total); each transcript was required to have counts of at least 10 in a minimum number of samples equal to the smaller condition group size (min_feature_expr = 10, min_samps_feature_expr = n_small); and each transcript was required to contribute a minimum proportion of 0.1 of its parent gene's total expression in at least that same number of samples (min_feature_prop = 0.1, min_samps_feature_prop = n_small)[8].

The DRIMSeq model was specified using the design matrix ~condition + ZT, with ALF as the reference condition and zeitgeber time modeled as a blocking factor to control for time-of-day variation. A random seed of 42 was set prior to precision estimation. Precision parameters were estimated using dmPrecision with a random subset of 50% of genes (prec_subset = 0.5) and common moderation (prec_moderation = “common”). Model fitting used dmFit, and hypothesis testing used dmTest with the conditionLRF coefficient as the test contrast. Gene-level and transcript-level results were extracted separately. Missing adjusted p values at both levels were replaced with 1 prior to downstream classification. Genes were classified as showing significant DTU if the gene-level false discovery rate (FDR)-adjusted p value was below 0.05. Precision diagnostics were exported for each sex.

For each DTU gene, mean isoform proportions were calculated per condition by dividing individual transcript counts by the total gene count across all transcripts within that gene, computed separately for ALF and LRF samples. The delta isoform proportion (Δ) was defined as the mean LRF proportion minus the mean ALF proportion for each transcript. The maximum absolute Δ per gene was used to quantify the magnitude of isoform usage change and visualized as violin plots comparing all tested genes against significant DTU genes. DTU genes were classified into three switch categories: Major Switch, genes in which the dominant isoform changes between ALF and LRF with maximum absolute $\Delta \geq 0.15$; Partial Redistribution, genes in which the dominant isoform remains the same but maximum absolute $\Delta \geq 0.08$; and Minor Shift, all remaining DTU genes.

Sex-specific and shared DTU genes were identified by comparing significant gene lists between female and male analyses. For genes shared between sexes, the dominant isoform was defined as the isoform with the highest proportion across either condition, selected using the maximum of the mean ALF and mean LRF proportions. The delta isoform proportion of this dominant isoform was extracted independently for each sex and compared in a scatter plot to assess concordance of switching direction.

The overlap between DTU genes and gene-level differentially expressed genes was assessed by cross-referencing significant DTU gene identifiers against gene-level DESeq2 results using the same significance thresholds applied in the gene-level analysis ($p < 0.01$ and $|\log_2 \text{fold change}| \geq 1$). Version suffixes were stripped from Ensembl gene identifiers prior to matching. The proportion of DTU genes also showing gene-level differential expression was visualized as stacked bar charts, and Fisher's exact test was used to assess whether DTU genes were enriched for concurrent gene-level differential expression relative to all DRIMSeq-tested genes, with the background restricted to genes passing dmFilter regardless of DTU significance.

Representative isoform proportion plots were generated for the top four DTU genes per sex, ranked first by switch class and then by FDR-adjusted p value within each class; for the male gene set, genes already selected for the female panel were excluded to ensure the two panels display non-overlapping genes. Stacked bar charts showed mean isoform proportions under ALF and LRF, with isoforms ranked by peak expression across either condition and annotated by structural category derived from gffcompare class codes. The maximum absolute isoform proportion change across all isoforms and the gene-level FDR significance were annotated above each plot. Extended isoform proportion plots were generated for curated sets of biologically relevant cardiac genes showing significant DTU, organized by functional category including transcription and developmental regulators, stress and signaling, mitochondrial metabolism and translation, RNA splicing and transcript regulation, circadian rhythmicity, sarcomere organization, and ion channel and mTOR signaling. These were visualized separately for female-specific and male-specific DTU genes. For shared DTU genes, side-by-side female and male proportion plots were generated. All isoform proportion data, structural annotations, switch classifications, and significance values were exported as annotated CSV files.

Between-sex differential transcript usage was assessed independently within each feeding condition using DRIMSeq with the design formula ~sex + ZT, with Female as the reference level for the Sex factor. The same dmFilter criteria and precision estimation parameters described above for within-sex DTU were applied. Prior to modeling, genes containing any isoform with zero total counts across all samples of one sex were excluded to prevent non-convergence of the Dirichlet-multinomial precision estimates. Gene-level and transcript-level results were extracted as above, and genes with FDR-adjusted $p < 0.05$ were classified as significant. Switch classification, DTU–differential expression overlap, and GO enrichment followed the same procedures described for the within-sex analyses.

**Splicing architecture of differential transcript usage genes**

Splicing event annotation was performed using IsoformSwitchAnalyzeR[9] (v2.6.0) independently for females and males. The merged assembled transcriptome annotation (gffcompare annotated GTF) was used as the isoform exon annotation input, enabling structural event classification across both reference-matched and novel isoforms. The transcript count matrix was filtered using preFilter with a gene expression cutoff of 1, an isoform expression cutoff of 0, and with single-isoform genes removed. A random seed of 42 was set prior to analysis. Eight structural event types were annotated per transcript: exon skipping, multi-exon skipping, mutually exclusive exons, intron retention, alternative 5′ splice site, alternative 3′ splice site, alternative transcription start site, and alternative transcription termination site. The analyzeAlternativeSplicing function was applied with alpha = 0.05, dIFcutoff = 0.08, and onlySwitchingGenes = FALSE to extract structural event flags for all expressed isoforms regardless of switching significance. Where the annotation step required a prior switch test, DEXSeq was run as a fallback using isoformSwitchTestDEXSeq with results treated as informational only, and the DRIMSeq FDR gate was never overridden.

All downstream analyses were restricted to genes with significant DTU (DRIMSeq FDR < 0.05). For each DTU gene, a switch pair of isoforms was defined per sex. For genes classified as a Major Switch (the dominant isoform differs between ALF and LRF), the switch pair was the isoform dominant under ALF and the isoform dominant under LRF, the identity-flip pair that defines the switch. For genes classified as Partial Redistribution, and for Minor Shift genes in which the dominant isoform did not change, the switch pair was instead the isoform with the largest positive change in proportion (most LRF-gaining) and the isoform with the largest negative change (most LRF-losing). Gene-level event flags were defined as the union of both sides of the switch pair, such that a gene was considered to carry an event if either isoform did. Female and male splicing architecture analyses were performed independently, and no statistical comparison was made between sexes.

The top 20 DTU genes per sex ranked by maximum absolute isoform proportion shift were visualized as dumbbell plots showing mean ALF and mean LRF proportions, with arrows indicating direction of change and a colored strip indicating switch magnitude class. For each sex, the number and percentage of DTU genes carrying each of the eight event types were counted using the union flags defined above.

Directional enrichment of each event type at the switch was assessed per DTU gene by classifying the event as gained (present on the LRF-side isoform of the switch pair only), lost (present on the ALF-side isoform only), or concordant (present on both isoforms or neither). For each event type, an exact binomial test was performed on the discordant (gained-only vs. lost-only) gene counts, analogous to McNemar's test for paired binary data, testing the null hypothesis that an event is equally likely to be gained or lost at the switch. A Haldane-corrected odds ratio was calculated as (n gained + 0.5) / (n lost + 0.5) to avoid undefined values when one discordant count was zero, with a 95% confidence interval obtained by transforming the exact binomial confidence interval on the proportion gained to the odds-ratio scale. Benjamini-Hochberg correction was applied within each sex across all eight event types. Event type prevalence across switch magnitude classes (Major Switch, Partial Redistribution, Minor Shift) was visualized as a bubble plot using the union flags defined above, and was formally compared between Major Switch and Partial Redistribution genes for each event type using Fisher's exact test on isoform-level 2×2 contingency tables, with Benjamini-Hochberg correction applied as above. Minor Shift genes were visualized but excluded from this statistical comparison due to small sample size in some categories.

Co-occurrence of splicing event types was assessed by computing the Jaccard index for each pair of event types, defined as the number of DTU genes carrying both events divided by the number carrying either event. Fisher's exact test was applied independently to each off-diagonal pair to test for non-random co-occurrence, with

Benjamini-Hochberg correction applied within each sex across all off-diagonal pairs. Only event types present in at least three DTU genes were included.

**Gene Ontology enrichment analysis**

GO biological process enrichment analysis was performed using clusterProfiler[10] (v4.14.6) with the org.Mm.eg.db annotation database (v3.20.0). Gene symbols were converted to Entrez identifiers using the bitr function. Enrichment was performed using enrichGO with the Biological Process ontology, Benjamini-Hochberg multiple testing correction, and pvalueCutoff and qvalueCutoff both set to 1 to retrieve all terms, followed by filtering on p value < 0.05. Analyses were performed only when at least five genes with valid symbols were available for a given gene set. The top 10 enriched terms per gene set were selected by p value and visualized as bubble plots, with bubble size representing gene count and the x-axis representing $-\log_{10}$(p value).

Background universes were defined to match each analysis. For gene-level differential expression, enrichment was performed separately for each sex and direction of change (higher under LRF and higher under ALF), with the background defined as all gene symbols from the DESeq2-tested gene set for that sex with non-missing annotations. For within-sex condition-specific analyses, the background was the union of ALF-only and LRF-only genes for that sex. For cross-sex condition-specific analyses, the background was the union of all ALF-specific genes across both sexes for ALF enrichment, and the union of all LRF-specific genes across both sexes for LRF enrichment. For transcript-level condition-specific analyses, gene symbols were extracted from the gene_name field of the transcript annotation table and equivalent background definitions were applied. For DTU analyses, the background was restricted to all genes passing dmFilter regardless of significance, rather than the full annotation database, to prevent inflated enrichment estimates.

##### Short-read junction support and splice site analysis

Short-read support for long-read transcript models was tabulated from the SQANTI3 classification output by structural category. For each isoform, the minimum short-read junction coverage across all of its splice junctions was used, and the percentage of isoforms with support was computed at thresholds of at least one, three, and five short reads. Support rates were reported separately for the full assembled catalog and for the expressed subset, defined as transcripts with TPM ≥ 1 in at least two of the 16 long-read samples. Because the short-read reference libraries were generated from separate animals maintained under constant darkness, junction coverage was interpreted as supporting rather than defining evidence.

Splice junctions of isoforms containing at least one unannotated splice site were characterized using the SQANTI3 junction-level output. For each junction classified as novel, the donor and acceptor site categories, the splice site dinucleotide, and the absolute distance from each novel site to the nearest annotated splice site of the same type were extracted. The proportion of novel junctions pairing an annotated donor with a novel acceptor was computed, together with the median and cumulative distribution of distances from novel acceptor sites to the nearest annotated acceptor.

##### Statistical reporting

All statistical analyses and visualizations were performed in R (v4.4.2) using RStudio (v2024.09.1+394). Statistical details for each analysis, including the test used, the definition of significance thresholds, and the number of genes or transcripts included, are provided in the relevant subsections above and in the corresponding figure legends. Comparisons between feeding conditions were performed independently within each sex, and comparisons between sexes were performed on depth-matched count matrices within each feeding condition. Each sex-condition group comprised four animals, one at each zeitgeber time point, with no within-time-point replication; N = 4 biological replicates per sex per feeding condition distributed across the 24 h cycle. Where random sampling or stochastic estimation was involved, a fixed seed of 42 was used to ensure reproducibility.

**Figure S1. Nanopore long-read RNA sequencing quality metrics and transcript structural annotation, related to Figure 2**

(A) Total reads per sample (millions).

(B) Mean read length (bp) per sample.

(C) Read length N50 (bp) per sample (the read length above which 50% of total sequenced bases are contained).

(D) Primary aligned reads per sample (millions).

(E) Percentage of supplementary (split) alignments per sample.

(F) Distribution of gffcompare structural class codes across the merged transcriptome. Artifact class codes (i, e, s, p) were excluded (90 transcripts removed, <1%). Categories: charcoal, Known; magenta, Novel Isoform; pink, Novel Gene; green, Antisense; gray, Other. Transcript counts are annotated above each bar.

(G) Number of detected transcripts (TPM ≥ 1) per sample.

In (A–E) and (G), each bar represents one sample, grouped by sex. Light orange, ALF females; dark orange, ALF males; light teal, LRF females; dark teal, LRF males. Read-level metrics in (A–C) were obtained with NanoStat (v1.6.0); alignment metrics in (D) and (E) with samtools flagstat and samtools stats (v1.12). Structural classification in (F) was performed using gffcompare (v0.12.10) against the GENCODE vM36 mouse annotation. Transcript detection in (G) was computed from the unfiltered transcript-level TPM matrix (305,327 transcripts).

**Figure S2. Gene body coverage profiles across all samples, related to Figure 2**

Gene body coverage for all 16 samples across 5,000 randomly selected transcripts (summed exonic length ≥ 1 kb) from the GENCODE vM36 annotation. The x-axis shows normalized gene body position from 5′ to 3′; the y-axis shows normalized coverage. Blue, males; red, females. Solid lines, ALF; dashed lines, LRF. Each line represents one sample. Coverage was computed using geneBody_coverage.py from the RSeQC package (v5.0.4).

**Figure S3. Pairwise sample correlation heatmap, related to Figure 2**

Heatmap of pairwise Pearson correlation coefficients across all 16 samples. Transcript-level TPM values were filtered (TPM ≥ 1 in at least 2 of 16 samples; 31,621 transcripts retained) and $log_2$-transformed as $log_2$(TPM + 1). Color scale: orange, high correlation; teal, low correlation. Hierarchical clustering was applied independently to rows and columns using Euclidean distance and complete linkage. Colored annotation bars indicate group (light orange, ALF females; dark orange, ALF males; light teal, LRF females; dark teal, LRF males), condition (yellow, ALF; blue, LRF), and sex (pink, females; blue, males).

**Figure S4. Comparison of SQANTI3 and gffcompare structural classifications, related to Figure 2**

Side-by-side classification of all assembled transcripts by SQANTI3 (v5.5.4; left) and gffcompare (v0.12.10; right). Three transcripts lacking strand information were discarded during the SQANTI3 correction stage, yielding 305,324 classified transcript isoforms in both tables. These three transcripts were classified as gffcompare class "u" (Novel Gene) in the unfiltered assembly (Figure S1), accounting for the small difference in "u" counts between the two classifications (1,473 vs. 1,470). SQANTI3 structural categories include full-splice match (FSM), incomplete-splice match (ISM), novel in catalog (NIC), novel not in catalog (NNC), antisense, fusion, intergenic, genic intron, and genic. gffcompare class codes are listed with structural descriptions. Counts and percentages are shown for each category. Both classifications were derived from the same merged transcriptome assembly compared against the GENCODE vM36 mouse annotation.

**Figure S5. SQANTI3 quality-control attributes by structural category, related to Figure 2**

SQANTI3 quality attributes for the four major structural categories: FSM (blue), ISM (orange), NIC (green), and NNC (yellow). Top left, transcript isoform count per category on a $\log_{10}$ scale (full catalog, n = 305,324). Top right, percentage of transcript isoforms with canonical splice site dinucleotides at all junctions. Middle left, percentage of transcript isoforms with annotated donor and acceptor splice sites. Middle right, percentage of transcript isoforms with a polyA motif detected at the 3′ end. Bottom left, percentage of transcript isoforms flagged as predicted reverse transcriptase (RT)-switching artifacts. Bottom right, percentage of transcript isoforms with cap analysis of gene expression (CAGE) peak support at the 5′ end, assessed against the mouse refTSS v3.1 annotation. Values are annotated above each bar.

**Figure S6. Short-read junction support for long-read transcript models, related to Figure 2**

Percentage of transcript isoforms with short-read splice junction support by SQANTI3 structural category, evaluated at three minimum coverage thresholds: ≥ 1 read (blue), ≥ 3 reads (orange), and ≥ 5 reads (green). Top, all transcript isoforms in the full merged catalog. Bottom, expressed transcript isoforms only (TPM ≥ 1 in at least 2 of 16 long-read samples). Sample sizes per category are shown below each group. Short-read support was derived from eight publicly available paired-end RNA sequencing libraries from mouse ventricle (GEO: GSE262714; Zhang et al., 2024), realigned to GRCm39 with STAR (v2.7.11b) independently of the reference annotation. For each transcript isoform, the minimum short-read junction coverage across all of its splice junctions was used. Values are annotated above each bar.

**Figure S7. Transcript structural classification, transcript isoform diversity, and transcript isoform-rich genes by sex and feeding condition, related to Figure 2**

Panels are organized in four rows, one per sex-by-condition group: Female ALF (A–C), Female LRF (D–F), Male ALF (G–I), and Male LRF (J–L).

(A, D, G, J) Distribution of detected transcripts by gffcompare structural category. Categories: charcoal, Known; magenta, Novel Isoform; pink, Novel Gene; green, Antisense; gray, Other. Transcript counts and percentages are annotated above each bar. A transcript was considered detected within a sex-by-condition group if TPM ≥ 1 in at least one of the four samples in that group.

(B, E, H, K) Number of genes grouped by transcript isoform count (1, 2, 3, 4, or ≥ 5 detected transcript isoforms per gene). Only Known ("=") and Novel Isoform ("j") transcripts were included.

(C, F, I, L) Top 30 genes ranked by total transcript isoform count within each sex-by-condition group. Horizontal stacked bars show the number of Known (charcoal) and Novel Isoform (magenta) transcripts per gene.

**Figure S8. Condition-specific gene expression under ad libitum and light-cycle time-restricted feeding, related to Figure 3**

(A and B) Venn diagrams of genes expressed under ALF or LRF within females (A) and males (B). A gene was considered expressed within a sex-by-condition group if the sum of raw counts across all four samples exceeded 5. Orange, ALF; teal, LRF. Counts and percentages for ALF-only, shared, and LRF-only genes are shown.

(C and D) GO biological process enrichment for condition-specific genes in females (C) and males (D). Top 10 enriched terms per direction ($p < 0.05$) for ALF-specific genes (left column; light orange in females, dark orange in males) and LRF-specific genes (right column; light teal in females, dark teal in males). Dot size is proportional to gene count. The x-axis shows $-\log_{10}$(p-value). Background universe: the union of ALF-only and LRF-only genes within each sex.

(E) Four-way Venn diagram showing overlap of condition-specific gene sets across sexes. Each circle represents the condition-specific genes for one sex-by-condition group (Female ALF-only, Female LRF-only, Male ALF-only, Male LRF-only). ALF-specific genes shared across both sexes: 137 (1.8%); LRF-specific genes shared across both sexes: 97 (1.3%). Counts and percentages are shown for all intersections.

(F and G) GO biological process enrichment for condition-specific genes shared across both sexes. (F) ALF-specific shared genes (n = 137); background: all ALF-specific genes across both sexes. (G) LRF-specific shared genes (n = 97); background: all LRF-specific genes across both sexes. The remaining intersections (135 and 115 genes) represent condition-specific sets that differ in feeding condition between sexes. Top 10 enriched terms ($p < 0.05$); dot size and color are proportional to gene count. The x-axis shows $-\log_{10}$(p-value).

**Figure S9. Between-sex differential gene expression using depth-matched data, related to Figure 3**

(A and B) Venn diagrams of genes expressed in females and males under ALF (A) and LRF (B). A gene was considered expressed within a sex-by-condition group if the sum of raw counts across all four depth-matched samples exceeded 5. Pink, females; blue, males. Counts and percentages for female-only, shared, and male-only expressed genes are shown.

(C and D) Volcano plots of gene-level DE between sexes under ALF (C) and LRF (D). The x-axis shows $\log_2$ fold change (Male/Female); the y-axis shows $-\log_{10}$(p-value). Dashed lines indicate significance thresholds ($p < 0.01$ and $|\log_2 FC| \geq 1$). Pink, significantly higher in females; blue, significantly higher in males; gray, not significant. The top 15 genes per direction are labeled. DEG counts per direction are annotated: ALF, 2,803 up in females and 2,616 up in males; LRF, 2,779 up in females and 2,704 up in males.

(E and F) GO biological process enrichment for between-sex DEGs under ALF (E) and LRF (F). Top 10 enriched terms per direction ($p < 0.05$) for genes higher in females (left column; pink) and genes higher in males (right column; blue). Dot size is proportional to gene count. The x-axis shows $-\log_{10}$(p-value). Background universe: all DESeq2-tested genes within each feeding condition.

(G) Venn diagram of between-sex DEGs identified under ALF and LRF. Orange, ALF-specific DEGs (1,550; 22.0%); teal, LRF-specific DEGs (1,614; 22.9%); overlap, shared DEGs (3,869; 55.0%).

(H) Scatter plot of $\log_2$ fold change (Male/Female) for the 3,869 shared between-sex DEGs under ALF (x-axis) versus LRF (y-axis). Pink, concordant higher in females in both conditions; blue, concordant higher in males in both conditions; gray, discordant direction between conditions. The top 15 genes per direction are labeled by fold change magnitude. Dashed lines indicate $|\log_2 FC| \geq 1$.

**Figure S10. Condition-specific transcript expression across sexes, related to Figure 4**

(A and B) Venn diagrams showing the overlap between transcripts expressed under ALF and LRF within females (A) and males (B). Transcript expression was defined as row sum > 5 across the four samples per sex-by-condition group after exclusion of artifact-class transcripts (gffcompare class codes i, e, s, p). Orange circles represent ALF-expressed transcripts; teal circles represent LRF-expressed transcripts. In females, 7,888 transcripts (18.5%) were ALF-specific, 7,790 (18.3%) were LRF-specific, and 26,920 (63.2%) were shared. In males, 11,231 transcripts (16.5%) were ALF-specific, 9,746 (14.3%) were LRF-specific, and 47,273 (69.3%) were shared.

(C and D) GO biological process enrichment for condition-specific transcripts in females (C) and males (D). The top 10 enriched terms ($p < 0.05$) are shown for each direction (ALF-specific and LRF-specific) within each sex. Dot size is proportional to gene count. The x-axis shows $-\log_{10}$(p-value). Background universe: the union of ALF-only and LRF-only transcript-derived gene symbols within each sex.

(E) Four-way Venn diagram showing the intersection of condition-specific transcripts across sexes. ALF-specific transcripts shared between both sexes: 731; LRF-specific transcripts shared between both sexes: 625. The remaining intersections (725 and 649 transcripts) represent condition-specific sets that differ in feeding condition between sexes.

(F and G) GO biological process enrichment for condition-specific transcripts shared across both sexes for ALF (F; n = 731 transcripts) and LRF (G; n = 625 transcripts). The top 10 enriched terms ($p < 0.05$) are shown. Background universe: the union of all ALF-specific transcripts across both sexes for ALF enrichment (F); the union of all LRF-specific transcripts across both sexes for LRF enrichment (G).

(H) Transcript isoform category composition of condition-specific transcripts. Stacked bar plots show the percentage of transcripts in each gffcompare annotation category for ALF-specific and LRF-specific transcript sets within each sex. Known transcripts (charcoal) comprise 77–85% of each set; Novel Isoform transcripts (magenta) comprise 13–19%. Remaining categories include Novel Gene, Antisense, and Other (additional colors as shown).

(I) Parent gene status of condition-specific transcripts. Stacked bar plots show the percentage of transcripts whose parent gene was classified as Gene ALF-only (orange), Gene LRF-only (teal), Gene shared (green), or Gene not detected (gray), based on independent gene-level expression detection (row sum > 5) within each sex-by-condition group. Across all sets, 72–79% of condition-specific transcripts derive from genes expressed under both feeding conditions.

**Figure S11. Between-sex differential transcript expression under each feeding condition, related to Figure 4**

(A and B) Venn diagrams showing the overlap between transcripts expressed in females and males under ALF (A) and LRF (B). Transcript expression was defined as row sum > 5 across the four samples per sex-by-condition group using the depth-matched transcript count matrix, after exclusion of artifact-class transcripts (gffcompare class codes i, e, s, p). Pink circles represent female-expressed transcripts; blue circles represent male-expressed transcripts. Under ALF, 9,660 transcripts were female-specific, 15,581 were male-specific, and 23,756 were shared. Under LRF, 9,708 were female-specific, 15,276 were male-specific, and 23,225 were shared.

(C and D) Volcano plots showing differential transcript expression between sexes under ALF (C) and LRF (D). Only transcripts expressed in both sexes within each condition were retained for differential testing. Each point represents one transcript. The x-axis shows $\log_2$FC (Male/Female); the y-axis shows $-\log_{10}$(adjusted p-value). Horizontal dashed line indicates padj = 0.05; vertical dashed lines indicate $|\log_2\text{FC}| = 1$. Pink, transcripts significantly higher in females ($\log_2\text{FC} < -1$, padj < 0.05); blue, transcripts significantly higher in males ($\log_2\text{FC} > 1$, padj < 0.05); gray, not significant. Under ALF, 4,837 DETs were higher in females and 3,650 were higher in males. Under LRF, 4,833 were higher in females and 3,681 were higher in males.

(E) DETs classified by gffcompare transcript isoform annotation category and direction of change. Diverging bar plots show the number of DETs in each category, with transcripts higher in males (positive direction) above the axis and transcripts higher in females (negative direction) below. Bars are colored by transcript isoform category

(Known, Novel Isoform, Novel Gene, Antisense, Other). Results are shown separately for ALF and LRF.

(F) Venn diagram comparing DETs identified under ALF and LRF. ALF-specific DETs: 3,676; shared DETs: 4,811; LRF-specific DETs: 3,703.

(G) Scatter plot of $log_2FC$ values for the 4,811 DETs shared between ALF and LRF. Each point represents one transcript, with ALF $log_2FC$ on the x-axis and LRF $log_2FC$ on the y-axis. Pink, higher in females under both conditions; blue, higher in males under both conditions; gray, discordant direction between conditions.

(H) Transcript isoform-specific versus gene-level expression between sexes. Stacked bar plots show the proportion of DETs classified as showing concurrent gene-level DE (Tx, transcript; Tx DE and Gene DE; red), transcript isoform-specific regulation (Tx DE, Gene NOT DE; blue), or parent gene not tested (gray). Gene-level DE was assessed using the depth-matched gene count matrix and defined as $p < 0.01$ and $|log_2FC| \geq 1$. Results are shown separately for ALF and LRF, stratified by direction.

**Figure S12. Expression heatmaps of top differentially expressed transcripts between feeding conditions within each sex, related to Figure 4**

(A and B) Heatmaps showing z-scored expression of the top 50 DETs (ranked by adjusted p-value) between ALF and LRF within females (A) and males (B). Rows represent individual transcripts (labeled by gene symbol); columns represent individual samples. Expression values were z-score normalized across samples within each sex. Color scale ranges from purple (−2) through white (0) to red (+2). Rows are hierarchically clustered by Euclidean distance. Top annotation bars indicate ZT (ZT1, light yellow; ZT7, dark yellow; ZT13, saffron; ZT19, dark red) and Condition (ALF, orange; LRF, teal).

**Figure S13. Expression heatmaps of top differentially expressed transcripts between sexes within each feeding condition, related to Figure 4**

(A and B) Heatmaps showing z-scored expression of the top 50 DETs (ranked by adjusted p-value) between females and males under ALF (A) and LRF (B). Rows represent individual transcripts (labeled by gene symbol); columns represent individual samples from both sexes within each feeding condition. Expression values were z-score normalized across samples within each condition. Color scale ranges from purple (−1) through white (0) to red (+1). Rows are hierarchically clustered by Euclidean distance. Top annotation bars indicate ZT (ZT1, light yellow; ZT7, dark yellow; ZT13, saffron; ZT19, dark red) and Sex (Female, pink; Male, blue).

**Figure S14. Characterization of differential transcript usage between sexes within each feeding condition, related to Figure 5**

(A) Bar plots showing the number of genes with significant DTU between sexes (gene-level BH-FDR < 0.05) identified independently under ALF (177 genes) and LRF (166 genes).

(B) Violin plots showing the distribution of Δmax (maximum absolute change in transcript isoform proportion between males and females) for DTU genes under ALF and LRF. Horizontal dashed lines indicate the thresholds for Major Switch (Δmax ≥ 0.15) and Partial Redistribution (Δmax ≥ 0.08).

(C) Stacked bar plots showing the classification of DTU genes into three switch categories per feeding condition. Under ALF, 91 genes (52%) were classified as Major

Switch (red), 76 (43%) as Partial Redistribution (orange), and the remainder as Minor Shift (gray). Under LRF, 84 (51%) were Major Switch, 76 (46%) were Partial Redistribution, and the remainder were Minor Shift.

(D) Venn diagram showing the overlap between ALF and LRF DTU gene sets. ALF-specific DTU genes: 136 (45.0%); shared between feeding conditions: 41 (13.6%); LRF-specific DTU genes: 125 (41.4%).

(E) Scatter plot comparing the dominant transcript isoform Δ proportion under ALF (x-axis) and LRF (y-axis) for the 41 DTU genes shared between feeding conditions. The dominant transcript isoform was defined as the transcript isoform with the highest mean proportion across either sex. Green points indicate concordant direction of sex-difference between feeding conditions (31 genes); red points indicate discordant direction (10 genes).

(F) Stacked bar plots showing the overlap between DTU genes and DEGs per feeding condition. Maroon: genes showing both DTU and gene-level DE (DEG+DTU; expression + switching); teal: genes showing DTU without concurrent gene-level DE (DTU only; no expression change). Under ALF, 98 genes (55%) showed concurrent DEG+DTU and 79 (45%) showed DTU only. Under LRF, 93 (56%) showed DEG+DTU and 73 (44%) showed DTU only. OR and p-values from Fisher's exact test are annotated (ALF: OR = 1.46, $p = 0.051$; LRF: OR = 1.33, $p = 0.15$). Gene-level DE thresholds: $p < 0.01$ and $|\log_2 FC| \geq 1$. Background for Fisher's test: all genes passing DRIMSeq dmFilter regardless of DTU significance.

(G and H) Transcript isoform proportion plots for the top four DTU genes ranked by switch class and then by BH-FDR under ALF (G) and LRF (H). Stacked bars show mean transcript isoform proportions for females (left) and males (right). The switch class, gene-level BH-FDR, Δmax value, and significance level (asterisks) are annotated above each plot.

(I and J) GO biological process enrichment for DTU genes under ALF (I) and LRF (J). The top 10 enriched terms ($p < 0.05$) are shown. Dot size is proportional to gene count. The x-axis shows $-\log_{10}$(p-value). Background universe: all genes passing DRIMSeq dmFilter regardless of DTU significance.

**Figure S15. Transcript isoform proportion shifts in shared DTU genes between females and males, related to Figure 5**

Transcript isoform proportion plots for selected DTU genes identified in both females and males (shared DTU genes), showing sex-specific differences in transcript isoform switching patterns. Each row displays one gene with the female result (left) and male result (right) shown side by side. Stacked bars show mean transcript isoform proportions under ALF and LRF. The switch class, gene-level BH-FDR, Δmax value, and significance level (asterisks) are annotated for each sex independently. Genes shown: *Ptgds*, *Kremen1*, *Tfpi*, *Tubgcp4*, *Phc1*, *Adrm1*, and *Nrap*. Several genes exhibit different switch classifications between sexes (e.g., Partial Redistribution in one sex and Major Switch in the other), reflecting sex-specific differences in the magnitude or pattern of transcript isoform usage change.

**Figure S16. Transcript isoform proportion plots for female cardiac DTU genes, related to Figure 5**

Extended transcript isoform proportion plots for a curated set of biologically relevant cardiac genes showing significant DTU in females. Each subplot displays one gene, with stacked bars showing mean transcript isoform proportions under ALF (left) and

LRF (right). Genes are organized by functional category including transcription and developmental regulators, stress and signaling, mitochondrial metabolism and translation, RNA splicing and transcript regulation, circadian rhythmicity, sarcomere organization, and ion channel and mTOR signaling.

**Figure S17. Transcript isoform proportion plots for male cardiac DTU genes, related to Figure 5**

Extended transcript isoform proportion plots for a curated set of biologically relevant cardiac genes showing significant DTU in males. Each subplot displays one gene, with stacked bars showing mean transcript isoform proportions under ALF (left) and LRF (right). Genes are organized by functional category including transcription and developmental regulators, stress and signaling, mitochondrial metabolism and translation, RNA splicing and transcript regulation, circadian rhythmicity, sarcomere organization, and ion channel and mTOR signaling.

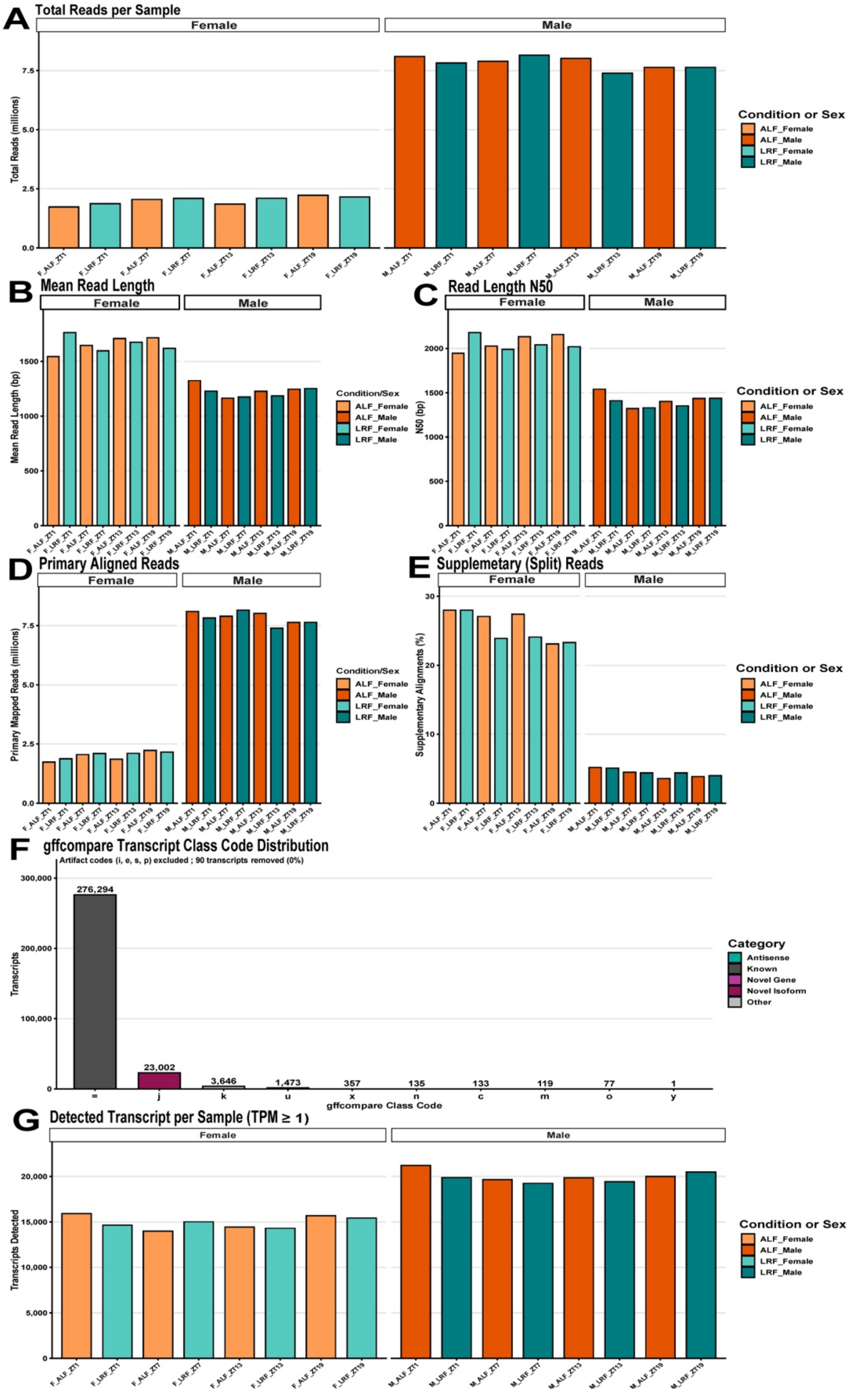
A
Total Reads per Sample
Female
Male
Total Reads (millions)
Condition or Sex
ALF_Female
ALF_Male
LRF_Female
LRF_Male
B
Mean Read Length
Mean Read Length (bp)
Condition/Sex
C
Read Length N50
N50 (bp)
D
Primary Aligned Reads
Primary Mapped Reads (millions)
E
Supplemetary (Split) Reads
Supplementary Alignments (%)
F
gffcompare Transcript Class Code Distribution
Artifact codes (i, e, s, p) excluded ; 90 transcripts removed (0%)
Transcripts
gffcompare Class Code
276,294
23,002
3,646
1,473
357
135
133
119
77
1
Category
Antisense
Known
Novel Gene
Novel Isoform
Other
G
Detected Transcript per Sample (TPM ≥ 1)
Transcripts Detected

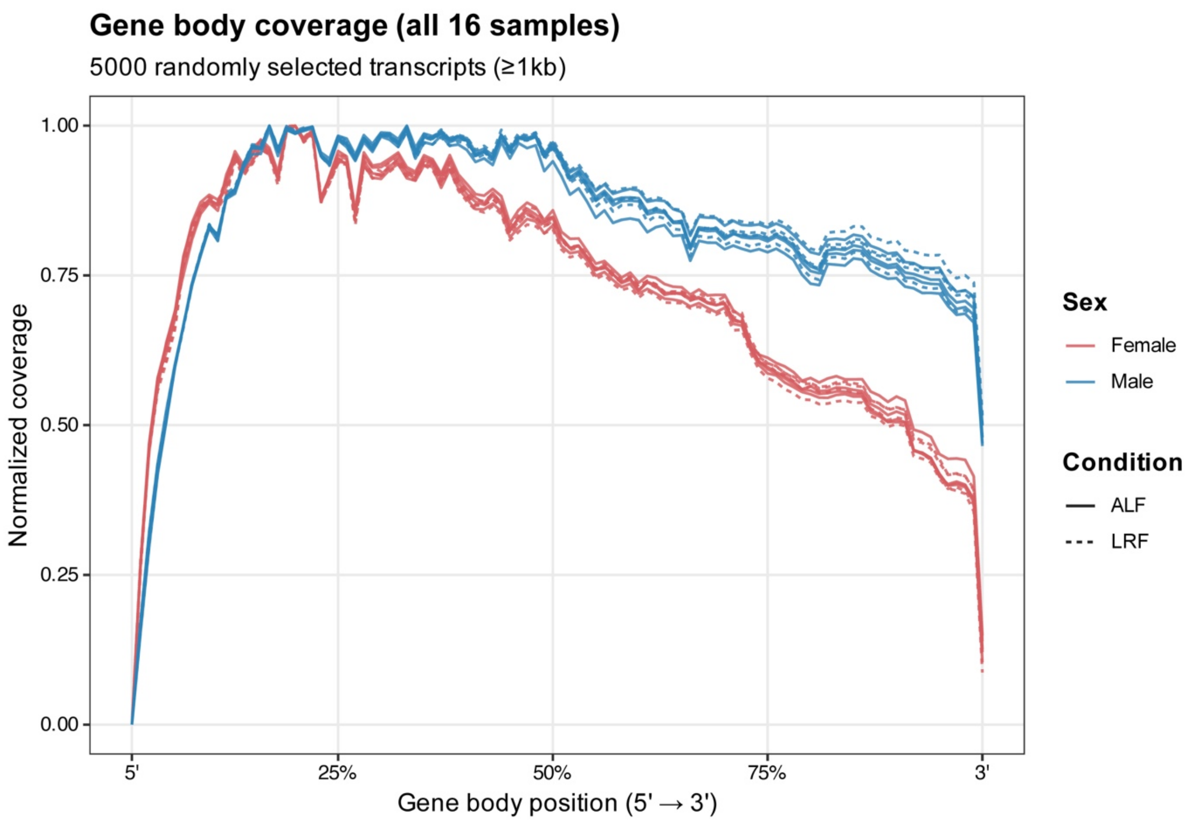
Gene body coverage (all 16 samples)
5000 randomly selected transcripts (≥1kb)
Normalized coverage
1.00
0.75
0.50
0.25
0.00
5'
25%
50%
75%
3'
Gene body position (5' → 3')
Sex
Female
Male
Condition
ALF
LRF

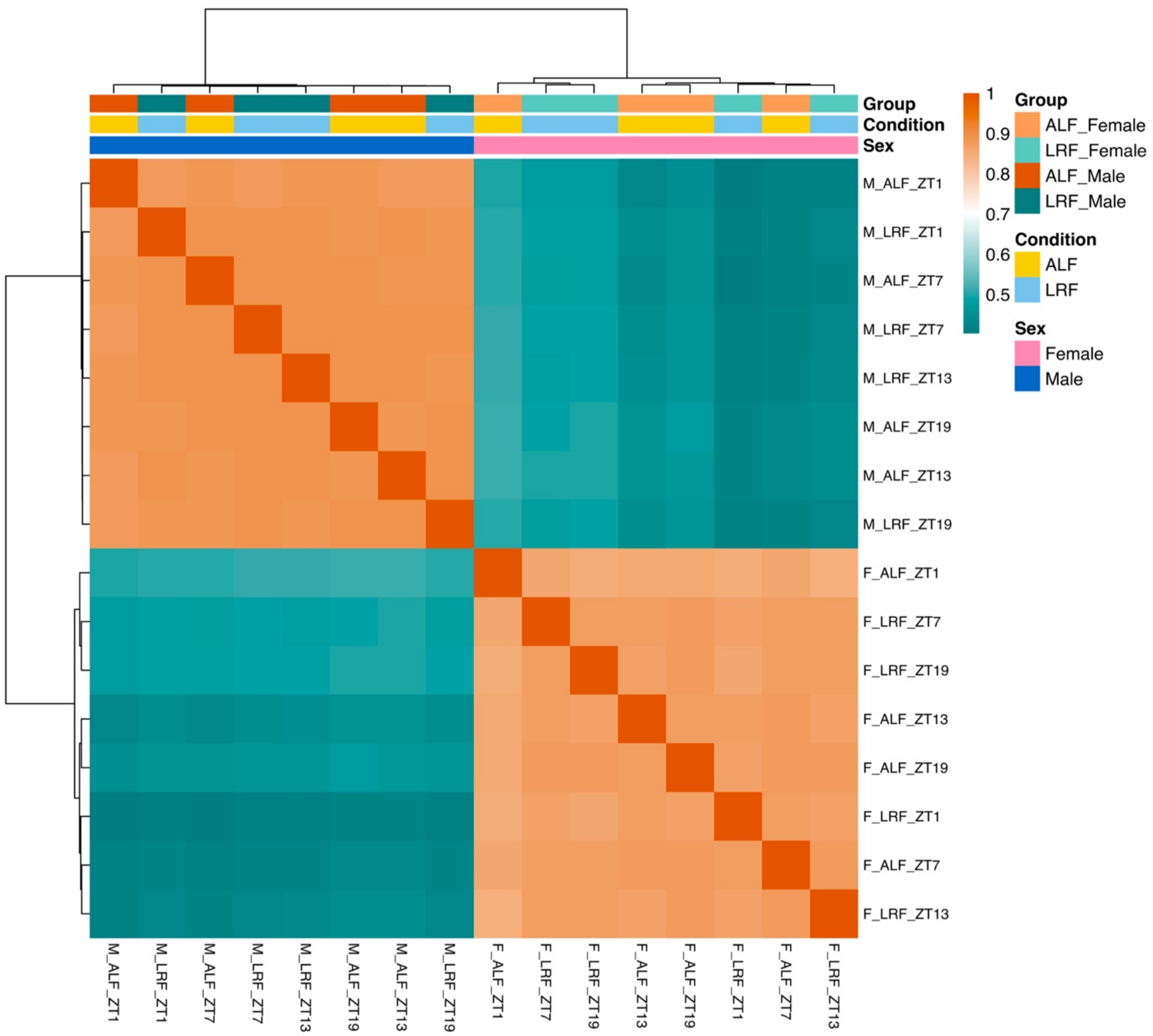

Group
Condition
Sex
1
0.9
0.8
0.7
0.6
0.5
Group
ALF_Female
LRF_Female
ALF_Male
LRF_Male
Condition
ALF
LRF
Sex
Female
Male
M_ALF_ZT1
M_LRF_ZT1
M_ALF_ZT7
M_LRF_ZT7
M_LRF_ZT13
M_ALF_ZT19
M_ALF_ZT13
M_LRF_ZT19
F_ALF_ZT1
F_LRF_ZT7
F_LRF_ZT19
F_ALF_ZT13
F_ALF_ZT19
F_LRF_ZT1
F_ALF_ZT7
F_LRF_ZT13

### Classification Breakdown: SQANTI3 vs gffcompare

SQANTI3 classification (n = 305,324)

| SQANTI3 category | n | % |
|---|---|---|
| Full-Splice Match (FSM) | 278,363 | 91.17% |
| Novel Not in Catalog (NNC) | 23,362 | 7.65% |
| Novel In Catalog (NIC) | 2,804 | 0.92% |
| antisense | 350 | 0.11% |
| fusion | 168 | 0.06% |
| intergenic | 133 | 0.04% |
| genic_intron | 59 | 0.02% |
| Incomplete-Splice Match (ISM) | 54 | 0.02% |
| genic | 31 | 0.01% |

gffcompare classification (n = 305,324)

| gffcompare class_code | n | % |
|---|---|---|
| = (exact match) | 276,294 | 90.49% |
| j (novel junction combo) | 23,002 | 7.53% |
| k (containment) | 3,646 | 1.19% |
| u (intergenic/unclassified) | 1,470 | 0.48% |
| x (opposite strand exonic overlap) | 357 | 0.12% |
| n (partial intron retention) | 135 | 0.04% |
| c (contained) | 133 | 0.04% |
| m (full intron retention) | 119 | 0.04% |
| o (other same-strand overlap) | 77 | 0.03% |
| i (within ref. intron) | 77 | 0.03% |
| p (polymerase run-on) | 13 | <0.01% |
| y (contains ref. intron) | 1 | <0.01% |

#### SQANTI3 quality-control attributes by structural category

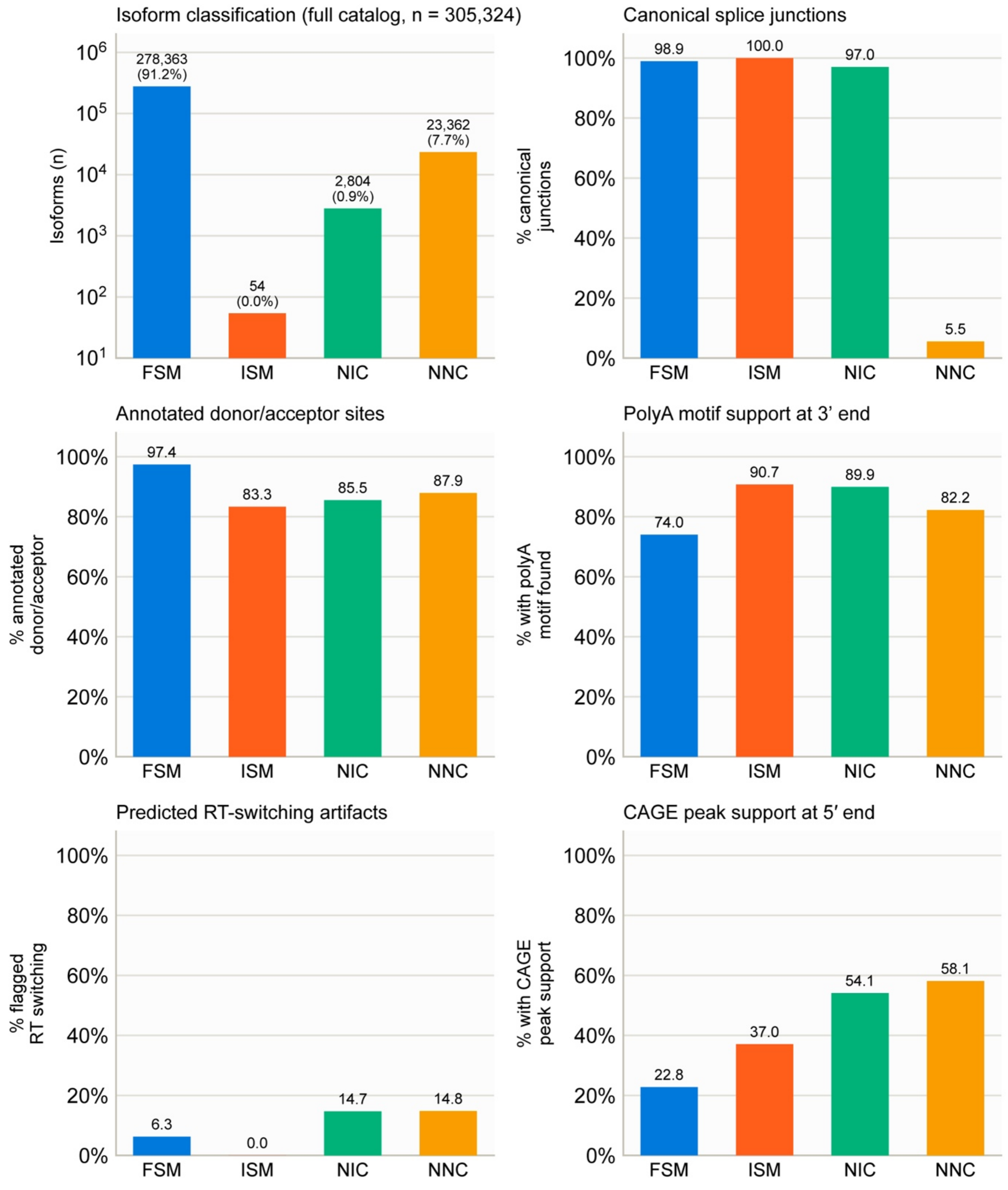

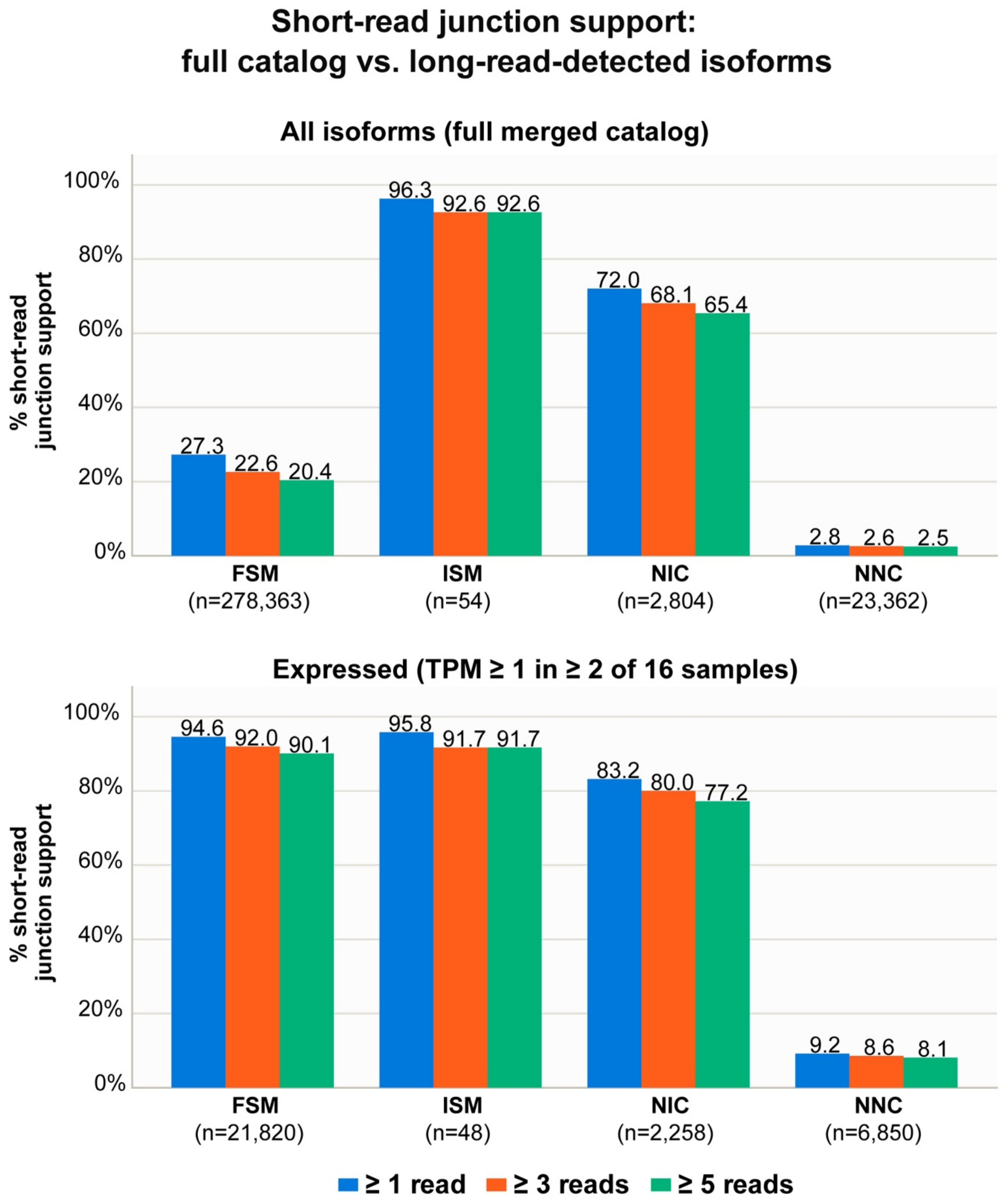
Short-read junction support:
full catalog vs. long-read-detected isoforms
All isoforms (full merged catalog)
% short-read junction support
100%
80%
60%
40%
20%
0%
27.3
22.6
20.4
96.3
92.6
92.6
72.0
68.1
65.4
2.8
2.6
2.5
FSM (n=278,363)
ISM (n=54)
NIC (n=2,804)
NNC (n=23,362)
Expressed (TPM ≥ 1 in ≥ 2 of 16 samples)
% short-read junction support
100%
80%
60%
40%
20%
0%
94.6
92.0
90.1
95.8
91.7
91.7
83.2
80.0
77.2
9.2
8.6
8.1
FSM (n=21,820)
ISM (n=48)
NIC (n=2,258)
NNC (n=6,850)
≥ 1 read
≥ 3 reads
≥ 5 reads

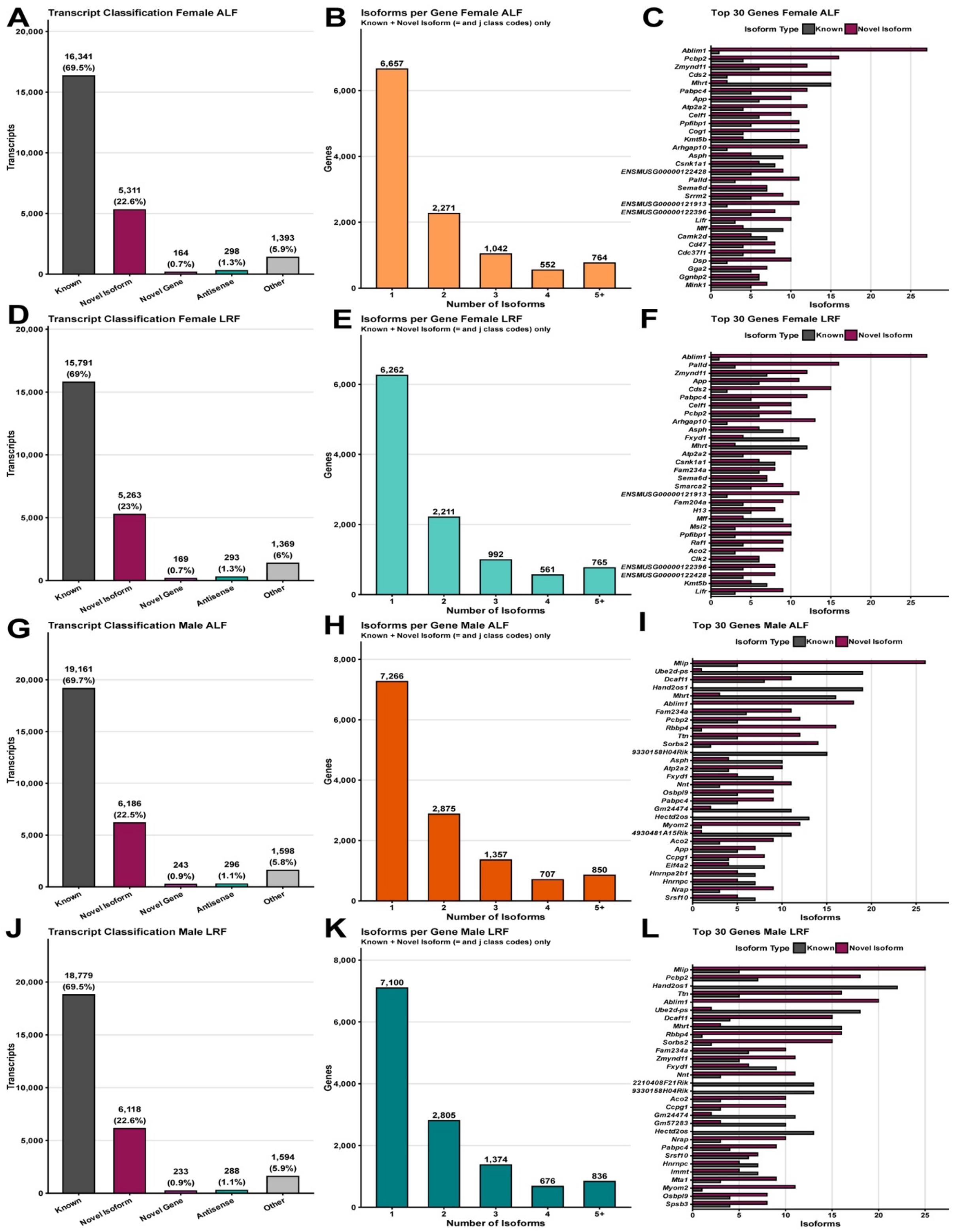

A
Transcript Classification Female ALF
Transcripts
16,341 (69.5%)
5,311 (22.6%)
164 (0.7%)
298 (1.3%)
1,393 (5.9%)
Known
Novel Isoform
Novel Gene
Antisense
Other
B
Isoforms per Gene Female ALF
Known + Novel Isoform (= and j class codes) only
Genes
6,657
2,271
1,042
552
764
Number of Isoforms
C
Top 30 Genes Female ALF
Isoform Type
Known
Novel Isoform
Ablim1
Pcbp2
Zmynd11
Cds2
Mhrt
Pabpc4
App
Atp2a2
Celf1
Ppfibp1
Cog1
Kmt5b
Arhgap10
Asph
Csnk1a1
ENSMUSG00000122428
Palld
Sema6d
Srrm2
ENSMUSG00000121913
ENSMUSG00000122396
Lifr
Mff
Camk2d
Cd47
Cdc37l1
Dsp
Gga2
Gnbp2
Mink1
Isoforms
D
Transcript Classification Female LRF
Transcripts
15,791 (69%)
5,263 (23%)
169 (0.7%)
293 (1.3%)
1,369 (6%)
Known
Novel Isoform
Novel Gene
Antisense
Other
E
Isoforms per Gene Female LRF
Known + Novel Isoform (= and j class codes) only
Genes
6,262
2,211
992
561
765
Number of Isoforms
F
Top 30 Genes Female LRF
Isoform Type
Known
Novel Isoform
Ablim1
Palld
Zmynd11
App
Cds2
Pabpc4
Celf1
Pcbp2
Arhgap10
Asph
Fxyd1
Mhrt
Atp2a2
Csnk1a1
Fam234a
Sema6d
Smarca2
ENSMUSG00000121913
Fam204a
H13
Mff
Msi2
Ppfibp1
Raf1
Aco2
Clk2
ENSMUSG00000122396
ENSMUSG00000122428
Kmt5b
Lifr
Isoforms
G
Transcript Classification Male ALF
Transcripts
19,161 (69.7%)
6,186 (22.5%)
243 (0.9%)
296 (1.1%)
1,598 (5.8%)
Known
Novel Isoform
Novel Gene
Antisense
Other
H
Isoforms per Gene Male ALF
Known + Novel Isoform (= and j class codes) only
Genes
7,266
2,875
1,357
707
850
Number of Isoforms
I
Top 30 Genes Male ALF
Isoform Type
Known
Novel Isoform
Mlip
Ube2d-ps
Dcaf11
Hand2os1
Mhrt
Ablim1
Fam234a
Pcbp2
Rbbp4
Ttn
Sorbs2
9330158H04Rik
Asph
Atp2a2
Fxyd1
Nnt
Osbpl9
Pabpc4
Gm24474
Hectd2os
Myom2
4930481A15Rik
Aco2
App
Ccpg1
Eif4a2
Hnrnpa2b1
Hnrnpc
Nrap
Srsf10
Isoforms
J
Transcript Classification Male LRF
Transcripts
18,779 (69.5%)
6,118 (22.6%)
233 (0.9%)
288 (1.1%)
1,594 (5.9%)
Known
Novel Isoform
Novel Gene
Antisense
Other
K
Isoforms per Gene Male LRF
Known + Novel Isoform (= and j class codes) only
Genes
7,100
2,805
1,374
676
836
Number of Isoforms
L
Top 30 Genes Male LRF
Isoform Type
Known
Novel Isoform
Mlip
Pcbp2
Hand2os1
Ttn
Ablim1
Ube2d-ps
Dcaf11
Mhrt
Rbbp4
Sorbs2
Fam234a
Zmynd11
Fxyd1
Nnt
2210408F21Rik
9330158H04Rik
Aco2
Ccpg1
Gm24474
Gm57283
Hectd2os
Nrap
Pabpc4
Srsf10
Hnrnpc
Immt
Mta1
Myom2
Osbpl9
Spsb3
Isoforms

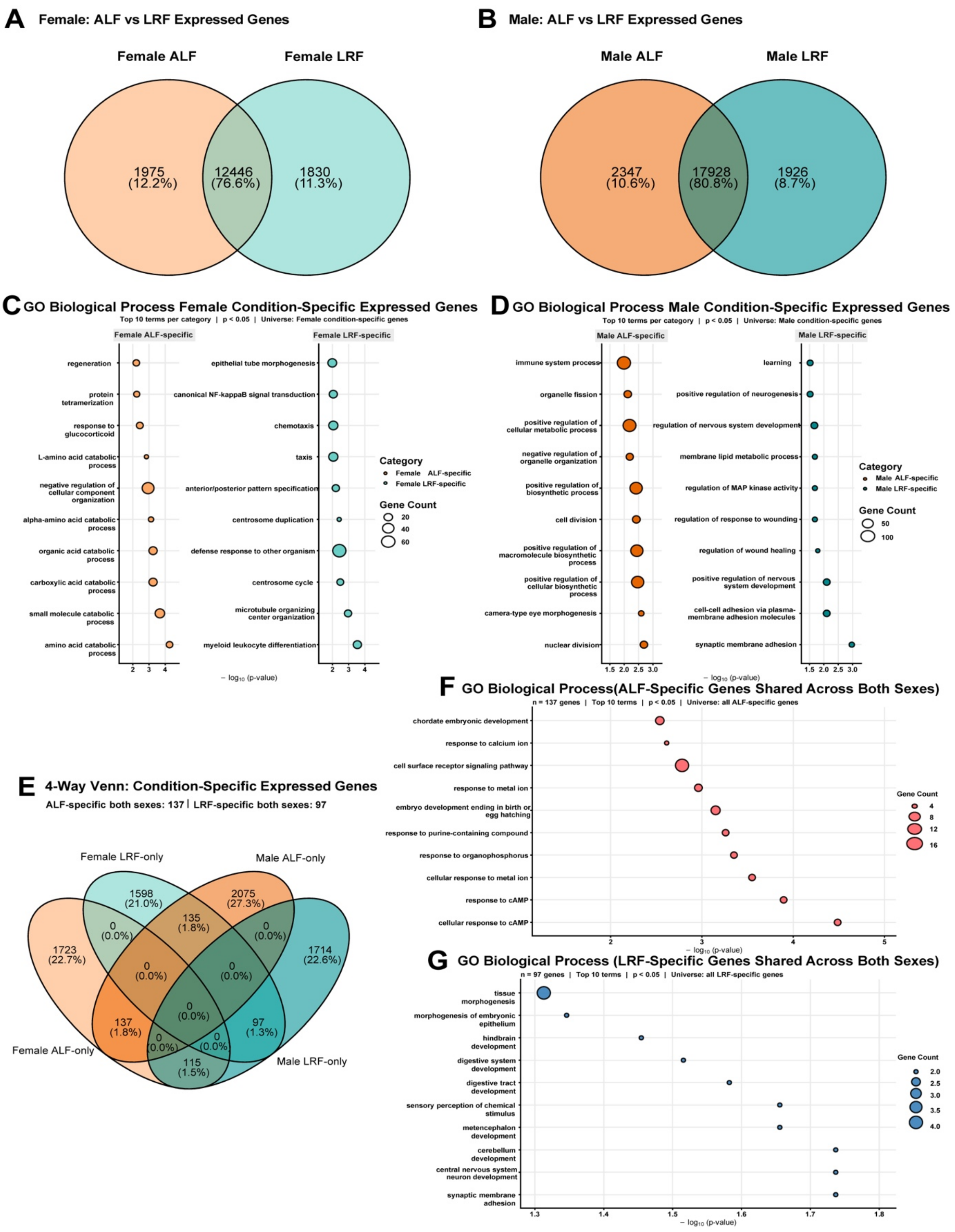
A Female: ALF vs LRF Expressed Genes
Female ALF
Female LRF
1975 (12.2%)
12446 (76.6%)
1830 (11.3%)
B Male: ALF vs LRF Expressed Genes
Male ALF
Male LRF
2347 (10.6%)
17928 (80.8%)
1926 (8.7%)
C GO Biological Process Female Condition-Specific Expressed Genes
Top 10 terms per category | p < 0.05 | Universe: Female condition-specific genes
Female ALF-specific
regeneration
protein tetramerization
response to glucocorticoid
L-amino acid catabolic process
negative regulation of cellular component organization
alpha-amino acid catabolic process
organic acid catabolic process
carboxylic acid catabolic process
small molecule catabolic process
amino acid catabolic process
Female LRF-specific
epithelial tube morphogenesis
canonical NF-kappaB signal transduction
chemotaxis
taxis
anterior/posterior pattern specification
centrosome duplication
defense response to other organism
centrosome cycle
microtubule organizing center organization
myeloid leukocyte differentiation
− log10 (p-value)
Category
Female ALF-specific
Female LRF-specific
Gene Count
20
40
60
D GO Biological Process Male Condition-Specific Expressed Genes
Top 10 terms per category | p < 0.05 | Universe: Male condition-specific genes
Male ALF-specific
immune system process
organelle fission
positive regulation of cellular metabolic process
negative regulation of organelle organization
positive regulation of biosynthetic process
cell division
positive regulation of macromolecule biosynthetic process
positive regulation of cellular biosynthetic process
camera-type eye morphogenesis
nuclear division
Male LRF-specific
learning
positive regulation of neurogenesis
regulation of nervous system development
membrane lipid metabolic process
regulation of MAP kinase activity
regulation of response to wounding
regulation of wound healing
positive regulation of nervous system development
cell-cell adhesion via plasma-membrane adhesion molecules
synaptic membrane adhesion
− log10 (p-value)
Category
Male ALF-specific
Male LRF-specific
Gene Count
50
100
E 4-Way Venn: Condition-Specific Expressed Genes
ALF-specific both sexes: 137 | LRF-specific both sexes: 97
Female LRF-only
Male ALF-only
Female ALF-only
Male LRF-only
1598 (21.0%)
2075 (27.3%)
135 (1.8%)
1723 (22.7%)
1714 (22.6%)
0 (0.0%)
137 (1.8%)
97 (1.3%)
115 (1.5%)
F GO Biological Process(ALF-Specific Genes Shared Across Both Sexes)
n = 137 genes | Top 10 terms | p < 0.05 | Universe: all ALF-specific genes
chordate embryonic development
response to calcium ion
cell surface receptor signaling pathway
response to metal ion
embryo development ending in birth or egg hatching
response to purine-containing compound
response to organophosphorus
cellular response to metal ion
response to cAMP
cellular response to cAMP
− log10 (p-value)
Gene Count
4
8
12
16
G GO Biological Process (LRF-Specific Genes Shared Across Both Sexes)
n = 97 genes | Top 10 terms | p < 0.05 | Universe: all LRF-specific genes
tissue morphogenesis
morphogenesis of embryonic epithelium
hindbrain development
digestive system development
digestive tract development
sensory perception of chemical stimulus
metencephalon development
cerebellum development
central nervous system neuron development
synaptic membrane adhesion
− log10 (p-value)
Gene Count
2.0
2.5
3.0
3.5
4.0

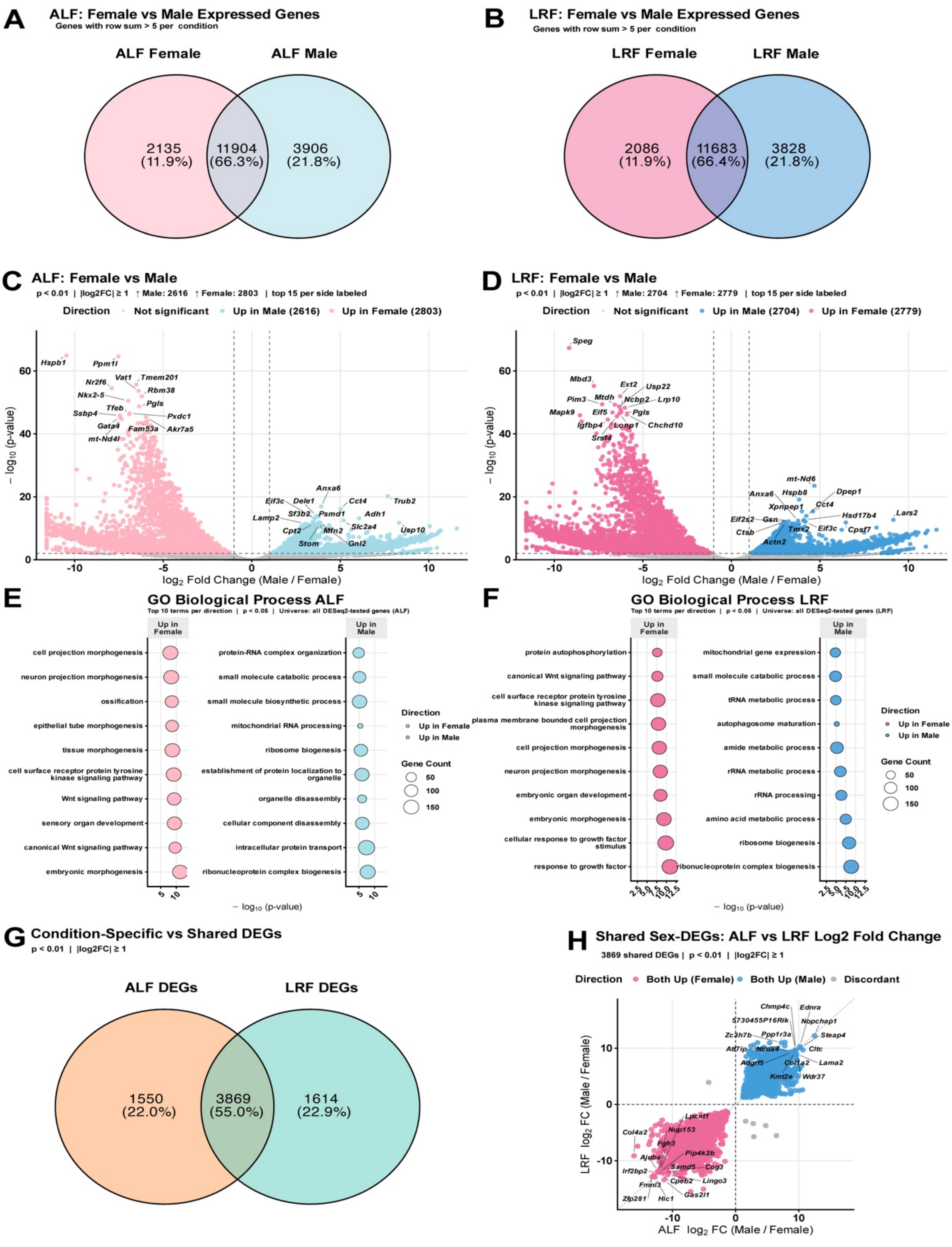
A
ALF: Female vs Male Expressed Genes
Genes with row sum > 5 per condition
ALF Female
ALF Male
2135 (11.9%)
11904 (66.3%)
3906 (21.8%)
B
LRF: Female vs Male Expressed Genes
Genes with row sum > 5 per condition
LRF Female
LRF Male
2086 (11.9%)
11683 (66.4%)
3828 (21.8%)
C
ALF: Female vs Male
p < 0.01 | |log2FC| ≥ 1 ↑ Male: 2616 ↑ Female: 2803 | top 15 per side labeled
Direction
Not significant
Up in Male (2616)
Up in Female (2803)
− log10 (p-value)
log2 Fold Change (Male / Female)
Hspb1
Ppm1l
Vat1
Tmem201
Nr2f6
Rbm38
Nkx2-5
Pgls
Tfeb
Ssbp4
Pxdc1
Gata4
Fam53a
Akr7a5
mt-Nd4l
Anxa6
Eif3c
Dele1
Cct4
Trub2
Sf3b2
Psmd1
Adh1
Lamp2
Slc2a4
Cpt2
Mfn2
Usp10
Stom
Gnl2
D
LRF: Female vs Male
p < 0.01 | |log2FC| ≥ 1 ↑ Male: 2704 ↑ Female: 2779 | top 15 per side labeled
Direction
Not significant
Up in Male (2704)
Up in Female (2779)
− log10 (p-value)
log2 Fold Change (Male / Female)
Speg
Mbd3
Ext2
Usp22
Mtdh
Pim3
Ncbp2
Lrp10
Mapk9
Eif5
Pgls
Igfbp4
Lonp1
Chchd10
Srsf4
mt-Nd6
Anxa6
Hspb8
Dpep1
Xpnpep1
Cct4
Eif2s2
Gsn
Hsd17b4
Lars2
Ctsb
Tmx2
Eif3c
Cpsf7
Actn2
E
GO Biological Process ALF
Top 10 terms per direction | p < 0.05 | Universe: all DESeq2-tested genes (ALF)
Up in Female
cell projection morphogenesis
neuron projection morphogenesis
ossification
epithelial tube morphogenesis
tissue morphogenesis
cell surface receptor protein tyrosine kinase signaling pathway
Wnt signaling pathway
sensory organ development
canonical Wnt signaling pathway
embryonic morphogenesis
Up in Male
protein-RNA complex organization
small molecule catabolic process
small molecule biosynthetic process
mitochondrial RNA processing
ribosome biogenesis
establishment of protein localization to organelle
organelle disassembly
cellular component disassembly
intracellular protein transport
ribonucleoprotein complex biogenesis
Direction
Up in Female
Up in Male
Gene Count
50
100
150
− log10 (p-value)
F
GO Biological Process LRF
Top 10 terms per direction | p < 0.05 | Universe: all DESeq2-tested genes (LRF)
Up in Female
protein autophosphorylation
canonical Wnt signaling pathway
cell surface receptor protein tyrosine kinase signaling pathway
plasma membrane bounded cell projection morphogenesis
cell projection morphogenesis
neuron projection morphogenesis
embryonic organ development
embryonic morphogenesis
cellular response to growth factor stimulus
response to growth factor
Up in Male
mitochondrial gene expression
small molecule catabolic process
tRNA metabolic process
autophagosome maturation
amide metabolic process
rRNA metabolic process
rRNA processing
amino acid metabolic process
ribosome biogenesis
ribonucleoprotein complex biogenesis
Direction
Up in Female
Up in Male
Gene Count
50
100
150
− log10 (p-value)
G
Condition-Specific vs Shared DEGs
p < 0.01 | |log2FC| ≥ 1
ALF DEGs
LRF DEGs
1550 (22.0%)
3869 (55.0%)
1614 (22.9%)
H
Shared Sex-DEGs: ALF vs LRF Log2 Fold Change
3869 shared DEGs | p < 0.01 | |log2FC| ≥ 1
Direction
Both Up (Female)
Both Up (Male)
Discordant
LRF log2 FC (Male / Female)
ALF log2 FC (Male / Female)
Chmp4c
Ednra
5730455P16Rik
Nopchap1
Zc3h7b
Ppp1r3a
Steap4
Atf7ip
Ncoa4
Cltc
Adgrf5
Col1a2
Lama2
Kmt2e
Wdr37
Lpcat1
Col4a2
Nup153
Fgfr3
Pip4k2b
Ajuba
Samd5
Cog3
Irf2bp2
Cpeb2
Lingo3
Fmnl3
Gas2l1
Zfp281
Hic1

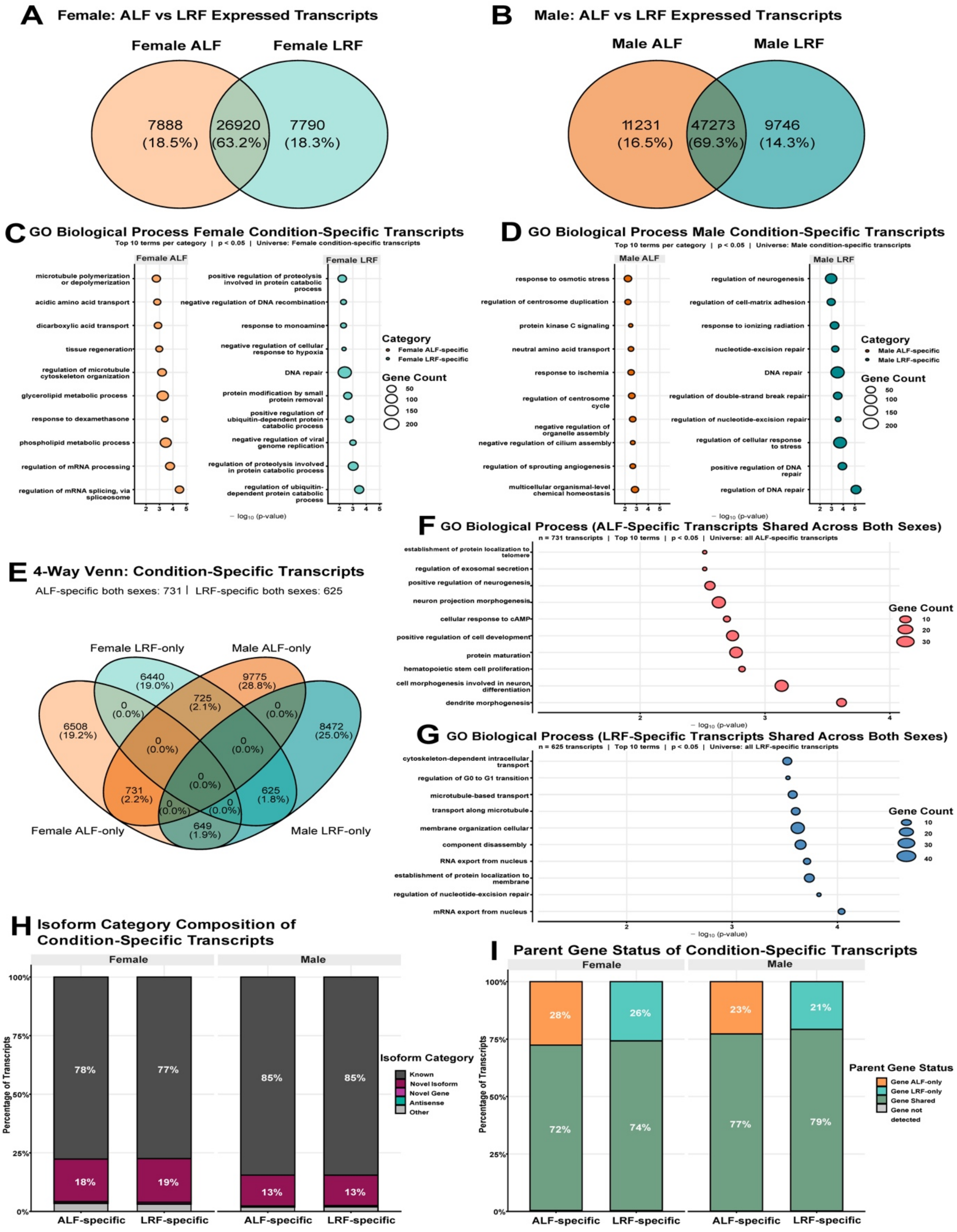
A Female: ALF vs LRF Expressed Transcripts
Female ALF
Female LRF
7888 (18.5%)
26920 (63.2%)
7790 (18.3%)
B Male: ALF vs LRF Expressed Transcripts
Male ALF
Male LRF
11231 (16.5%)
47273 (69.3%)
9746 (14.3%)
C GO Biological Process Female Condition-Specific Transcripts
Top 10 terms per category | p < 0.05 | Universe: Female condition-specific transcripts
D GO Biological Process Male Condition-Specific Transcripts
Top 10 terms per category | p < 0.05 | Universe: Male condition-specific transcripts
E 4-Way Venn: Condition-Specific Transcripts
ALF-specific both sexes: 731 | LRF-specific both sexes: 625
Female LRF-only
Male ALF-only
Female ALF-only
Male LRF-only
F GO Biological Process (ALF-Specific Transcripts Shared Across Both Sexes)
n = 731 transcripts | Top 10 terms | p < 0.05 | Universe: all ALF-specific transcripts
G GO Biological Process (LRF-Specific Transcripts Shared Across Both Sexes)
n = 625 transcripts | Top 10 terms | p < 0.05 | Universe: all LRF-specific transcripts
H Isoform Category Composition of Condition-Specific Transcripts
Isoform Category
Known
Novel Isoform
Novel Gene
Antisense
Other
I Parent Gene Status of Condition-Specific Transcripts
Parent Gene Status
Gene ALF-only
Gene LRF-only
Gene Shared
Gene not detected

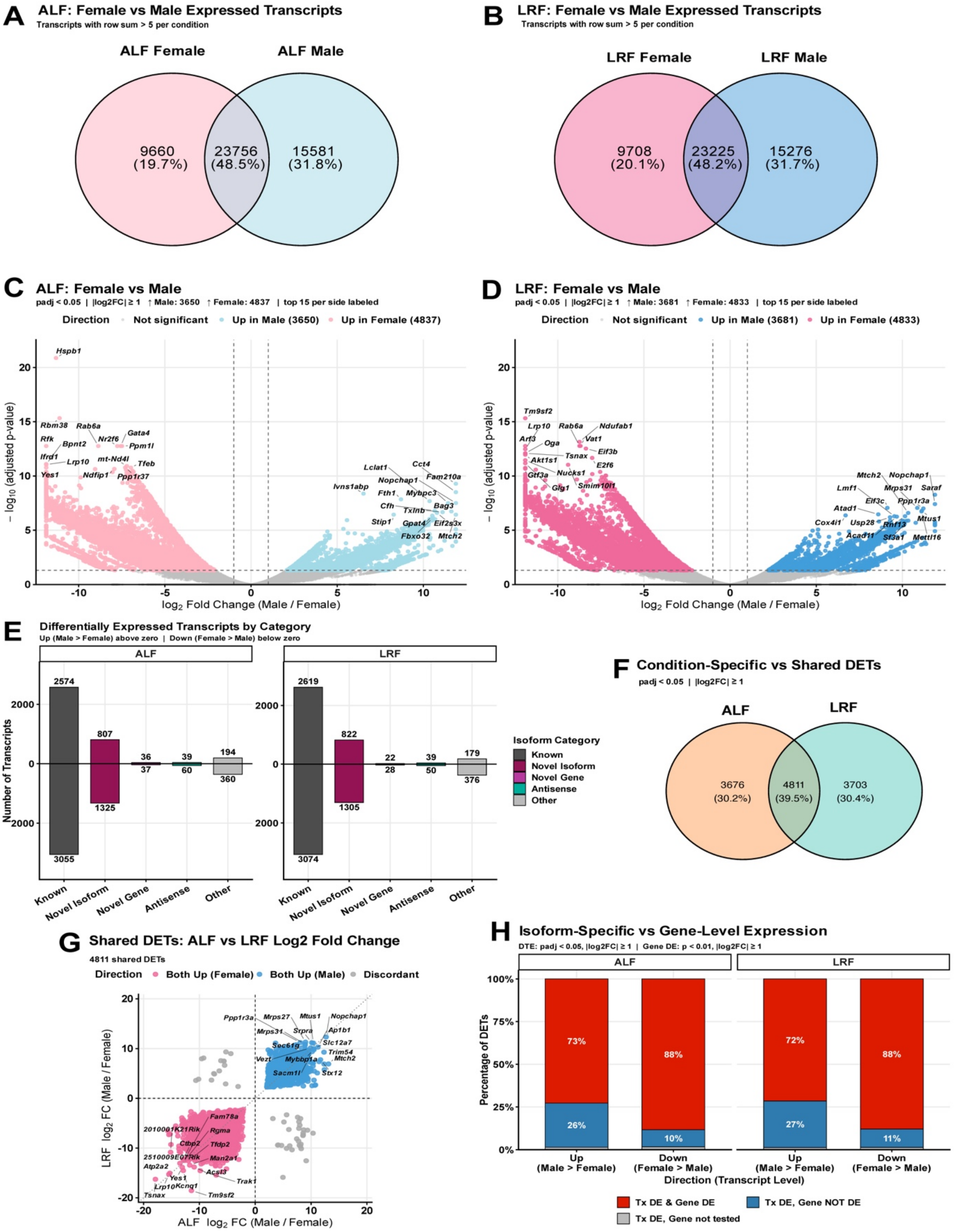

A
ALF: Female vs Male Expressed Transcripts
Transcripts with row sum > 5 per condition
ALF Female
ALF Male
9660 (19.7%)
23756 (48.5%)
15581 (31.8%)
B
LRF: Female vs Male Expressed Transcripts
Transcripts with row sum > 5 per condition
LRF Female
LRF Male
9708 (20.1%)
23225 (48.2%)
15276 (31.7%)
C
ALF: Female vs Male
padj < 0.05 | |log2FC| ≥ 1 ↑ Male: 3650 ↑ Female: 4837 | top 15 per side labeled
Direction
Not significant
Up in Male (3650)
Up in Female (4837)
Hspb1
Rbm38
Rab6a
Rfk
Bpnt2
Nr2f6
Gata4
Ppm1l
Ifrd1
Lrp10
mt-Nd4l
Tfeb
Yes1
Ndfip1
Ppp1r37
Lclat1
Cct4
Fam210a
Ivns1abp
Nopchap1
Fth1
Mybpc3
Cfh
Txlnb
Bag3
Stip1
Gpat4
Eif2s3x
Fbxo32
Mtch2
– log10 (adjusted p-value)
log2 Fold Change (Male / Female)
D
LRF: Female vs Male
padj < 0.05 | |log2FC| ≥ 1 ↑ Male: 3681 ↑ Female: 4833 | top 15 per side labeled
Direction
Not significant
Up in Male (3681)
Up in Female (4833)
Tm9sf2
Lrp10
Rab6a
Ndufab1
Arf3
Oga
Vat1
Tsnax
Eif3b
Akt1s1
E2f6
Gtf3a
Nucks1
Glg1
Smim10l1
Mtch2
Nopchap1
Lmf1
Mrps31
Saraf
Eif3c
Ppp1r3a
Atad1
Cox4i1
Usp28
Mtus1
Rnf13
Acad11
Sf3a1
Mettl16
– log10 (adjusted p-value)
log2 Fold Change (Male / Female)
E
Differentially Expressed Transcripts by Category
Up (Male > Female) above zero | Down (Female > Male) below zero
ALF
LRF
2574
807
36
39
194
3055
1325
37
60
360
2619
822
22
39
179
3074
1305
28
50
376
Number of Transcripts
Known
Novel Isoform
Novel Gene
Antisense
Other
Isoform Category
Known
Novel Isoform
Novel Gene
Antisense
Other
F
Condition-Specific vs Shared DETs
padj < 0.05 | |log2FC| ≥ 1
ALF
LRF
3676 (30.2%)
4811 (39.5%)
3703 (30.4%)
G
Shared DETs: ALF vs LRF Log2 Fold Change
4811 shared DETs
Direction
Both Up (Female)
Both Up (Male)
Discordant
Ppp1r3a
Mrps27
Mtus1
Nopchap1
Mrps31
Srpra
Ap1b1
Sec61g
Slc12a7
Trim54
Vezt
Mybbp1a
Mtch2
Sacm1l
Stx12
Fam78a
2010001K21Rik
Rgma
Ctbp2
Tfdp2
2510009E07Rik
Man2a1
Atp2a2
Acsl3
Yes1
Trak1
Lrp10
Kcnq1
Tsnax
Tm9sf2
LRF log2 FC (Male / Female)
ALF log2 FC (Male / Female)
H
Isoform-Specific vs Gene-Level Expression
DTE: padj < 0.05, |log2FC| ≥ 1 | Gene DE: p < 0.01, |log2FC| ≥ 1
ALF
LRF
73%
26%
88%
10%
72%
27%
88%
11%
Percentage of DETs
Up (Male > Female)
Down (Female > Male)
Direction (Transcript Level)
Tx DE & Gene DE
Tx DE, Gene NOT DE
Tx DE, Gene not tested

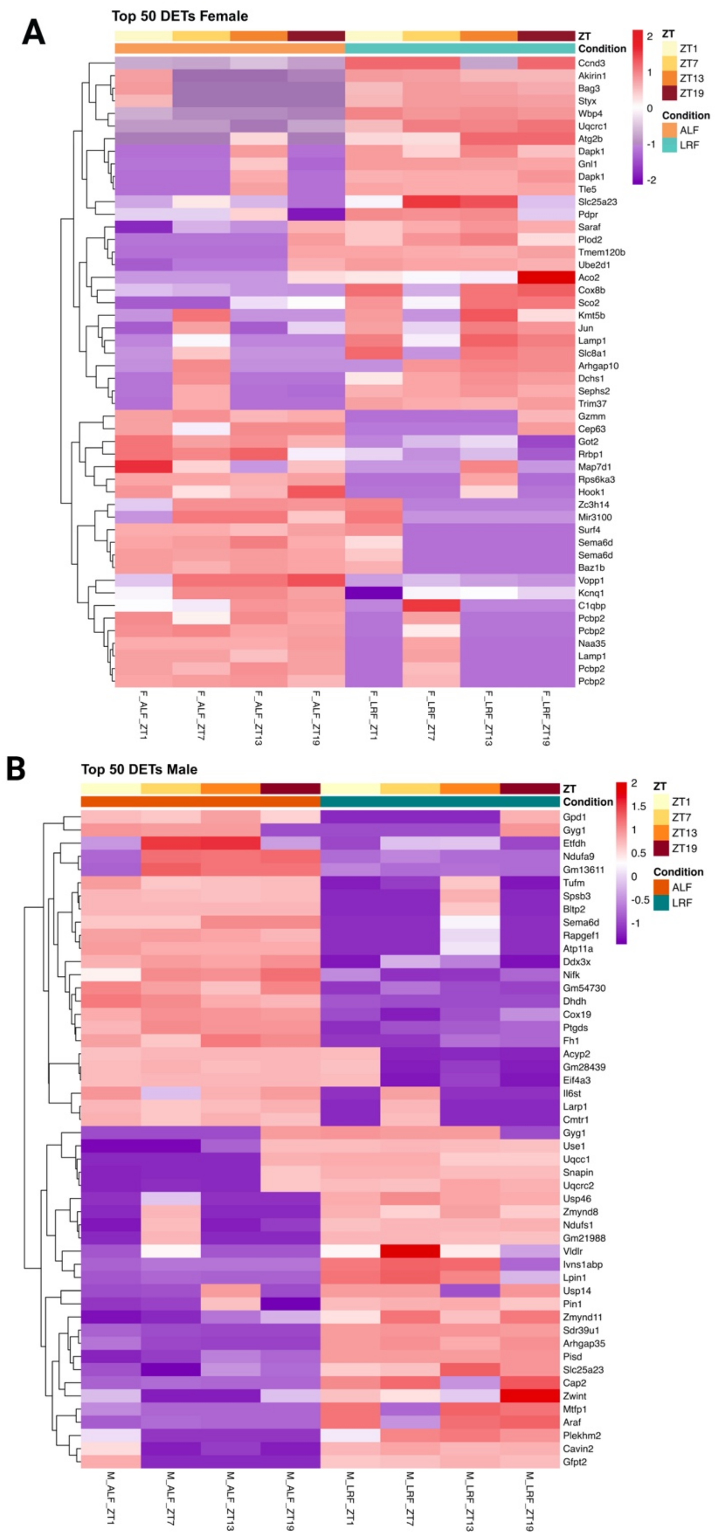
A
Top 50 DETs Female
ZT
Condition
Ccnd3
Akirin1
Bag3
Styx
Wbp4
Uqcrc1
Atg2b
Dapk1
Gnl1
Dapk1
Tle5
Slc25a23
Pdpr
Saraf
Plod2
Tmem120b
Ube2d1
Aco2
Cox8b
Sco2
Kmt5b
Jun
Lamp1
Slc8a1
Arhgap10
Dchs1
Sephs2
Trim37
Gzmm
Cep63
Got2
Rrbp1
Map7d1
Rps6ka3
Hook1
Zc3h14
Mir3100
Surf4
Sema6d
Sema6d
Baz1b
Vopp1
Kcnq1
C1qbp
Pcbp2
Pcbp2
Naa35
Lamp1
Pcbp2
Pcbp2
F_ALF_ZT1
F_ALF_ZT7
F_ALF_ZT13
F_ALF_ZT19
F_LRF_ZT1
F_LRF_ZT7
F_LRF_ZT13
F_LRF_ZT19
2
1
0
-1
-2
ZT
ZT1
ZT7
ZT13
ZT19
Condition
ALF
LRF
B
Top 50 DETs Male
ZT
Condition
Gpd1
Gyg1
Etfdh
Ndufa9
Gm13611
Tufm
Spsb3
Bltp2
Sema6d
Rapgef1
Atp11a
Ddx3x
Nifk
Gm54730
Dhdh
Cox19
Ptgds
Fh1
Acyp2
Gm28439
Eif4a3
Il6st
Larp1
Cmtr1
Gyg1
Use1
Uqcc1
Snapin
Uqcrc2
Usp46
Zmynd8
Ndufs1
Gm21988
Vldlr
Ivns1abp
Lpin1
Usp14
Pin1
Zmynd11
Sdr39u1
Arhgap35
Pisd
Slc25a23
Cap2
Zwint
Mtfp1
Araf
Plekhm2
Cavin2
Gfpt2
M_ALF_ZT1
M_ALF_ZT7
M_ALF_ZT13
M_ALF_ZT19
M_LRF_ZT1
M_LRF_ZT7
M_LRF_ZT13
M_LRF_ZT19
2
1.5
1
0.5
0
-0.5
-1
ZT
ZT1
ZT7
ZT13
ZT19
Condition
ALF
LRF

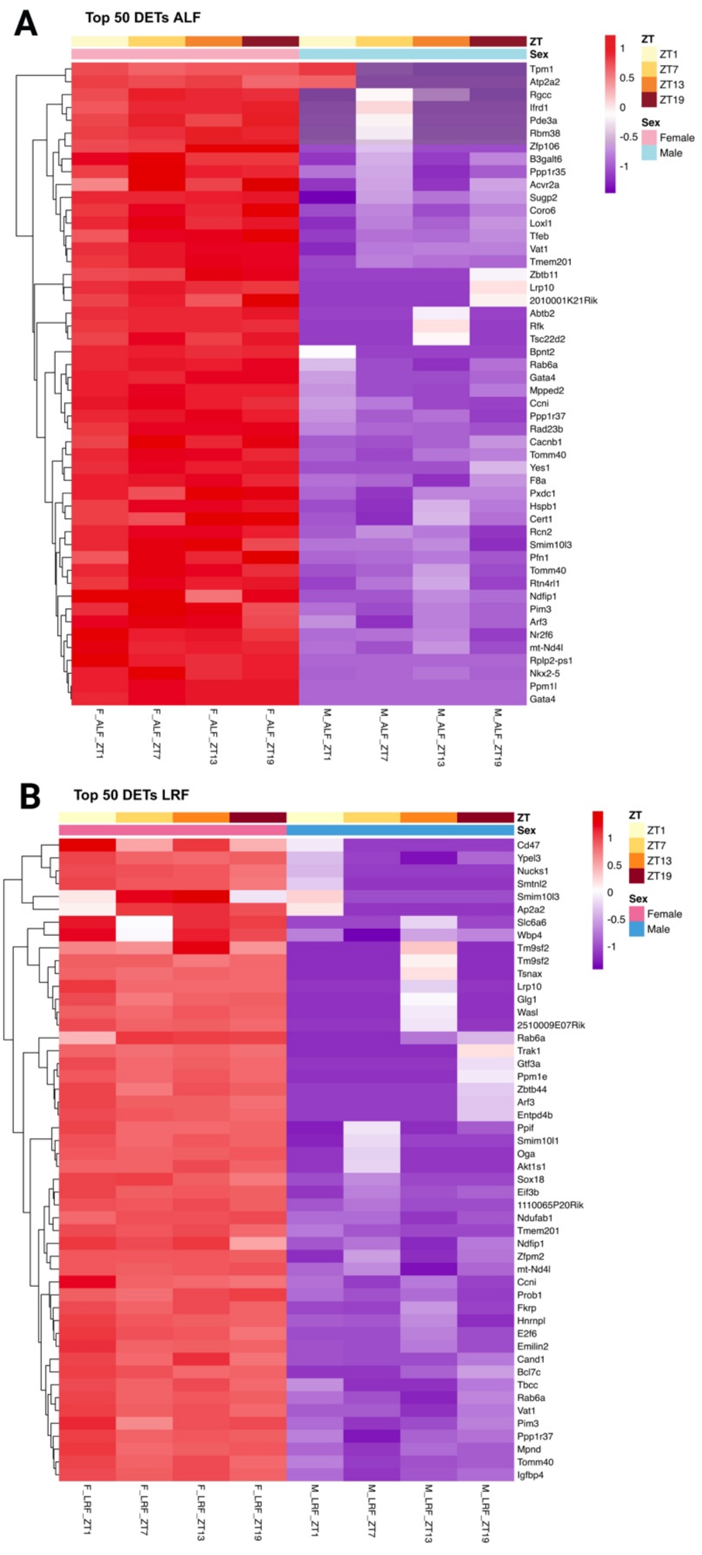
A
Top 50 DETs ALF
ZT
Sex
Tpm1
Atp2a2
Rgcc
Ifrd1
Pde3a
Rbm38
Zfp106
B3galt6
Ppp1r35
Acvr2a
Sugp2
Coro6
Loxl1
Tfeb
Vat1
Tmem201
Zbtb11
Lrp10
2010001K21Rik
Abtb2
Rfk
Tsc22d2
Bpnt2
Rab6a
Gata4
Mpped2
Ccni
Ppp1r37
Rad23b
Cacnb1
Tomm40
Yes1
F8a
Pxdc1
Hspb1
Cert1
Rcn2
Smim10l3
Pfn1
Tomm40
Rtn4rl1
Ndfip1
Pim3
Arf3
Nr2f6
mt-Nd4l
Rplp2-ps1
Nkx2-5
Ppm1l
Gata4
F_ALF_ZT1
F_ALF_ZT7
F_ALF_ZT13
F_ALF_ZT19
M_ALF_ZT1
M_ALF_ZT7
M_ALF_ZT13
M_ALF_ZT19
1
0.5
0
-0.5
-1
ZT
ZT1
ZT7
ZT13
ZT19
Sex
Female
Male
B
Top 50 DETs LRF
ZT
Sex
Cd47
Ypel3
Nucks1
Smtnl2
Smim10l3
Ap2a2
Slc6a6
Wbp4
Tm9sf2
Tm9sf2
Tsnax
Lrp10
Glg1
Wasl
2510009E07Rik
Rab6a
Trak1
Gtf3a
Ppm1e
Zbtb44
Arf3
Entpd4b
Ppif
Smim10l1
Oga
Akt1s1
Sox18
Eif3b
1110065P20Rik
Ndufab1
Tmem201
Ndfip1
Zfpm2
mt-Nd4l
Ccni
Prob1
Fkrp
Hnrnpl
E2f6
Emilin2
Cand1
Bcl7c
Tbcc
Rab6a
Vat1
Pim3
Ppp1r37
Mpnd
Tomm40
Igfbp4
F_LRF_ZT1
F_LRF_ZT7
F_LRF_ZT13
F_LRF_ZT19
M_LRF_ZT1
M_LRF_ZT7
M_LRF_ZT13
M_LRF_ZT19
1
0.5
0
-0.5
-1
ZT
ZT1
ZT7
ZT13
ZT19
Sex
Female
Male

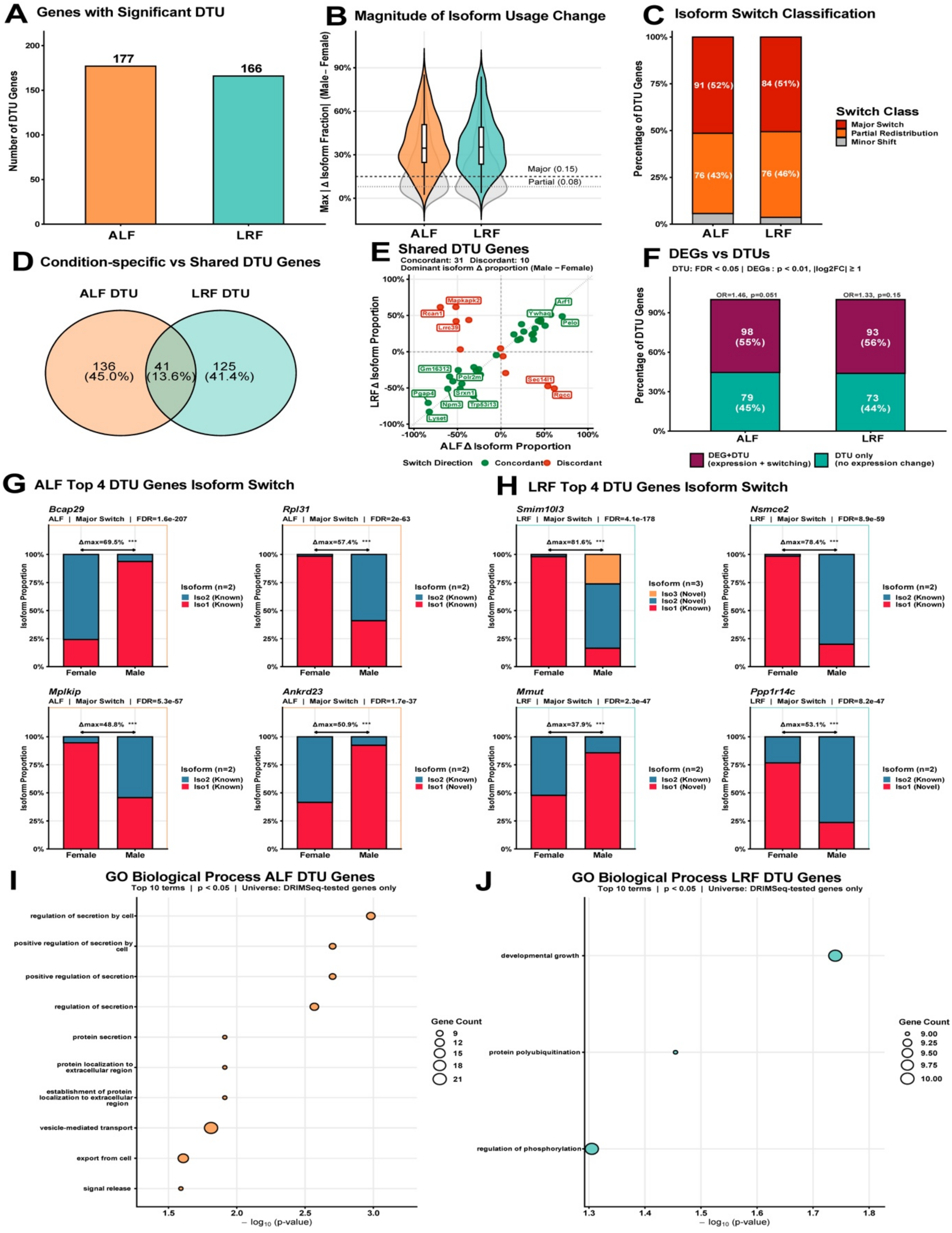
A Genes with Significant DTU
B Magnitude of Isoform Usage Change
C Isoform Switch Classification
D Condition-specific vs Shared DTU Genes
E Shared DTU Genes
F DEGs vs DTUs
G ALF Top 4 DTU Genes Isoform Switch
H LRF Top 4 DTU Genes Isoform Switch
I GO Biological Process ALF DTU Genes
J GO Biological Process LRF DTU Genes

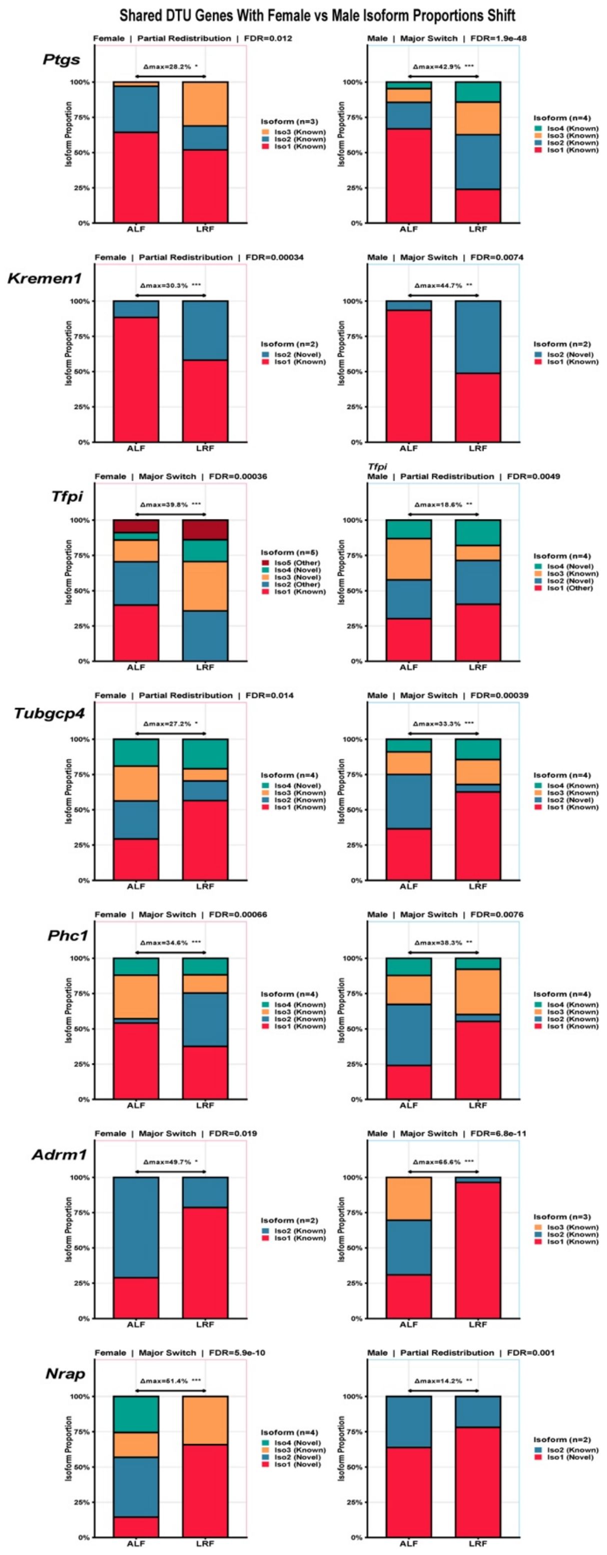
Shared DTU Genes With Female vs Male Isoform Proportions Shift
Ptgs
Female | Partial Redistribution | FDR=0.012
Δmax=28.2% *
Isoform (n=3)
Iso3 (Known)
Iso2 (Known)
Iso1 (Known)
Male | Major Switch | FDR=1.9e-48
Δmax=42.9% ***
Isoform (n=4)
Iso4 (Known)
Iso3 (Known)
Iso2 (Known)
Iso1 (Known)
Kremen1
Female | Partial Redistribution | FDR=0.00034
Δmax=30.3% ***
Isoform (n=2)
Iso2 (Novel)
Iso1 (Known)
Male | Major Switch | FDR=0.0074
Δmax=44.7% **
Isoform (n=2)
Iso2 (Novel)
Iso1 (Known)
Tfpi
Female | Major Switch | FDR=0.00036
Δmax=39.8% ***
Isoform (n=5)
Iso5 (Other)
Iso4 (Novel)
Iso3 (Novel)
Iso2 (Other)
Iso1 (Known)
Tfpi
Male | Partial Redistribution | FDR=0.0049
Δmax=18.6% **
Isoform (n=4)
Iso4 (Novel)
Iso3 (Known)
Iso2 (Novel)
Iso1 (Other)
Tubgcp4
Female | Partial Redistribution | FDR=0.014
Δmax=27.2% *
Isoform (n=4)
Iso4 (Novel)
Iso3 (Known)
Iso2 (Known)
Iso1 (Known)
Male | Major Switch | FDR=0.00039
Δmax=33.3% ***
Isoform (n=4)
Iso4 (Known)
Iso3 (Known)
Iso2 (Novel)
Iso1 (Known)
Phc1
Female | Major Switch | FDR=0.00066
Δmax=34.6% ***
Isoform (n=4)
Iso4 (Known)
Iso3 (Known)
Iso2 (Known)
Iso1 (Known)
Male | Major Switch | FDR=0.0076
Δmax=38.3% **
Isoform (n=4)
Iso4 (Known)
Iso3 (Known)
Iso2 (Known)
Iso1 (Known)
Adrm1
Female | Major Switch | FDR=0.019
Δmax=49.7% *
Isoform (n=2)
Iso2 (Known)
Iso1 (Known)
Male | Major Switch | FDR=6.8e-11
Δmax=65.6% ***
Isoform (n=3)
Iso3 (Known)
Iso2 (Known)
Iso1 (Known)
Nrap
Female | Major Switch | FDR=5.9e-10
Δmax=51.4% ***
Isoform (n=4)
Iso4 (Novel)
Iso3 (Known)
Iso2 (Novel)
Iso1 (Novel)
Male | Partial Redistribution | FDR=0.001
Δmax=14.2% **
Isoform (n=2)
Iso2 (Known)
Iso1 (Novel)
Isoform Proportion
0%
25%
50%
75%
100%
ALF
LRF

#### Female Cardiac DTU Genes: Isoform Proportions (ALF vs LRF)

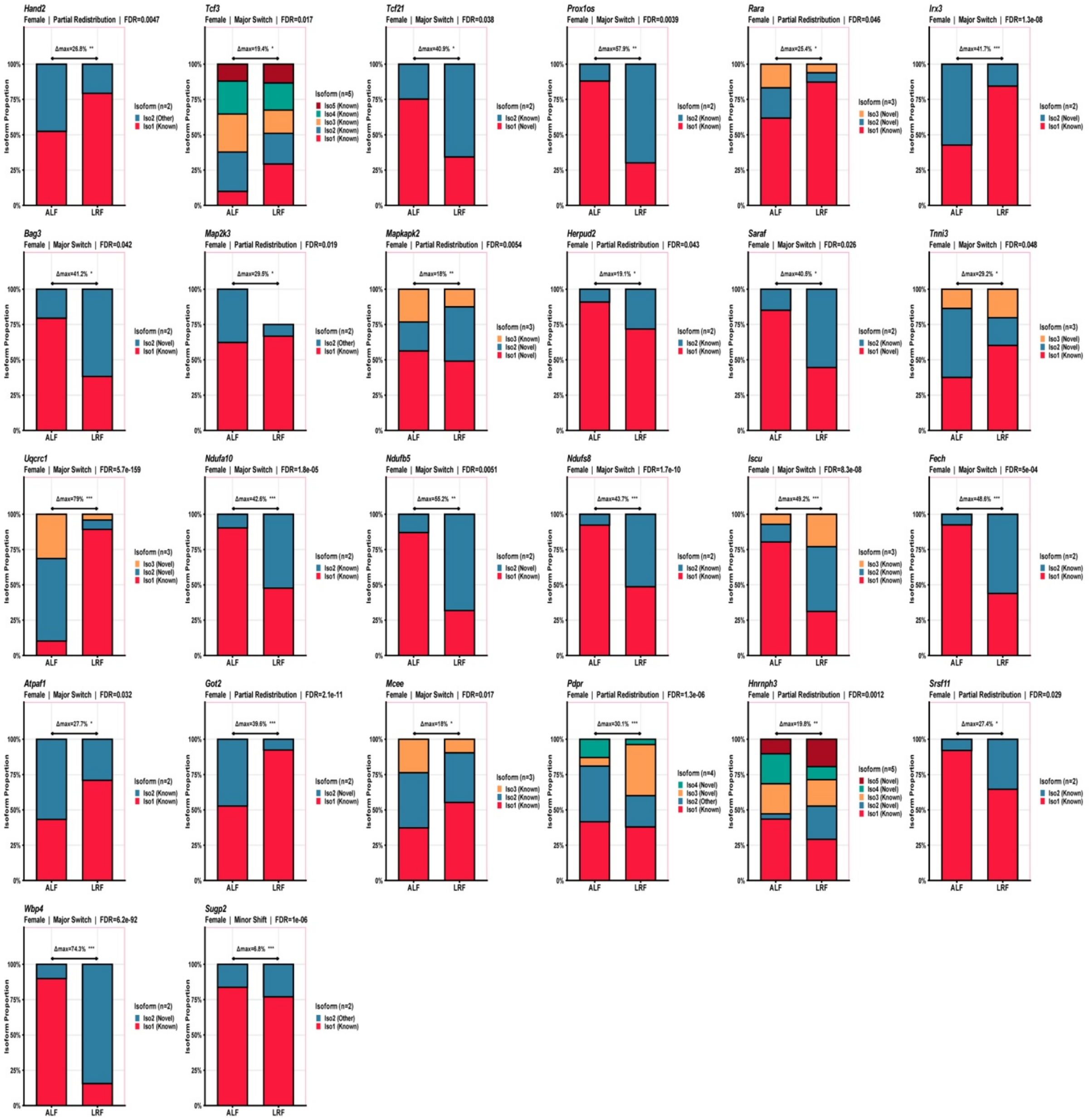

#### Male Cardiac DTU Genes: Isoform Proportions (ALF vs LRF)

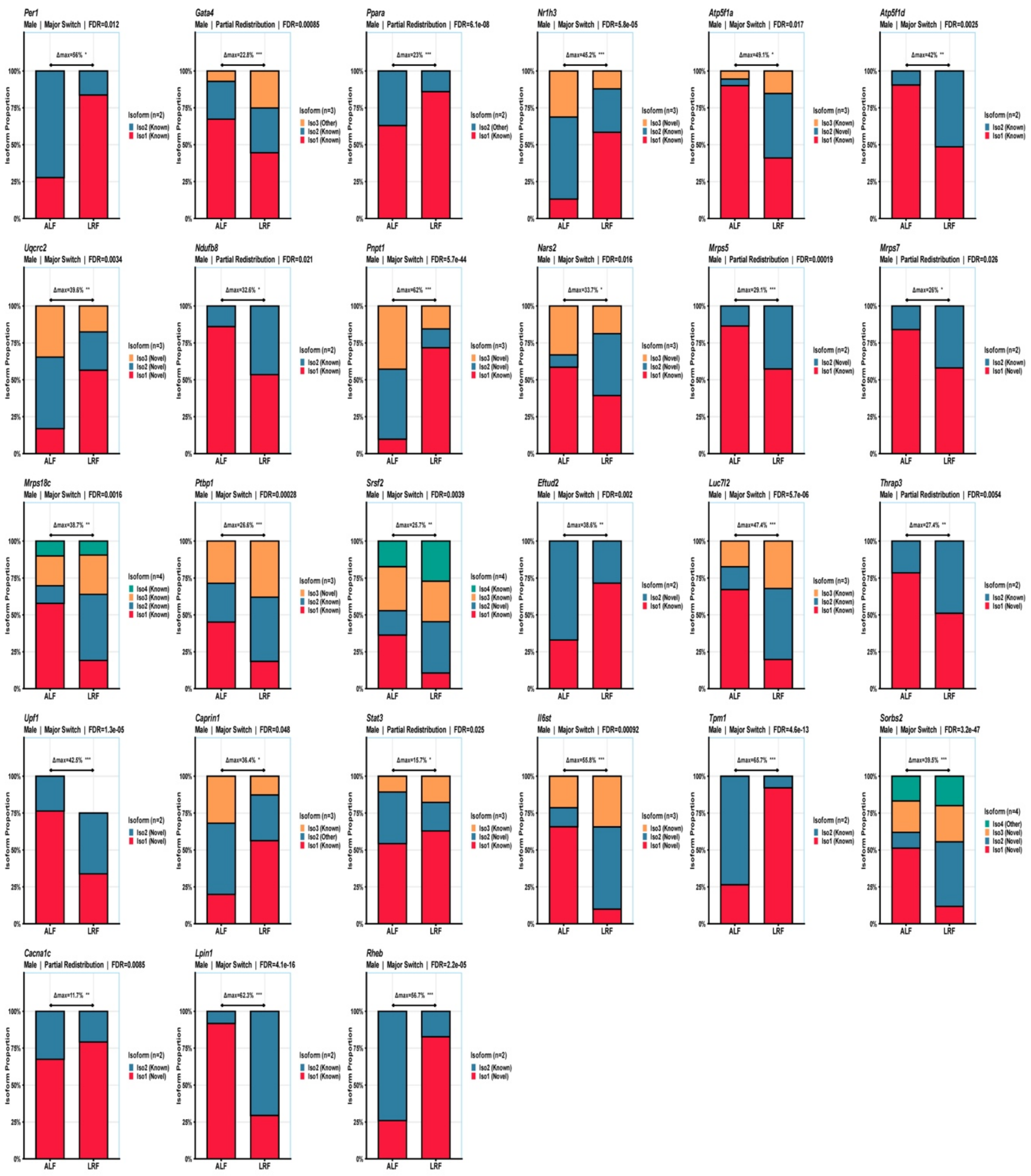
